# Methylomic alteration in peripheral blood lymphocytes of prodromal stage and first-episode Chinese Han schizophrenia patients

**DOI:** 10.1101/2021.02.07.430160

**Authors:** Yi Liu, Bing Lang, Robert C. Smith, Guodong Wang, Jijun Wang, Yong Liu, Hua Jin, Hailong Lyu, John M. Davis, Alessandro R. Guidotti, Jingping Zhao, Renrong Wu

## Abstract

**Background:** Although epigenetic dysregulation has long been proposed to promote the onset of schizophrenia, the landscape of the methylomic changes across the whole genome is yet established.

**Methods:** Using Infinium Human Methylation 850 BeadChip Array and MethylTarget sequencing method, we investigated the genome-wide methylation profiles and further validated methylation profiles of target genes in peripheral blood lymphocytes between individuals with psychosis risk syndrome (PRS), patients with first-episode schizophrenia (FES) and healthy controls (HC) in Chinese Han population.

**Results:** We detected 372 sites between psychosis risk syndrome (PRS) and healthy controls (HC), which increased to 460 sites in first-episode schizophrenia (FES) with 207 sites shared. Both PRS and FES featured profound hypomethylation within gene body. Gene ontology and network annotation merged on loci enriched in disease associated signaling pathways (MAPK(Mitogen Activated Protein Kinases), Glutamatergic, GABAergic etc.).

**Conclusions:** Our study implicated characteristic hypomethylation in both the discovery and validation cohorts in *SYNGAP1*, one of the frequently studied genes in neurodevelopmental disorders. This is the first methylome-wide association study between PRS and FES in Chinese Han population. Our findings provide potential biomarkers that can be used for future development of disease therapy and management.

## Introduction

Schizophrenia is a severe, chronic and socially disabling mental disorder with a life time risk around 1%. Despite strenuous research efforts, the precise etiology of schizophrenia still remains unclear. Both genetic vulnerability and environmental factors are involved in the disease progress. Although the heritability is high (60-80%) ^1^, current common and rare genetic variants discovered by GWAS ^2^ or CNVs ^3^ studies only explains a very small percentage of the total heritability. This suggests that other biological mechanisms contribute the heritability of schizophrenia.

DNA methylation involves addition of extra methyl group on the cytosine of CpG dinucleotides without changing DNA sequence. It is a major epigenetic mechanism which effectively regulates gene expression profile during development and also in response to environmental challenges ^4^, thus actively reshaping the landscape of gene transcription during organogenesis and tissue homeostasis. These changes are often retained after mitosis and some can be inherited across generations ^5^. Epigenetic changes are prominent in cancer ^6^ and have been reported in depression and schizophrenia ^7,8^.

Indeed, a growing number of studies has described aberrant methylomic signatures presented by a range of schizophrenia candidate genes or relevant genetic loci. In monozygotic twins discordant for schizophrenia, considerable disease-associated DNA methylation at specific loci have been reported, which strongly highlights their roles of mediating the phenotypic differences ^9^. Apart from monozygotic twin studies, many candidate genes including *GAD1, RELN, COMT*, and *SOX10* also display site-specific CpG methylation near the gene promoter regions, suggesting potentially regulatory roles for gene transcription ^10–13^. Despite these advances, current hypotheses cannot fully address the discrepancy between epigenetic alteration and changing of gene expression. For example, decreased reelin protein and mRNA transcripts are frequently reported in schizophrenia patients and animal models ^14^. However, the epigenetic data aimed at explaining this phenomenon seem controversial ^15–17^. Similarly, the aberrant methylation profile of catechol-O-methyltransferase gene (*COMT*) in schizophrenia ^18,19^ is inconsistent with fully quantitative methylation interrogation ^20^. A potential explanation is that previous studies mostly investigated gene promoters or upstream regulatory regions due to the design of microarray or the choice of DNA sequencing. As such, there is much less information about the large number of CpGs sparsely located in broad genome region. Notably, recent progress strongly suggests that gene body methylation also correlates with gene expression and may serve as a novel therapeutic target ^21^. For example, *DRD3* gene body hypermethylation significantly associates with the risk of schizophrenia ^22^. Similar association has been achieved in Rett syndrome ^23^, depression ^24^ and even neural tube defects ^25^. However, it is too early to draw any definitive conclusion about the influence of methylomic differences when the full methylomic picture across the whole genome remains uncertain in schizophrenia.

It is widely speculated that epigenetic alterations may vary during different phases of diseases. In schizophrenia, 73% patients experience prodromal stage (PRS) featuring attenuated psychotic symptoms prior to full conversion into psychosis. If methylomic changes could be reliably identified during the prodromal state it could potentially provide information to help in early diagnosis or prevention strategies.

In this study, we performed cross-sectional methylome-wide association studies using Illumina infinium methylation 850 BeadChip array and compared blood methylation patterns between PRS, first-episode schizophrenia patients (FES) and healthy controls (HC). Both PRS and FES groups presented distinct but overlapping methylomic signatures which were significantly differentiated from healthy controls. We also conducted an original exploration of abnormally-methylated CpG sites across the whole genome followed by cluster and pathway enrichment analysis. Top differentially methylated genes were further validated in separate cohort using MethylTarget sequencing method. Our study provides new insights to understand disease-phase associated methylation biomarkers in peripheral blood for schizophrenia.

## Materials and Methods

### Subjects

Our experimental paradigm consisted of two independent cohorts for genome-wide DNA methylation analysis (discovery cohort) and targeted bisulfite sequencing (validation cohort). In discovery cohort, forty psychosis risk syndrome (PRS) subjects, forty-eight patients with first episode schizophrenia (FES) and forty-seven age and sex matched healthy controls (HC) were recruited. In validation cohort, separate groups of PRS, FES and age, sex matched healthy controls with 41, 40 and 15 subjects were included respectively. All the subjects were recruited from the Department of Psychiatry at the Second Xiangya Hospital, Central South University, China and associated satellite partners at Guangzhou, Nanjing, Zhengzhou, Xinxiang, Ji’ning, Wuhan, Qingdao and Shanghai. They were subject to stringent admission pipeline management as per the following criteria: 1) Age range between 18-40 years in all groups; 2) Schizophrenia subjects met DSM-IV criteria for schizophrenia patients with first-episode onset; 3) PRS subjects met COPS criteria for psychosis risk syndrome. Healthy controls were collected from hospital staff or University undergraduates. They did not have documented family history of psychiatric problems or current diagnosis of psychotic illness including major depressive disorder, bipolar disorder, schizophrenia or other psychosis (please refer to the exclusion criteria below).

The exclusion criteria included: 1) subjects who met DSM-IV criteria for any psychotic disorder, or had delirium, dementia, amnesia, other severe cognitive impairment in the past or intellectual developmental disabilities (IQ<80) before meeting the COPS criteria for individuals with psychosis risk syndrome or DSM-IV criteria for schizophrenia diagnosis; 2) subjects with clinically significant somatic diseases; 3) subjects with substance abuse in the past 3 months; 4) subjects with a documented history of brain injury, epilepsy, or other known diseases of central nervous system; 5) subjects who received recent treatment with valproate or heavy smokers (> 1 pack/day, due to the potential effects of smoking on some epigenetic markers ^26–28^.

The diagnosis of schizophrenia or PRS syndrome was made by ratings of two independent psychiatrists using DSM-IV and COPS criteria, respectively, and accepted into the study if both agreed on the diagnosis. Written informed consent was obtained from all participants, and the study was approved by the second Xiangya Hospital Ethical Committee and review boards of all other participating institutions.

### General Psychiatric and Cognitive Assessment

The clinical symptoms of patients with schizophrenia were assessed using Positive and Negative Syndrome Scale (PANSS), prodromal symptoms of psychosis risk syndrome individuals were assessed using the Structured Interview for Psychosis-risk Syndromes (SIPS). We used Chinese version of MATRICS Consensus Cognitive battery (MCCB) to test the cognitive performance of all the subjects ^29^. The cognitive performance of all subjects was tested using Chinese version of MATRICS Consensus Cognitive battery (MCCB), including 7 cognitive domains (speed of processing, attention/vigilance, reason/problem solving, visual learning, spatial working memory, verbal learning, social cognition) and 9 test items of the original MCCB ^29^.

### Lymphocytes collection, Genomic DNA (gDNA) preparation and Genome-wide DNA methylation profiling

Ten milliliter venous blood was retrieved from each subject. Peripheral blood lymphocytes were harvested using Ficoll-Paque Plus mediated density gradient centrifuge immediately after the blood collection as reported previously ^30^. Samples were stored at −80°C freezer (Thermo) before gDNA extraction.

Genomic DNA was then extracted using QIAamp DNA Blood Mini Kit (Qiagen) and stored at −80°C freezer. All of the gDNA underwent strict quality control pipeline prior to bisulfate treatment which included Nanodrop quantification (Nanodrop 2000, Nanodrop technologies, Wilmington, DE, USA) and 1% agarose gel electrophoresis.

About 1μg of high-quality gDNA was bisulfite converted using the EZ DNA methylation kit (ZYMO, CA, USA) according to manufacturer’s instructions. Genome-wide DNA methylation profiles were generated with Illumina Infinium Methylation EPIC BeadChip Array (Illumina, Inc., San Diego, CA, USA). In detail, bisulfite-converted DNA was used for the whole genome amplification reaction, enzymatic fragmentation, precipitation and resuspension in hybridization buffer. Subsequent steps were carried out according to the standard Infinium HD Assay Methylation Protocol Guide. Bisulfited DNA was randomly assigned to Illumina Infinium Methylation EPIC BeadChip Array (Illumina, Inc., San Diego CA, USA) and methylomic profiling was read out by Illumina iScan System as per the manufacturers’ standard protocol (Illumina). The technical schemes, accuracy, and reproducibility of this microarray have been validated by previous report ^31^.

CpG dinucleotides sites spanning the whole genome including promoter, gene body, enhancer, and untranslated regions (UTR) were assessed. They also covered over 96% of CpG islands (CGIs) from the UCSC database with additional coverage in CGI shores (0-2 kb from CGI) and CGI shelves (2-4 kb from CGI). Detailed information on the contents of the array is available in the protocol of Infinium Methylation EPIC BeadChip Array Assay (www.illumina.com) and a recent publication ^31^.

### Data processing and filtering

For methylation analysis, raw Illumina IDAT files were run in the R environment and underwent background correction, probe type I/II correction and normalization. The probes with low-quality signals (*p*-value >0.05) or those overlapped with single nucleotide polymorphisms were discarded. We further removed the probes on the X and Y chromosomes, which left 805760 probes in total for analysis. Next, we calculated the methylation value-β value, which was calculated as the ratio of methylated signal intensity to the sum of the methylated and unmethylated signals after background subtraction. β value was used to represent methylation level of each CpG. Finally, each CpG site was related to a particular chromosome in the 37^th^ version genome build. Most of the CpG sites could match with a particular gene symbol in the UCSC database provided by Illumina, apart from those which were located in the intergenic regions.

### Analysis of genome wide methylation differences

Differentially methylated sites across the whole genome were detected using the limma package in the R environment. This procedure uses linear models to assess differential methylation, whereby information is shared across various sites. Sites were considered to be differentially methylated if the resulting adjusted *p*-value was < 0.05. The Benjamini–Hochberg method was used to adjust *p*-values and to ensure that the false discovery rate was <0.05.

### Pathway enrichment analysis

To investigate the biological relevance, pathway enrichment analysis was performed using the Kyoto Encyclopedia of Genes and Genomes (KEGG) human pathways and modules (www.genome.jp/kegg/download), along with gene function analysis using human Gene Ontology (GO) term associations defined in the GO biological process category (www.geneontology.org). KEGG is a database resource for understanding the high-level functions and utilizing the biological system, which integrates the information such as pathways, module, genome, compound, drug, disease, and enzyme. GO database is parallel to KEGG, which helps to understand the biological interactions such as protein-protein and protein-DNA interactions at the molecular, cellular component, and organism level.

### Targeted bisulfite sequencing of selected gene loci

We applied targeted bisulfite sequencing methods (MethylTarget™) to validate the selected differently methylated positions. Seventeen pairs of primers were designed by primer3 software (http://primer3.ut.ee/) with the bisulfite-converted control DNA as templates (see primers details in supplementary, Table S1). Following the quality control of the bisulfite primers, high-quality gDNA specimens were subject to sodium bisulfite conversion using EZ DNA methylation kit (ZYMO, CA, USA), followed by targeted amplification, loading, harvesting, pooling, barcoding and PCR amplification. The length distribution of the targeted libraries was validated and normalized by the Agilent 2100 Bioanalyzer. Finally, after the precise quantification of concentration, the targeted libraries were sequenced on the Illumina Hiseq/Miseq platform, which features a 150-base/250-base paired-end high-throughput sequencing run pattern (Hiseq/MiSeq, Illumina, Inc., San Diego, CA). All the aforementioned processes were performed as per manufacturer’s guidelines unless specified otherwise.

We used standard Illumina base-calling software to identify sequence readout from the targeted libraries sequencing. The sequence reads were then aligned and analyzed using a Zymo Research proprietary analysis procedure. Next, we removed poorer quality sequences and nucleotides during quality control analysis. The methylation level of each sampled cytosine was estimated as the number of reads reporting a C, divided by the total number of reads reporting a C or T.

### Correlation analysis

Correlation between these selected DMPs and cognitive performance was carried out in subjects with FES and PRS. Analyses were performed using SPSS 19 and 23. Ratings on the PANSS scale were performed based on PANSS total score and PANSS 5-factor scores as reported in previously international studies and Chinese adaption version ^32–34^. Multiple corrections were performed by using Benjamini-Hochberg method with the false discovery rate set at <0.05.

## Results

### Demographic, clinical and cognitive features of cohorts

After DNA quality control assessment, 40 subjects were finally included in each group of the discovery cohort whilst 41, 39, and 10 subjects were separately included in the PRS, FES and HC groups of the independent validation cohort. The demographic and clinical features of these subjects were collected and described in Supplementary, Table S2-a and S2-b.

No significant difference was found on the gender or age distribution between healthy controls and individuals with either FES or PRS. However, those with PRS or FES had significantly lower education level. Compared to healthy controls, they also displayed prevalently poor cognitive performance (Supplementary, Table S3-a and S3-b).

### Abnormal DNA methylome pattern in discovery cohort

#### Differential methylated sites analysis

In the discovery cohort, 372 differential methylation probes (DMPs) were revealed between PRS individuals and healthy controls which increased to 460 DMPs in FES patients (*p* value <0.05, Benjamini-Hochberg corrected). The intergroup methylation difference absolute value was over 0.08 based on Empirical Bayes method *t* test analysis. Strikingly, both the PRS and FES groups presented a global hypomethylation profile which correctly differentiated them from the healthy controls (Fig. 1; Supplementary, Table S4-a and S4-b). The probes and sample clustering pattern of these DMPs were illustrated in Fig. 1 (Heatmap, Fig. 1A, B; Volcano map, Fig. 1C, D).

**Figure 1.**
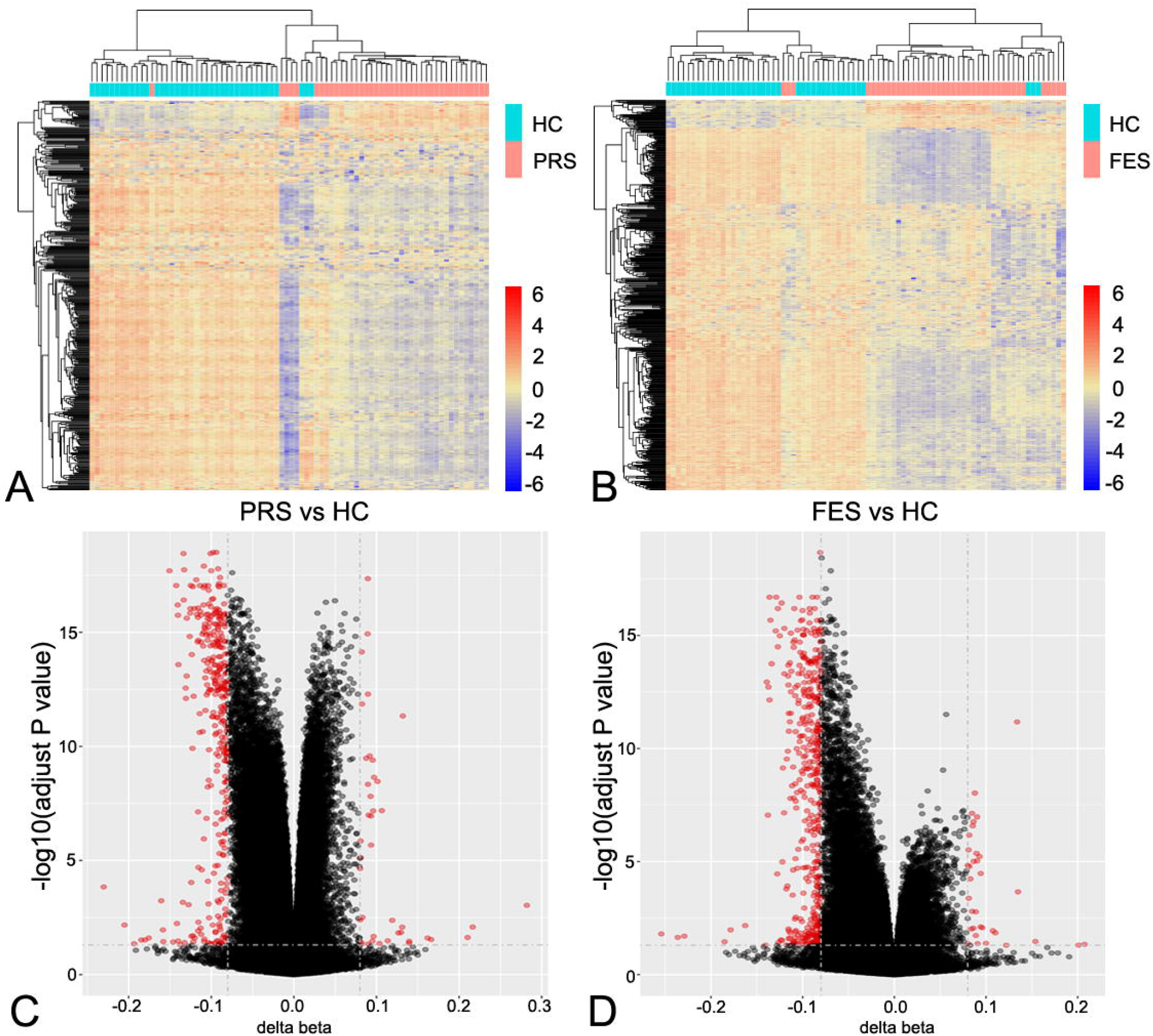
Comparation of DNA methylation profiles in individuals with PRS, FES and HC. **(A, B)** Both PRS (A) and FES (B) presented overall DNA hypomethylation landscapes as shown by heatmaps. X-row represents the clustered subjects and DMPs were aligned against the Y column. Blue and red colors represented DNA hypo- and hypermethylation signals respectively. **(C, D)** Similarly, volcano maps also showed a dominated DNA hypomethylation as opposed to hypermethylation in PRS (C) and FES (D) individuals. Y-column represents probes with or without significant detection (red or gray dots). Delta beta values were calculated and aligned against x-row showing the mean difference between PRS, FES and HC subjects. PRS, psychosis risk syndrome; FES, first-episode schizophrenia; HC, healthy controls.

Notably, 207 DMPs (Supplementary, Table S5-a) were shared by PRS and FES with the majority (198 DMPs) hypomethylated (Fig. 2A) and their distribution was shown in Fig. 2B, C. They all located in or within the proximity of 148 unique genes after being mapped against the whole genome (Supplementary, Table S5-a and S5-b).

**Figure 2.**
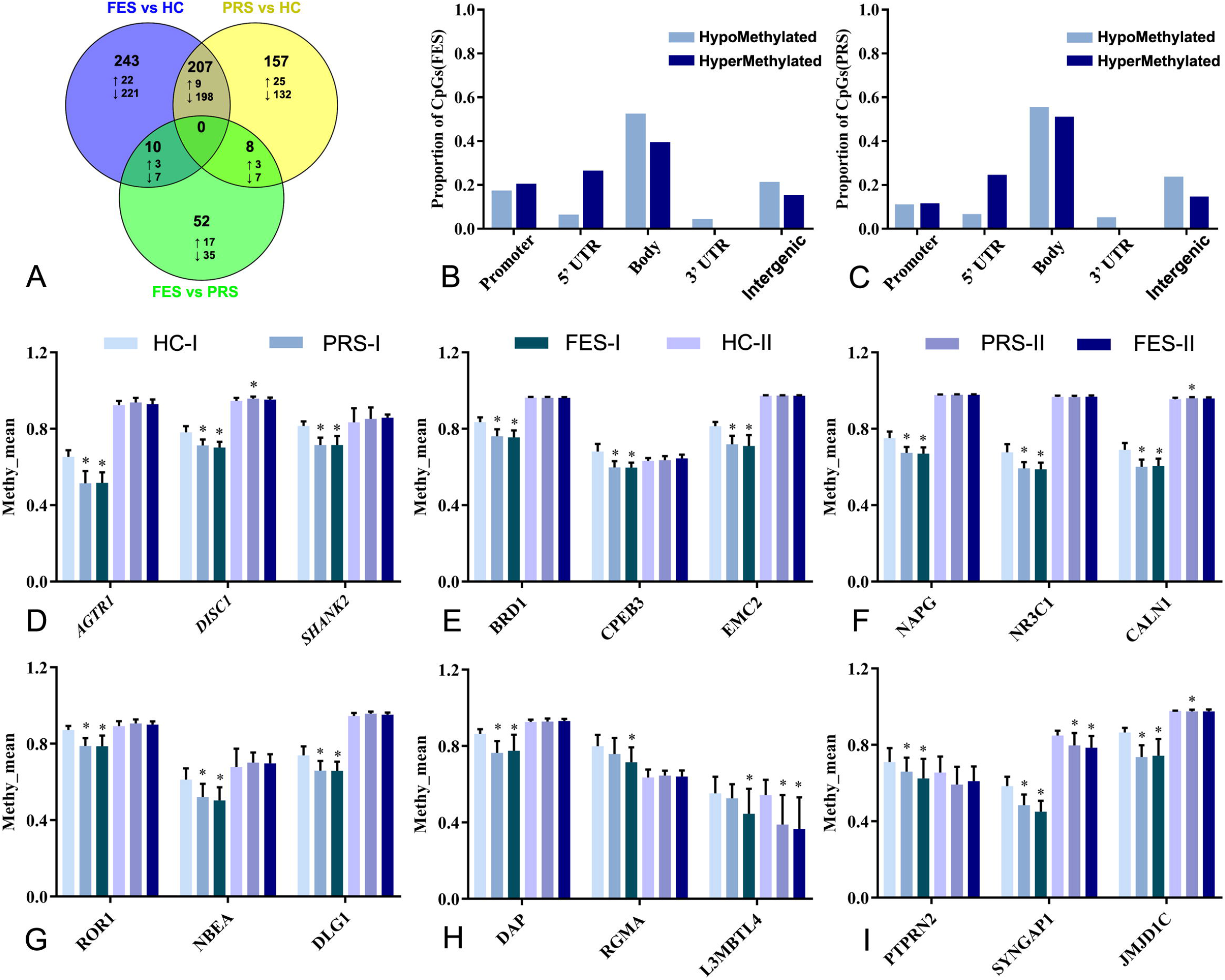
**(A)** PRS and FES individuals displayed distinct but overlapping DNA methylation profiles. Blue circle represented significant methylation changes between FES patients and HCs. Yellow circle showed significant methylation changes in PRS individuals and HCs. Green circle displayed significant changes between individuals with FES and PRS. 460 DMPs was detected between FES patients and HC group, 243 of these 460 DMPs only showed difference in FES and HC comparison. 372 DMPs was revealed between those with PRS and HC, 157 of these DMPs only showed difference in PRS and HC comparison. FES and PRS groups shared 207 DMPs, 198 of these DMPs showed hypomethylation in the group comparisons. Up arrow or down arrow represents probes which show hypermethylation or hypomethylation between group comparison. **(B, C)** Distribution of the differential methylated probes across the whole genome in FES (B) and PRS (C) individuals. The *x*-axis stands for the location of methylated CpG sites in different genomic regions. The *y* column represents the proportion of CpG sites with abnormal methylation. The majority of these hypomethylated DMPs were situated in promoter, gene body, 3’-UTR and intergenic regions. On the contrary, they were massively outnumbered by DMPs with hypermethylation in 5’UTR regions. **(D-I)** Eighteen genes were prioritized and subject to vigorous validation in independent cohort. Notably *L3MBTL4* (2H) showed robust hypomethylation in FES individuals in both the discovery and validation cohorts whilst *SYNGAP1* (2I) showed consistent difference in both PRS and FES individuals of these two cohorts. The error bars indicate standard deviation. The stars indicate significance between PRS or FES and HCs. I, discovery cohort; II, validation cohort; UTR, untranslated regions; PRS, psychosis risk syndrome; FES, patients with first-episode schizophrenia; HC, healthy controls. \**p*<0.05, Benjamini-Hochberg adjusted in discovery cohort.

### DNA methylome abnormalities were enriched in neuropsychiatric disease risk genes

Of the 148 genes, 128 had been previously reported and were associated with epigenetic regulation or neurodevelopmental disorders. The rest of 20 genes were newly reported and may potentially associate with schizophrenia biology. The top-ranked significant DMPs of the 128 genes (Supplementary, Table S5-b and S5-c) that show association with neuropsychiatric disease were enriched in the following genes: *HIST1H3I* (delta β= 0.085, *p* < 10^−17^), which was hypermethylated and located in the 1st Exon, *AGTR1* (delta β = −0.136, *p* < 10^−16^), which was hypermethylated and located 200 base pairs near the gene transcription starting site; *SHANK2* (delta β = −0.099,*p* < 10^−15^), *SYNGAP1* (delta β = −0.134, *p* < 10^−13^), *TSNARE1* (delta β = −0.092, *p* < 10^−15^), the DMPs preferentially resided within the gene body and were hypomethylated (Supplementary, Table S5-b). These results are in line with the current conception of polygenic origin of schizophrenia.

### Biological processes associated with DNA methylation abnormalities in discovery cohort

To fully understand the structured function and molecular interactions of the genes with abnormal methylation, the GO and KEGG database were applied to annotate precise function of the gene products and enrichment of signaling pathways. The top five ontology terms lied in the following annotation: (1) cytoplasm, cellular component; (2) nucleus, cellular component; (3) protein binding, molecular function; (4) cytosol, cellular component; (5) ATP binding, molecular function (FDR correction, adjusted *p* value <0.05; Supplementary, Table S6-a and S6-b). Similarly, the most relevant regulatory network revealed by KEGG analysis were: (1) MAPK signaling pathway, environmental information processing; (2) Pentose and glucuronate interconversions, carbohydrate metabolism; (3) Endocytosis in transport, cellular processes; (3) Glutamatergic/GABAergic synapse, neurotransmitters in nervous system; (4) ErbB, mTOR, PI3K-Akt signaling pathways, signal transduction and information processing; (5) Fc epsilon RI signaling pathway, immune systems (Supplementary, Table S7-a and S7-b).

### Relationship between psychosis-associated DMP and schizophrenia-associated gene SNP loci in discovery cohort

Out of these 460 DMPs, one DMP (cg00622907) was identified both in discovery and validation cohorts to locate approximately 2500 base pairs near one gene region reported in a 2014 Psychiatric Genomics Consortium mega-analysis: *TSNARE1*, region 5 ^2^ (Supplementary, Table S4 and S5). In region5 (chromosome8, 143309503-143330533), one schizophrenia-relevant SNP (rs4129585) is associated with methylation changes at this DMP (cg00622907) involved both in PRS and FES, which was located in a CpG island in body region of *TSNARE1*.

### DNA methylation abnormalities and haplotype analysis of target genes in discovery and validation cohorts

In order to replicate the DMPs in validation cohort, we prioritized them at a level of adjusted *p* <0.01 and selected the top genes based on but not limited to the following criteria: (1) association with psychiatric disorder, specially schizophrenia, (2) associated with neurodevelopment, (3) expression in the schizophrenia-associated brain regions, (4) support of animal model studies. Eighteen DMPs which were mapped to 18 genes were selected for validation by targeted bisulfite sequencing (Table 1).

**Table 1.** Association of eighteen DMPs between FES or PRS when compared to controls.

| Gene symbol | Probes | Chr | Gene context | Discovery cohort |  |  |  | Validation cohort |  |  |  |
| --- | --- | --- | --- | --- | --- | --- | --- | --- | --- | --- | --- |
| | | | | deltaβ(PRS-CON) | Adjusted $p$ (PRS-CON) | deltaβ(FES-CON) | Adjusted $p$ (FES-CON) | deltaβ(PRS-CON) | $p$ (PRS-CON) | deltaβ(FES-CON) | $p$ (FES-CON) |
| *AGTR1* | cg10297223 | 3 | TSS1500 | -0.1403447 | 1.80E-16 | -0.1362209 | 2.05E-17 | 0.0143093 | 0.0961220 | 0.0051993 | 0.5522000 |
| *DISC1* | cg25215021 | 1 | BODY | -0.0699056 | 6.34E-13 | -0.0805372 | 9.08E-16 | 0.0116174 | 0.0070989 | 0.0064331 | 0.1372040 |
| *SHANK2* | cg18440889 | 11 | BODY | -0.1000931 | 2.01E-18 | -0.0993903 | 9.45E-16 | 0.0182911 | 0.4126200 | 0.0240465 | 0.3380200 |
| *SYNGAP1* | cg14079545 | 6 | BODY | -0.1026416 | 1.41E-10 | -0.1338503 | 1.79E-14 | -0.0515072 | 0.0003050 | -0.0634096 | 1.80458E-05 |
| *L3MBTL4* | cg00451011 | 18 | 5'UTR | -0.0262376 | >0.05 | -0.1074379 | 0.00289839 | -0.1545190 | 0.0001421 | -0.1770100 | 3.8454E-05 |
| *BRD1* | cg19984503 | 22 | BODY | -0.0740183 | 9.96E-15 | -0.0802842 | 6.89E-14 | -0.0007316 | 0.6570800 | -0.0005139 | 0.7056600 |
| *CPEB3* | cg10967735 | 10 | BODY | -0.0846818 | 2.26E-13 | -0.0842451 | 1.02E-14 | 0.0033741 | 0.6398000 | 0.0127949 | 0.0695440 |
| *EMC2* | cg13442241 | 8 | BODY | -0.0960803 | 4.29E-16 | -0.1044153 | 4.92E-14 | 0.0006815 | 0.4841200 | -0.0005339 | 0.6294000 |
| *NAPG* | cg22389991 | 18 | BODY | -0.0784957 | 1.52E-13 | -0.0816163 | 1.94E-13 | 0.0006360 | 0.5763200 | 0.0001741 | 0.8767400 |
| *NR3C1* | cg19645279 | 5 | BODY | -0.085698 | 7.35E-13 | -0.0896682 | 4.54E-13 | -0.0008362 | 0.7095600 | 0.0006380 | 0.7958800 |
| *CALN1* | cg04158140 | 7 | BODY | -0.0904155 | 3.02E-14 | -0.0853187 | 6.02E-13 | 0.0050409 | 0.0217430 | 0.0041309 | 0.0731030 |
| *ROR1* | cg19963700 | 1 | BODY | -0.0832814 | 2.42E-15 | -0.0845379 | 7.09E-11 | 0.0142255 | 0.0658370 | 0.0074160 | 0.2810800 |
| *JMJD1C* | cg17983571 | 10 | BODY | -0.1297133 | 2.41E-16 | -0.1214894 | 7.25E-10 | -0.0028992 | 0.0495160 | -0.0024918 | 0.1264990 |
| *NBEA* | cg00500007 | 13 | BODY | -0.0947138 | 2.55E-07 | -0.1098392 | 1.54E-08 | 0.0230093 | 0.4801400 | 0.0181785 | 0.5737600 |
| *DLG1* | cg07068730 | 3 | BODY | -0.0804616 | 2.45E-08 | -0.0807601 | 2.32E-08 | -0.0010603 | 0.3013400 | -0.0005426 | 0.5993200 |
| *DAP* | cg03278573 | 5 | BODY | -0.0988018 | 5.03E-12 | -0.0882522 | 3.36E-06 | 0.0018058 | 0.7439400 | 0.0044883 | 0.2619400 |
| *RGMA* | cg17350690 | 15 | BODY | -0.0404528 | >0.05 | -0.0839641 | 0.0001222<br>6 | 0.0098382 | 0.4994000 | 0.0047293 | 0.6985600 |
| *PTPRN2* | cg21705926 | 7 | BODY | -0.0511071 | 0.0021957 | -0.0864364 | 0.0030875 | -0.0629056 | 0.0568180 | -0.0452916 | 0.1093800 |

DMP=Differential Methylated Probes, FES=First-episode schizophrenia, PRS=Psychosis risk syndrome, CON=Healthy controls, Chr=chromosome.
TSS=Transcription Start Site. Adjusted *p*: *p* value after Benjamini-Hochberg correction.

Although we were not able to replicate the down-regulated DNA methylation for the majority of these DMPs (Fig. 2D-I), one DMP cg14079545 in the gene *SYNGAP1*, stood out with consistent hypomethylation in both FES and PRS individuals after multiple correction (See Table 1 and Fig. 2I). Another DMP cg00451011 in the gene *L3MBTL4* also presented significant hypomethylation in FES patients in both cohorts (See Table 1 and Fig. 2H). These results potentially highlight a disease phase-specific methylomic changes and a collective effect may be implicated in PRS individuals and their transition towards FES. We also performed methylation specific haplotype analysis of individual cytosines and found significant association of the haplotypes of *SYNGAP1* and *L3MBTL4* gene (C and T) within individuals with FES and PRS (Supplementary, Table S8).

### Correlations between DNA methylation abnormalities and psychiatric, cognitive characteristics in discovery and validation cohorts

We then asked the possible association between domains of MCCB and abnormal DNA methylation. In the PRS group of discovery cohort, we detected significant correlations between MCCB performance and the methylation level of probes in *NR3C1, NBEA, CPEB3, DLG1, DAP, RGMA*, and *L3MBTL4* at an unadjusted *p* value <0.05 (Table 2-b). Similarly, in the FES patients, we also detected a strong association between MCCB performance and the methylation level of probes in *SHANK2, DLG1, CALN1*, and *JMJD1C* at an unadjusted *p* value <0.05 (Table 2-a). However, none of these correlations survived corrected Benjamini-Hochberg (BH) significance at α=.05.

**Table 2-a.**
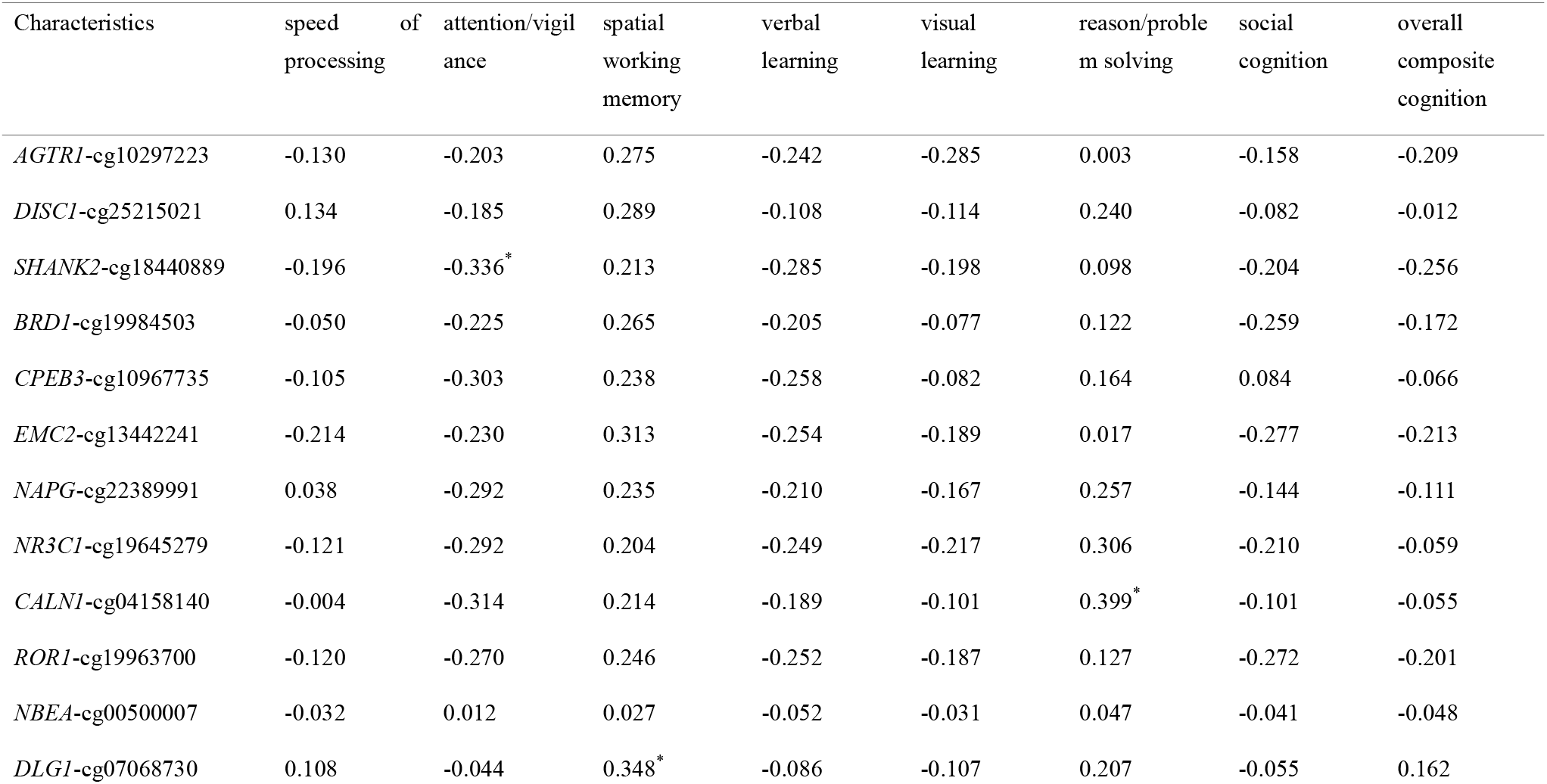
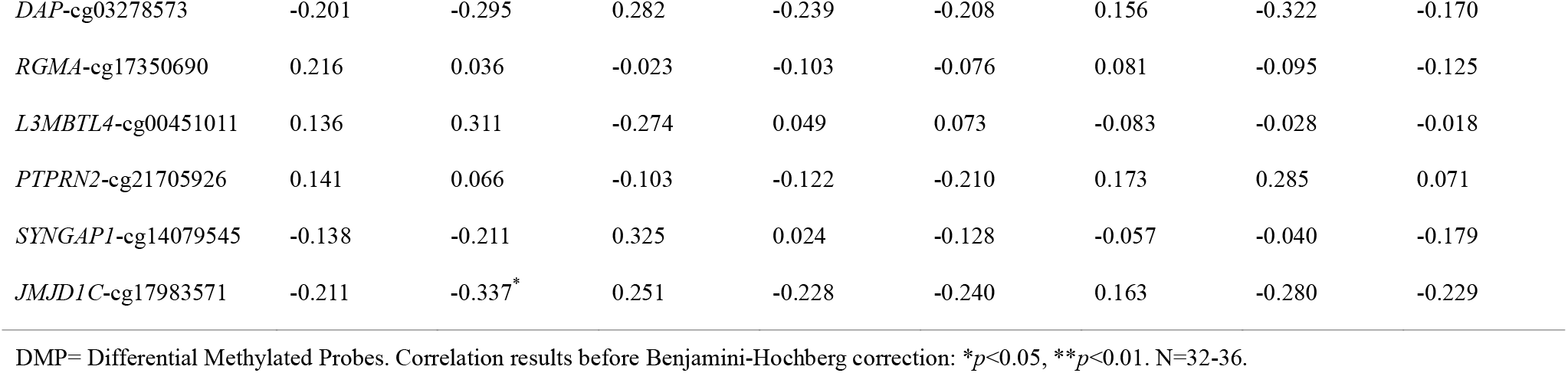
Correlation between MATRICS domain performance and the eighteen selected DMPs in First-Episode Schizophrenic Patients.

**Table 2-b.** Correlation between MATRICS domain performance and the selected DMPs in PRS subjects.

| Characteristics | speed of<br>processing | attention/vigil<br>ance | spatial<br>working<br>memory | verbal<br>learning | visual<br>learning | reason/problem<br>solving | social<br>cognition | overall<br>composite<br>cognition |
| --- | --- | --- | --- | --- | --- | --- | --- | --- |
| *AGTR1*-cg10297223 | 0.125 | 0.178 | -0.066 | -0.230 | 0.144 | 0.192 | 0.235 | 0.156 |
| *DISC1*-cg25215021 | -0.059 | -0.069 | 0.162 | -0.263 | -0.126 | 0.219 | -0.002 | 0.012 |
| *SHANK2*-cg18440889 | 0.123 | 0.081 | -0.023 | -0.181 | 0.136 | 0.263 | 0.100 | 0.109 |
| *BRD1*-cg19984503 | 0.060 | -0.013 | 0.153 | -0.255 | 0.124 | 0.317 | 0.056 | 0.137 |
| *CPEB3*-cg10967735 | -0.004 | -0.056 | 0.122 | -0.432** | 0.217 | 0.262 | 0.109 | 0.113 |
| *EMC2*-cg13442241 | 0.035 | 0.079 | -0.032 | -0.318 | 0.080 | 0.174 | 0.102 | 0.010 |
| *NAPG*-cg22389991 | -0.161 | -0.190 | 0.008 | -0.299 | -0.087 | 0.145 | -0.212 | -0.119 |
| *NR3C1*-cg19645279 | -0.019 | -0.087 | 0.168 | -0.142 | 0.051 | 0.347* | 0.126 | 0.189 |
| *CALN1*-cg04158140 | 0.159 | 0.103 | 0.293 | -0.144 | 0.052 | 0.329 | 0.088 | 0.119 |
| *ROR1*-cg19963700 | 0.148 | 0.129 | 0.202 | -0.180 | 0.152 | 0.314 | 0.115 | 0.195 |
| *NBEA*-cg00500007 | 0.214 | 0.349* | 0.133 | -0.292 | 0.029 | 0.142 | 0.147 | 0.122 |
| *DLG1*-cg07068730 | -0.058 | -0.011 | -0.068 | -0.406* | 0.114 | 0.223 | -0.058 | -0.039 |
| *DAP*-cg03278573 | 0.280 | 0.165 | 0.129 | -0.233 | 0.212 | 0.373* | 0.155 | 0.191 |
| *RGMA*-cg17350690 | -0.034 | -0.106 | -0.325 | -0.368* | 0.102 | -0.085 | -0.280 | -0.281 |
| *L3MBTL4*-cg00451011 | 0.024 | 0.331 | 0.124 | 0.209 | 0.013 | -0.265 | 0.480** | 0.248 |
| *PTPRN2*-cg21705926 | 0.048 | -0.064 | 0.038 | -0.086 | 0.165 | 0.075 | 0.240 | 0.111 |
| *SYNGAP1*-cg14079545 | 0.305 | 0.194 | 0.068 | -0.058 | 0.223 | 0.189 | 0.313 | 0.148 |
| *JMJD1C*-cg17983571 | 0.169 | 0.074 | 0.037 | -0.278 | 0.245 | 0.281 | 0.109 | 0.115 |

DMP= Differential Methylated Probes. PRS=Psychosis risk syndrome. Correlation results before Benjamini-Hochberg correction: \* $p < 0.05$ , \*\* $p < 0.01$ . N=19-20.

There were also significant correlations between PANSS scores and DMP’s β values (Table 3). The strongest and most consistent correlations were with PANSS Depression Factor (*AGRT1, SHANK2, BRD1, EMC2, ROR1, JMJDIC, NBEA, DAP, PTPRN2*) and Excitement Symptom Factor (*AGRT1, DISC1, SHANK2, BRD1, NAPG, CALN1, NR3C1, ROR1, JMJDIC, NBEA, DAP, PTPRN2*), most of which survived BH corrected significance. There were also a few significant correlations with PANSS Total Score (*AGTRI, SHANK2, NAPG, JMJD1C*). These correlation results again suggest that multiple methylation differences in multiple genetic components may align with or influence different dimensions of cognition or symptoms in PRS or FES.

**Table 3.** Correlation between PANSS symptoms and the eighteen selected DMPs in First-Episode Schizophrenic Patients.

| Characteristics | PANSS Total | PANSS Positive<br>Factor | PANSS Negative<br>Factor | PANSS Excitement<br>Factor | PANSS Depression<br>Factor | PANSS Cognitive<br>Factor |
| --- | --- | --- | --- | --- | --- | --- |
| *AGTR1*-cg10297223 | 0.400*<sup>A</sup> | 0.084 | 0.225 | 0.434**<sup>A</sup> | 0.540***<sup>A</sup> | 0.114 |
| *DISC1*-cg25215021 | 0.188 | 0.154 | -0.021 | 0.442**<sup>A</sup> | 0.301 | -0.073 |
| *SHANK2*-cg18440889 | 0.385*<sup>A</sup> | 0.168 | 0.180 | 0.452**<sup>A</sup> | 0.482**<sup>A</sup> | 0.103 |
| *SYNGAP1*-cg14079545 | -0.036 | -0.286 | -0.053 | 0.229 | 0.168 | -0.027 |
| *L3MBTL4*-cg00451011 | -0.190 | -0.094 | -0.087 | -0.245 | -0.111 | -0.074 |
| *BRD1*-cg19984503 | 0.169 | -0.005 | 0.036 | 0.402* | 0.367* | -0.040 |
| *CPEB3*-cg10967735 | 0.181 | -0.123 | 0.117 | 0.231 | 0.154 | 0.194 |
| *EMC2*-cg13442241 | 0.313 | -0.017 | 0.193 | 0.293 | 0.424** | 0.159 |
| *NAPG*-cg22389991 | 0.369* | 0.227 | 0.229 | 0.409* | 0.202 | 0.185 |
| *NR3C1*-cg19645279 | 0.260 | 0.132 | 0.157 | 0.446**<sup>A</sup> | 0.213 | 0.023 |
| *CALN1*-cg04158140 | 0.183 | 0.059 | 0.036 | 0.351* | 0.185 | 0.063 |
| *ROR1*-cg19963700 | 0.307 | 0.145 | 0.065 | 0.453**<sup>A</sup> | 0.517**<sup>A</sup> | 0.035 |
| *JMJD1C*-cg17983571 | 0.360* | 0.130 | 0.172 | 0.414**<sup>A</sup> | 0.442**<sup>A</sup> | 0.100 |
| *NBEA*-cg00500007 | 0.162 | 0.166 | -0.011 | 0.405**<sup>A</sup> | 0.441**<sup>A</sup> | -0.126 |
| *DLG1*-cg07068730 | 0.325 | 0.176 | 0.224 | 0.139 | 0.243 | 0.248 |
| *DAP*-cg03278573 | 0.305 | 0.086 | 0.116 | 0.352** | 0.480**<sup>A</sup> | 0.074 |
| *RGMA*-cg17350690 | 0.012 | -0.191 | 0.131 | -0.135 | -0.241 | 0.096 |
| *PTPRN2*-cg21705926 | -0.141 | -0.082 | 0.037 | -0.413**<sup>A</sup> | -0.539***<sup>A</sup> | 0.218 |

DMP=Differential Methylated Probes. Correlation results before Benjamini-Hochberg correction: \* $p < 0.05$ , \*\* $p < 0.01$ , \*\*\* $p \leq 0.001$ . N= 36. A=significantly different from healthy controls at Benjamini-Hochberg protected significance level ( $\alpha=0.05$ ) tested with the 5 PANSS measures for each DMP in the table.

In validation cohort, we did not find any correlation between methylation level of probes in *SYNGAP1* and *L3MBTL4* with MCCB cognitive performance and symptoms in individuals with FES and PRS.

## Discussion

To our best knowledge, this is the first systematic epigenome-wide association study to determine the phase-specific DNA methylation signatures in Chinese Han subjects. In this study, we observed a predominate DNA hypomethylation profile across the whole genome in FES or PRS groups, which clearly discriminated them from healthy controls. Particularly, the gene *SYNGAP1* showed robust hypomethylation in both PRS and FES patients after vigorous correction, whereas *L3MBTL4* presents characteristic disease-phase specific hypomethylation. Our results strongly suggest a potential value of collective DNA methylation changes for the development of schizophrenia.

In line with our findings, Nishioka et al. also demonstrated that all the top-ranked CpG-sites (603 in total) are hypomethylated ^35^ in peripheral blood DNA from FES patients. We were able to replicate 4 of them (*FEN1, ABL2, MICA, PANX2*) in our experimental setting. In a recent large EWAS study ^36^, 1223 DMPs were identified (*p*< 5 × 10^−5^) and five of them (*SHANK2, ANKRD55, ABL2, DISC1, NF1*) overlap with our findings. The primary factors lead to different results in the reviewed studies may include: a) Distinct microarray platform with featured gene sets or probe numbers; (b) Different cell types (lymphocytes vs whole blood cells); c) Difference in subject characteristics (HAN vs European, or stages of illness); d) Different medication history. Importantly, there was no difference of methylation values for the majority of our 18 DMP selected for bisulfite interrogation between FES in either discovery or validation cohort who were being treated with antipsychotics at the time of assessment vs those on no medication, highlighting a minimal effect of antipsychotics (Supplementary, Table S9). However, it would be proper to adjust for potential drug effects more fully in follow-up larger cohorts.

Wockner et al. examined human frontal cortex from 24 paired schizophrenia patients and controls and found that 4641 probes (corresponding to 2929 unique genes) were differentially methylated with 47.3% CpG sites hypomethylated ^37^. Of the 2929 genes, 38 overlap with our findings in discovery cohort (*WFS1, NBEA, SYT1, NETO2, PANX2*, *RGS7, MANBA, CCDC86, ROR1, FEN1, FASN, CDH5, DAP, HLA-J, DISC1, TSNAX-DISC1, ZNF323, PPP1R11, ATF6, TJP1, GTF2IRD1, EIF2AK2, SYNGAP1, NF1, RGMA, JMJD1C, ABL2, DAZAP1, SHANK2, TSNARE1, STRADA, PDE4DIP, MACF1, NAPG, UGGT2, GPX3, C10orf26, METAP1*). These results indicate that apart from tissue-specific features, peripheral blood cells do share some common methylation signatures with brain. It should be noted that a proportion of the genes discovered by us overlaps with candidate genetic loci identified in previous GWAS or DNA sequencing research (*SHANK2, TJP1, DLG1, NF1, DISC1, CALN1, NAPG, NBEA, PANX2, PEPD*, and *TSNARE1*) ^2,38-42^. DNA methylomic study (*DLG1* and *DISC1*) ^8,38,43,44^ and miRNA interference study (*SHANK2, SYT1, C10orf26*, and *CALN1*) ^45–48^. These shared loci may presumably promote the development and phase transition of schizophrenia.

Kebir et al.^49^ examined the longitudinal DNA methylation changes between converters and non-converters and uncovered two significantly different regions (1q21.1 and promoter region of *GSTM5*) which however was not replicated in our experimental setting. The primary reason may be different study design. Kebir et al. followed the same patient group as they converted from PRS to schizophrenia, whereas we performed cross-sectional study with independent cohorts. A better way to replicate their results would be to follow our PRS subjects for 1-2 years and compare longitudinal methylomic changes between converters and non-converters.

DNA methylation is one of the best-characterized epigenetic mechanisms which affects transcription factor binding and associates with gene expression ^50^ via alternative genomic splicing and promoter usage ^51^. We discovered 460 DMPs across the whole genome of FES, 426 of which (>92.61%) are significantly hypomethylated. Hypomethylation specifically dominates regions of gene body, 3’-untranslated and intergenic regions whereas it is outnumbered by hypermethylation in promoters and 5’-untranlated regions. Site-specific CpG hypermethylation has been frequently reported in promoters of schizophrenia risk genes (reelin, GAD_67_) ^52,53^ as well as some reports of hypomethylation (GRIN2B) ^54^ in promotors ^54^ and enhancer (IGF2) ^55^. Both hypo and hyper-methylation have been reported in loci related to schizophrenia with suicide attempts ^56^, or schizophrenia patients who have experienced the challenge of prenatal adverse hypoxia^57^. One study demonstrates that the hypomethylation of Il-6 promoter region in schizophrenic patients can be reversed by antipsychotic medication ^58^. Additionally, DNA methylation within gene body is also proposed as a feature of active transcription ^59^. Recent research revealed that in human brain, about 34% of CpG islands situated within gene body are methylated ^51,60^ which may serve as alternative promoters ^51^. Further, whole-genome studies have revealed different methylation degree within gene body. In general, introns are more highly hypomethylated than exons, and the level of methylation change at exon–intron boundaries possibly represents a regulatory role of DNA methylation during gene splicing ^61^. Notably, except the 207 shared DMPs, FES patients present more DMPs than PRS (460 vs 372), which strongly indicates a phase-dependent DNA methylation shift. The top disrupted pathways preferentially enriched in MAPK signaling, cellular processes, genetic information processing and inflammation, highlighting their important roles to promote disease development and phase transition.

In our study, we found a stable and significant hypomethylation of *SYNGAP1* in lymphocytes of FES and PRS after stringent filtration. Syngap1 protein is an essential component of N-methyl-D-aspartate receptor (NMDAR) complex abundantly expressed in the postsynaptic density (PSD) of excitatory glutamatergic neurons ^62,63^. It negatively regulates the activities of small GTPases (e.g. Ras, Rap) and controls the scale of growth-related process ^62^. Targeted deletion results in enhanced membrane excitability and disturbed excitation/inhibition balance ^64^, one of the most common mechanisms underpinning neurodevelopmental disorders. Syngap1 haploinsufficiency in mice causes profound core features of schizophrenia such as hyperactivity, decreased prepulse inhibition, impaired working and reference spatial memory ^65,66^, generalized cortical excitability ^67^ and abnormal gamma oscillation ^64^, whereas supplementation of syngap1 protein can reverse or improve the manifestation of behavioral problems, cognitive deficits and medically-refractory seizures ^68,69^ In humans, *de novo* loss-of-function mutations in *SYNGAP1* is one of the most common causes of nonsyndromic sporadic intellectual disability ^70^, and can be also associated with autism spectrum disorder ^71^, epilepsy ^71^, and schizophrenia ^72^. *SYNGAP1* is also proposed as one of the top 10 prioritized genes with susceptibility for schizophrenia ^73^. Decreased SYNGAP1 protein and its associated interacting protein PSD95 is well-documented in brains of schizophrenics ^74^. *SYNGAP1* is also a target of recurrent CNV-overlapped miRNAs which presumably coordinates normal brain development ^75^. Intriguingly, the coding sequence of miRNA-5004 aligns within the introns of *SYNGAP1* (See UCSC hg19). Additionally, miRNA-5004 is presumably implicated in neurodevelopment and has a spectrum of target genes (see Targetscan) ^76,77^ which is often implicated in neurodevelopmental disorders and covers 48 candidate genes listed in our discovery cohort. Therefore, it is likely that *SYNGAP1* may orchestrate brain development and function through multiple mechanisms.

There are several limitations to this study. Firstly, we had a relatively small group size of healthy control in the validation cohort which may reduce our statistical power to replicate the findings in the discovery sample. Secondly, without transcriptional data, it is hard to associate the detected hypomethylation with altered production of mRNA or related proteins. However, the current findings suggest that hypomethylation of *SYNGAP1* itself could be considered as an important risk factor for development of PRS and FES in Han Chinese, and, if replicated elsewhere, in other populations as well. A larger cohort size with additional assessment at transcriptional level would be warranted during follow-up study.

*L3MBTL4* is a newly discovered gene and the link with schizophrenia remains yet established. Recent GWAS studies suggest that *L3MBTL4* profoundly regulates cell cycle ^78,79^ and key substrates including transforming growth factor β, insulin/phosphoinositide 3 kinase, Wnt and epidermal growth factor receptor, which are all risk factors of schizophrenia ^80–84^. Therefore, it may be an attractive biomarker in need of further study and replications in methylation studies as well as functional studies in animal models.

## Conclusions and Implications of Future Studies

Our study indicated that the lymphocytes from peripheral blood of PRS and FES patients displays a unique hypomethylation pattern which discriminates them from healthy controls. Multiple DMPs associated with schizophrenia were identified which highlights the polygenic feature, and a number of the DPMs we identified in the lymphocytes were also found in post-mortem brain studies and matched loci identified from GWAS genetic (SNP) studies and other methylomic studies. This suggests that our methylation findings with the Illumina Infinium Human Methylation 850 BeadChip Array reliably cover many of the gene areas that were found to be important in previous studies.

The advantage of methylation studies using lymphocytes is that they are easily obtained and monitored through the long course of illness, to decipher changes that will lead to the novel treatments or those which persist and worsen after a chronic illness course. It is possible that after pooling results from multiple peripheral blood studies from many regions, certain abnormalities will consistently remain which may link to behavioral, cognitive and electrophysiological and MRI or PET imaging observations. Schizophrenia is a highly heterogeneous disorder and early obtainable methylation markers might help clarify subtypes, when linked to finding of subtypes characterized in living schizophrenics, identified by eye tracking defect, smell deficit, or other abnormalities which present in some but not all schizophrenics. It would also be important to determine whether components of this methylomic metric differ significantly between ethic population such as Han Chinese, other Asian groups, European, and those from African populations. Whether any of these markers can become reliable targets for future drug development needs careful consideration.

## Acknowledgements

This research was supported by the grant 2016YFC1306900 from the National Key Research and Development Program to Dr. Jingping Zhao, and grants 81630033 and 81471363 to Dr. Jingping Zhao and grant 81622018 to Dr. Renrong Wu, from the National Natural Science Foundation of China, and NIH grant 1R01MH101043 to Dr. Alessandro Guidotti, and Stanley foundation grant Z01 to Dr. Jingping Zhao and Dr. Hua Jin. The authors thank Genminix Informatics Ltd., Co for assistance on Genome-wide Infinium Methylation 850 BeadChip Array sequencing analyses and Genergy Bio-technology (shanghai) Co., Ltd for assistance on targeted bisulfite sequencing analyses.

## Author Contribution

YL, GDW, JJW, YL, HLL recruited and assessed all the subjects. YL, HLL performed DNA methylation assessment and acquired data together with GDW, YL, JJW. YL, BL, RCS did statistical analyses and data interpretation. RRW, JPZ, RCS, JMD, ARG, HJ conceived and designed the experiment. RRW, JPZ, RCS supervised the study. YL, BL, RCS drafted the manuscript. RCS, JMD, ARG, HJ, JPZ, RRW critically revised the manuscript. All authors reviewed and approved the final revision.

