## supplementary table s1 to s9 for "Methylomic alteration in peripheral blood lymphocytes of prodromal stage and first-episode Chinese Han schizophrenia patients"

***Supplementary information***

### Supplementary Table S1.

**Primers information.**

| Primer Name | Primer | Product Seq |
| --- | --- | --- |
| AGTR1_F | GAAGTTTTTATTTTTAGTTTGTTGAGTGTTAG | GAAGTTCCTATTCCTAGTTTGTTGAGTGTTAGCATGAAAAAGTGTTGAATTTTGTCCAATAGTTTTTAAAAATTTTTTTAGACAATCATGTAGGCTTTGTCCATTTTTTACTTCTTTAAATTTATTTTTATTGATACACAATAGATGTACACTTTTTAGGTACATGCAATAATTTAATGCCCCTCACTATAAATT**C**GGAGCTGCCTCCT**C**GC**C**GATGATTCCAGCGCCTGACAGCCAGGACCCCAGGCA |
| AGTR1_R | TACCTAAAATCCTAACTATCAAACRCTAAAATC |  |
| BRD1_F | GGTGTGTGTTTTGTGGATTTTAG | GGTGTGTGCCCTGTGGATCCCAGAGGT**C**GGCTTTGCCAACA**C**GGTGTTCAT**C**GAGCCCAT**C**GATGGGGTGAGGAACATCCCTCCAGCC**C**GGTGGAAACTGACATGCTACCTCTGTAAGCAGAAGGG**C**GTGGGTGCCTGCATCCAGTGCCACAAAGCAAACTGCTACACAGCATTCCATGTGA**C**GTGTGCCCAGAAGGCTGGCCTGTACATGAAAATGGAGCC**C**GTGAAGGAACTGACTGGCGGTGGCACCACC |
| BRD1_R | AATAATACCACCRCCAATCAATTC |  |
| CALN1_F | GTAGGATYGGTTTTTAGGGATATTG | GCAGGACCGGCTCTCAGGGACACTGCA**C**GCAGGTG**C**G**C**GTGGAATTTATGGACAACA**C**GAGC**C**GCTCCAGCATC**C**GCAGTGTAAAAGGCCC**C**GTGTGTGAGGGGGA**C**GTGCTCACCCTGTTGGAGTCAGAGCAAGAGGCCTGGAAGTTG**C**GCTGAACTTGGCTGCTCACTGGGTCTTGGATGT |
| CALN1_R | ACATCCAAAACCCAATAAACAAC |  |
| CPEB3_F | TTTTAATGTATTTGTAGAGTTTTYGTTTGAG | TCTTAATGCATTTGTAGAGCTTCCGCTTGAGG**C**GG**C**GAGAAGCCAGGAGCTG**C**G**C**GATGGGGTTCTGGTTGACCAGCAGCTCCTGGTAGGTG**C**GCTCCC**C**GGAACACACCTC**C**GCCTGAGCCT**C**GGGCCCTT**C**GGGATCTGCCTTTATTTTGGTCAT**C**GCAG**C**GGC**C**GCTGAAACCTAGTCCCAGGGAGGCTGC |
| CPEB3_R | ACAACCTCCCTAAAACTAAATTTCAAC |  |
| DAP_F | CCTACTTACTAAAATTTATTTATAACCCCAAAATC | GAAGAAGGCAGAGCACCACCTTCAGCCTCAGCTGGGAAAATGCAGGTTGATTGTCACAAAAAATACTACAAAAAGCCTT**C**GCTGCTCTGCCTGTGTCTGTGAATGACCA**C**GAAGGTGCTA**C**GTGTGTTTGATTTTGGGGTTACAAATAAATTTTAGCAAGTAGG |
| DAP_R | GAAGAAGGTAGAGTATTATTTTTAGTTTTAGTTG |  |
| DISC1_F | ATTATTTTTGATTTAGGATGGGTTTTG | ATCATCCTTGATTTAGGATGGGCCCTGAAATCCAATGATTAGTATTTTTTATGAGAAAGGAGTGGGAGATATGAGACACAGACACACAGAGGGGAATGACATATGAAGATGGAGG**C**GGAGACAGTAGTGAGGCATCTCAAGCCAAGGCATGCTA**C**GGGTTGCTGACAACCACCAGAAGCTGGGAGAGAGGCACGG |
| DISC1_R | CCRTACCTCTCTCCCAACTTCT |  |
| DLG1_F | AAGAGGATGTTTTGTGTTAGAAATAAGAG | AAGAGGATGCCTTGTGTTAGAAATAAGAGCTCAGAGTTGCAAAGAAAATGAGCACTCAAACAAAAGACTTCTCAGCAAGGCAAATTTCAGAAGGGTGCTGCCTG**C**GTTAGTTA**C**GAT**C**GCAAGAGCACACCAAACAAAGGAAAGCAGGGGTTTTTATTCCCAATG |
| DLG1_R | CATTAAAAATAAAAACCCCTACTTTCCT |  |
| EMC2_F | ATAGGTTTTTGTGTGATTGTGGAGT | ACAGGCCTCTGTGTGACTGTGGAGCAATGTGCAGATTGCAGTGATGAGTAGAGTAAAACCTCCACTTTGAGCTCAGCATCTCTGGTGAACACAGGGAACTGCAGGCAACAATGCTGCTGAATACACC**C**GAGTGCACAGTCAGA**C**GTGTTC**C**GTAAA**C**GGTAAATGTGTAAGACCAATTATCTTGAGAGGGTATCTTGCTCCAGAAGATGCT |
| EMC2_R | AACATCTTCTAAAACAAAATACCCTCTC |  |
| JMJD1C_F | CCACACTTATAACAACATTTACTATACACAAC | CTGGAGTGGGCTCCAGAGTTCCTGGGACCAGGTGTTCAC**C**GCCTTCTGGGAGACA**C**G**C**GGGGAC**C**GGAGCT**C**GG**C**GAACACC**C**GC**C**GCCAGGCT**C**GTAGCTCAGTCACCACTGCACCATGCCCAAGCAGCTGCTGC**C**GCCAC**C**GCCATGAGTTGTC**C**GCTGTCTTAATATATCTTAAAAGAGTTGTGTATAGCAAATGCTGTTATAAGTGTGG |
| JMJD1C_R | TTGGAGTGGGTTTTAGAGTTTTTG |  |
| L3MBTL4_F | TAAAAAATGAGAGTAGGTGGTTGTTATG | TAAAAAATGAGAGTAGGTGGCTGCCATGACATAAGGAAAGTGACAATATGAAGCCACTATGCAGACAGGCTCCACATGAGGG**C**GAGTGGTTCCCACAGGA**C**GTTTGGTGTCAGTTTCTGACTTAAATCATTGTGTGTGGCAAGGATTTGGCTAG**C**GTACTGAGTGGTTAAGTTTCAGATAACTAAACTTG |
| L3MBTL4_R | CAAATTTAATTATCTAAAACTTAACCACTCAATAC |  |
| NAPG_F | AAGGATTTAAGTTTTYGGTGAGATTAG | AAGGACCCAAGCCTCCGGTGAGATCAGCCCCCTC**C**GACACCTTCATTT**C**GGCCTGGTGAGACCCTAGCTCTGCTACCTAGCTACACTAGGACCTAGCTACACTGTATCCAGACTTCTGGCCTCATGAAACTGTGAGCTAATCAGTGAGTACTGTTTTAAGCTGCTAAGTTTGTAGCAA |
| NAPG_R | TTACTACAAACTTAACAACTTAAAACAATACTCAC |  |
| NBEA_F | GATTAGATTGAGATAATGAGAGAGTGTAAGAG | GATCAGACTGAGATAATGAGAGAGTGTAAGAGAAATACACCACAAGATATCAAAACACTGATCTTACAGAAGACAAAAATTAGACATAAAAGTAAAAAGGAAATGCAGACAAGGATAAGAGAAAACAATTAAAAGCATAGGAAAACC**C**GTGAAAGAGAATACTACAGAGCTCATAAAAGGAGAGTTAAGAATTCTGTCATAACAGGTGA |
| NBEA_R | TCACCTATTATAACAAAATTCTTAACTCTCCT |  |
| NR3C1_F | TCAAATACCTACAATCAACCACAA | GTGTGAAAACAACTAATACAGATGGCTAGTAAGGGACTAAC**C**GGCAGGGAG**C**GTCTCCAGTGTGGATATGCTGGACAAAGGGATGATTCA**C**GTTCCAGGGCATAAGATTTCATTACTCAGAATTGCACAGAATTTAAAACTTATTAATTATTTCTGGAATTTTCCACTTAATGTTTTCAAACTGTGGTTGACTGCAGGTACCTGA |
| NR3C1_R | GTGTGAAAATAATTAATATAGATGGTTAGTAAGG |  |
| PTPRN2_F | GATTGGTYGGTTGAGTAGTTTTGT | GATTGGCCGGTTGAGCAGTCCTGCTGCAGAA**C**GATGCTTTCCAAGGCATCC**C**GTTGCTTGTGGTCTTCCACCAGCAAGAATGTGGAGGCCAGCCCTGTGAGCTCACAGAGTCACAGTTGTGA**C**GGGAGGGT**C**GCCCCTGGCCATGTGGCCTGTAAGGGG |
| PTPRN2_R | CCCCTTACAAACCACATAACC |  |
| RGMA_F | TTYGAGTTTTGGGTATTGTTAAATGGT | TCCGAGTCCTGGGCACTGTCAAATGGTTCAA**C**GTC**C**GGAATGGTTA**C**GGATTCATCCACAGGAATGA**C**GCCAAGGAAGATGTCTTTGTTCACCAGACAGCTATTAAAAGAAGCCCAGGAAGTTTCTG**C**GCAG**C**GTTGGAGATGAGGAGACTGTGGAATTTGATGT**C**GTGGAAGGAGAGAAGGGTGC |
| RGMA_R | ACACCCTTCTCTCCTTCCAC |  |
| ROR1_F | TTTGGTATTGAAGATTTATGTAMTTTTATGT | TCTGGCATTGAAGACTTATGCAACTTCATGT**C**GTTGAACATGTTCATGACACCATCAGACAGCCACACCAGAGGCCATGTTTCCAGAAGTGAAAAGGAGAGGGCATGGAGAGC**C**GCAGAAGACAAGAGCACTGCAGCCGCTGCTGGGACC |
| ROR1_R | AATCCCAACAACRACTACAATACTCTTAT |  |
| SHANK2_F | ACCCAACCTAAATAAACTCCATAAAATAAC | CTGGATGAACTGGGATGAGGTCAGGCTGTGTT**C**GGGGAGCTGGCACCTGGGCACTGGGCAGGTGTGCCATCAGTTCAGTGTCTGGTGCTTAGATCAGCTGAAACAGAGACAGC**C**GCAT**C**GTGGGGAGAGGTGTCACCTCATGGAGTCCATCCAGGTTGGGC |
| SHANK2_R | TTGGATGAATTGGGATGAGGT |  |
| SYNGAP1_F | TTGTGGTAGGTATTGGTTTTTTTGT | TTGTGGTAGGTATTGGTCTCTCTGCAACTCAGCCATTAATTAGAAATTAGTTTTGAGCCTGAATTTTAAAAAGCCAAGTGTTGCCCCCAGCCCACACACACACACA**C**GGACATGTACAGTACAAACCCCAGATAATTACAACAGCCAAAGAGAGAAGGAAGTGAATTTCC |
| SYNGAP1_R | AAAAATTCACTTCCTTCTCTCTTTAACT |  |

### Supplementary Table S2-a.

**Demographic and clinical characteristics of subjects (discovery cohort)**

| Characteristics | First-episode schizophrenia | | Psychosis risk syndrome | | Healthy controls | | statistics | *p* |
| --- | --- | --- | --- | --- | --- | --- | --- | --- |
|  | N | Mean(S.D.) | N | Mean(S.D.) | N | Mean(S.D.) |  |  |
| age | 40 | 22.2(4.8) | 40 | 21.0(4.3) | 40 | 23.2(2.5) | F=2.928, DF=2,117, | 0.057 |
| gender(M/F) | 40 | 27/13 | 40 | 24/16 | 40 | 24/16 | *chi-square*=0.640 | 0.726 |
| years of education(y) | 36 | 12.1(3.0)*** | 40 | 11.9(2.4)*** | 40 | 16.1(2.1) | F=35.963, DF=2,113 | 0.000 |
| PANSS total | 36 | 72.9(18.3) |  |  |  |  |  |  |
| PANSS positive | 36 | 18.7(5.3) |  |  |  |  |  |  |
| PANSS negative | 36 | 19.0(7.4) |  |  |  |  |  |  |
| PANSS general | 36 | 35.3(9.5) |  |  |  |  |  |  |
| SIPS total |  |  | 40 | 33.6(17.9) |  |  |  |  |
| SIPS positive |  |  | 40 | 8.6(4.9) |  |  |  |  |
| SIPS negative |  |  | 39 | 7.2(6.0) |  |  |  |  |
| SIPS disrupted |  |  | 40 | 3.9(2.7) |  |  |  |  |
| SIPS general |  |  | 40 | 4.2(3.0) |  |  |  |  |
| Medication status at time of testing |  |  |  |  |  |  |  |  |
| Antipsychotic | 29 |  |  |  |  |  |  |  |
| Traditional (T) | 3 |  |  |  |  |  |  |  |
| Atypical (A) | 26 |  |  |  |  |  |  |  |
| Antidepressant | 0 |  |  |  |  |  |  |  |
| Valproate | 3 |  |  |  |  |  |  |  |
| Benzodiazepine | 4 |  |  |  |  |  |  |  |
| Antiseizure | 4 |  |  |  |  |  |  |  |
| Phenobarbital | 0 |  |  |  |  |  |  |  |
| None medicated | 11 |  |  |  |  |  |  |  |

Statistical significance of individual demographic and clinical feature in each group was calculated upon analysis of variance. S.D.=Standard Deviation. ^***^ *p*<0.001.

### Supplementary Table S2-b.

**Demographic and clinical characteristics of subjects (validation cohort)**

| Characteristics | First-episode schizophrenia | | Psychosis risk syndrome | | Healthy controls | | statistics | *p* |
| --- | --- | --- | --- | --- | --- | --- | --- | --- |
|  | N | Mean(S.D.) | N | Mean(S.D.) | N | Mean(S.D.) |  |  |
| age | 39 | 24.7(7.4) | 41 | 21.0(5.1) | 10 | 23.2(2.9) | F=3.714, DF=2, 87 | 0.028 |
| gender(M/F) | 39 | 17/22 | 41 | 18/23 | 10 | 5/5 | *chi square*=0.141,  DF=2 | 0.932 |
| years of education(y) | 37 | 11.4(3.3)*** | 41 | 11.5(3.7)** | 10 | 15.5(2.2) | F=6.064, DF=2, 85 | 0.003 |
| PANSS total | 38 | 81.7(18.6) |  |  |  |  |  |  |
| PANSS positive | 38 | 21.2(4.7) |  |  |  |  |  |  |
| PANSS negative | 38 | 20.5(8.1) |  |  |  |  |  |  |
| PANSS general | 38 | 40.0(9.6) |  |  |  |  |  |  |
| SIPS total |  |  | 40 | 37.4(17.6) |  |  |  |  |
| SIPS positive |  |  | 40 | 8.7(3.7) |  |  |  |  |
| SIPS negative |  |  | 40 | 10.8(5.9) |  |  |  |  |
| SIPS disrupted |  |  | 40 | 5.2(3.1) |  |  |  |  |
| SIPS general |  |  | 40 | 6.1(3.6) |  |  |  |  |
| Medication status at time of testing |  |  |  |  |  |  |  |  |
| Antipsychotic | 19 |  |  |  |  |  |  |  |
| Traditional (T) | 1 |  |  |  |  |  |  |  |
| Atypical (A) | 18 |  |  |  |  |  |  |  |
| Antidepressant | 0 |  |  |  |  |  |  |  |
| Valproate | 0 |  |  |  |  |  |  |  |
| Benzodiazepine | 0 |  |  |  |  |  |  |  |
| Antiseizure | 0 |  |  |  |  |  |  |  |
| Phenobarbital | 0 |  |  |  |  |  |  |  |
| None medicated | 20 |  |  |  |  |  |  |  |

Statistical significance of individual demographic and clinical feature in each group was calculated upon analysis of variance. S.D.=Standard Deviation. ^***^ *p*<0.001, ***p*<0.01.

### Supplementary Table S3-a.

**Scores on Cognitive Domains for Healthy Controls and First-episode schizophrenia (discovery cohort)**

| Domain Score | Healthy controls  (N’s=36-38) | Psychosis risk syndrome  (N’s=31-35) | First-episode schizophrenia  (N’s =32-36) | Overall Test GLM  F, DF, *p* |
| --- | --- | --- | --- | --- |
| Speed of Processing | 46.78±5.52 | 38.77±10.70*** | 36.04±9.83*** | F=14.616, DF=2, 106, *p*<0.000 |
| Attention / Vigilance | 46.08± 8.76 | 49.17 ±12.89 | 41.97±12.61 | F=3.482, DF=2, 105, *p*=0.034 |
| Spatial Working Memory | 43.42±9.23 | 39.31±10.58 | 36.31±10.88** | F=4.515, DF=2, 106, *p*=0.013 |
| Verbal Learning | 39.26±10.54 | 35.83±15.56 | 30.03±12.21** | F=4.838, DF=2, 106, *p*=0.010 |
| Visual Learning | 45.64±8.60 | 43.00±17.18 | 37.87± 12.00* | F=3.413, DF=2, 106, *p*=0.037 |
| Reasoning / Problem Solving | 50.03±9.65 | 40.06±12.14*** | 40.19±12.27*** | F=9.374, DF=2, 106, *p*<0.000 |
| Social Cognition | 53.89±12.92 | 50.16±13.37 | 46.06±14.02* | F=3.014, DF=2, 100, *p*=0.054 |
| Overall Composite  Score | 52.82±5.13 | 48.48±9.51* | 43.90±7.26*** | F=12.290, DF=2, 96, *p*<0.000 |

Statistical significance of individual component of each group was defined upon comparison with control group by Dunnett’s Test (two sided). Each number presents Mean ± S.D. of Overall Composite or Domain Test scores adjusted for age, sex, and education based on Chinese norms. S.D.=Standard Deviation. *** *p*<.0.001, ***p*<0.01, **p*<0.05.

### Supplementary Table S3-b.

**Description of cognitive performance of subjects (validation cohort)**

| Cognitive items | Healthy controls  (N’s=10) | Psychosis risk syndrome  (N’s =30-41) | First-episode schizophrenia  (N’s =10-26) | F, DF, *p* |
| --- | --- | --- | --- | --- |
| TMT-A | 30.40±10.06 | 44.51±19.82 | 46.95±22.42* | F=2.621, DF=2,74, *p*=0.080 |
| BACS-SC | 64.50±7.66 | 54.12±11.35* | 47.38±12.79*** | F=8.298, DF=2,74, *p*=0.001 |
| FLUENCY | 25.70±5.58 | 19.17±5.34** | 16.54±6.99*** | F=8.490, DF=2,74, *p*=0.000 |
| CPT-IP | 2.93± 0.56 | 2.26±0.76* | 1.88±0.39** | F=6.208, DF=2,57, *p*=0.004 |
| WMS | 16.80±2.66 | 15.49±3.47 | 15.58±2.40 | F=0.768, DF=2,74, *p*=0.468 |
| HVLT-R | 26.10±4.70 | 23.68±6.30 | 19.50±6.14** | F=5.697, DF=2,74, *p*=0.005 |
| BVMT-R | 29.10±3.18 | 26.02± 6.36 | 21.50± 7.54** | F=6.231, DF=2,73, *p*=0.003 |
| NAB-M | 18.30±4.76 | 14.02±5.68 | 12.92±6.38* | F=3.120, DF=2,74, *p*=0.050 |
| MSCEIT | 93.00±20.78 | 84.30±12.83 | 82.64±11.72 | F=2.084, DF=2,63, *p*=0.133 |

TMT-A=Trail-Making Test: Part A; BACS-SC=Brief Assessment of Cognition in Schizophrenia: Symbol Coding; WMS=Wechsler Memory Scale-III: Spatial Span Test; FLUENCY=Category Fluency: Animal Naming; HVLT-R=Hopkins Verbal Learning Test-Revised; NAB-M=Neuropsychological Assessment Battery: Mazes; BVMT-R=Brief Visuospatial Memory Test-Revised; MSCEIT=Mayer-Salovey-Caruso Emotional Intelligence Test: Managing Emotions; CPT-IP=Continuous Performance Test-Identical Pairs. Statistical significance of individual assessment value of each group was defined upon comparation with control group by Dunnett’s Test (two sided). Each number represents Mean ± S.D. of raw scores of each item. S.D.=Standard Deviation. *** *p*<.001, ***p*<0.01, **p*<0.05.

### Supplementary Table S4-a.

**The differentially methylated probes between patients with schizophrenia and controls.**

| Probe ID | Delta Beta | Adjust *P* Value | Chromosome | Gene Name | Gene Context | CpG Island |
| --- | --- | --- | --- | --- | --- | --- |
| cg19476594 | -0.092367054 | 6.93E-11 | 1 | ABL2;ABL2;ABL2;ABL2;ABL2;ABL2;ABL2;ABL2 | Body;Body;Body;Body;Body;Body;Body;Body |  |
| cg11853096 | -0.086586082 | 8.07E-13 | 1 | AGBL4 | Body |  |
| cg10297223 | -0.136220908 | 2.05E-17 | 3 | AGTR1;AGTR1;AGTR1;AGTR1 | TSS1500;TSS1500;TSS1500;TSS1500 | N_Shore |
| cg15004555 | -0.095103951 | 5.74E-08 | 1 | AIM2 | Body |  |
| cg08421459 | -0.081446483 | 0.004308406 | 9 | AK8 | Body |  |
| cg03590545 | -0.097837235 | 7.59E-10 | 4 | ANAPC10;ANAPC10;ANAPC10;ANAPC10;ANAPC10;ANAPC10;ANAPC10;ANAPC10;ANAPC10 | Body;Body;Body;Body;Body;Body;Body;Body;Body |  |
| cg20047349 | -0.093153994 | 0.015344926 | 17 | ANKFN1 | Body |  |
| cg20184537 | -0.082922475 | 8.72E-12 | 2 | ANKMY1;ANKMY1;ANKMY1;ANKMY1;ANKMY1;ANKMY1 | Body;Body;Body;Body;Body;Body |  |
| cg17049825 | -0.088254939 | 0.034920293 | 9 | ANKRD18B | TSS1500 | N_Shore |
| cg15431103 | -0.122083221 | 1.03E-05 | 5 | ANKRD55 | Body |  |
| cg15375051 | 0.080967846 | 3.05E-06 | 4 | APBB2;APBB2;APBB2 | 5'UTR;5'UTR;5'UTR |  |
| cg18794058 | -0.082738936 | 5.59E-05 | 17 | ARL17A | TSS1500 |  |
| cg12289553 | -0.091390164 | 1.44E-14 | 10 | ASAH2B | 5'UTR | S_Shore |
| cg14282460 | -0.087023986 | 0.018432305 | 8 | ASAP1;ASAP1 | Body;Body |  |
| cg15735386 | -0.08890573 | 0.011676865 | 1 | ATF6 | TSS1500 |  |
| cg26840922 | -0.082711832 | 3.70E-11 | 3 | ATP1B3 | Body |  |
| cg20312390 | -0.084523092 | 0.000124366 | 1 | ATP2B4;ATP2B4 | 5'UTR;5'UTR |  |
| cg12953621 | -0.100382676 | 2.05E-16 | 12 | ATP5G2;ATP5G2 | Body;Body |  |
| cg14922317 | 0.083874523 | 0.009553075 | 16 | ATXN2L;ATXN2L;ATXN2L;ATXN2L;ATXN2L;ATXN2L;ATXN2L;ATXN2L;ATXN2L;ATXN2L;ATXN2L;ATXN2L;ATXN2L;ATXN2L | 5'UTR;5'UTR;5'UTR;5'UTR;5'UTR;5'UTR;5'UTR;1stExon;1stExon;1stExon;1stExon;1stExon;1stExon;1stExon | Island |
| cg23525965 | -0.083288966 | 0.000276079 | 6 | BEND3 | 3'UTR | N_Shelf |
| cg18366541 | -0.081965755 | 0.039172166 | 17 | BPTF;BPTF | Body;Body |  |
| cg19984503 | -0.080284221 | 9.96E-15 | 22 | BRD1;BRD1 | Body;Body | N_Shore |
| cg07537139 | -0.087770414 | 0.040035809 | 9 | BRINP1 | Body |  |
| cg15227982 | -0.084581907 | 0.006981396 | 10 | C10orf26;C10orf26 | TSS200;Body |  |
| cg13961165 | -0.10760537 | 1.81E-11 | 2 | C2orf27A;C2orf27A | 1stExon;5'UTR |  |
| cg00435273 | -0.098118193 | 0.035123007 | 2 | C2orf76 | 5'UTR |  |
| cg26670636 | -0.084153753 | 2.47E-09 | 7 | C7orf61 | TSS1500 |  |
| cg17960959 | -0.089971022 | 0.04033597 | 7 | C7orf73 | TSS1500 | N_Shore |
| cg07804024 | -0.1154375 | 0.009453592 | 7 | C7orf73 | TSS1500 | N_Shore |
| cg04158140 | -0.08531868 | 6.02E-13 | 7 | CALN1;CALN1 | Body;Body |  |
| cg00356834 | -0.162642408 | 0.006763915 | 7 | CARD11 | 5'UTR |  |
| cg19114543 | -0.082879824 | 0.002821945 | 3 | CCDC12 | Body |  |
| cg08611431 | -0.088924857 | 1.90E-06 | 7 | CCDC126 | Body |  |
| cg22516161 | -0.086426795 | 0.002008862 | 17 | CCDC144NL | TSS200 | Island |
| cg02941728 | -0.099992049 | 8.05E-15 | 11 | CCDC86 | 3'UTR | N_Shelf |
| cg19972287 | -0.080002753 | 4.95E-16 | 10 | CDC10L | Body | N_Shelf |
| cg08066417 | -0.093277382 | 4.95E-13 | 16 | CDH5 | 5'UTR |  |
| cg08152083 | -0.098695455 | 6.08E-15 | 17 | CDRT15L2 | TSS1500 |  |
| cg03388324 | -0.089851062 | 3.21E-12 | 6 | CEP85L;CEP85L;CEP85L | Body;Body;Body |  |
| cg10182306 | -0.093885063 | 7.82E-10 | 1 | CLCNKA;CLCNKA;CLCNKA | Body;Body;Body |  |
| cg10967735 | -0.084245067 | 1.02E-14 | 10 | CPEB3 | Body |  |
| cg22682254 | -0.091519608 | 1.57E-09 | 17 | CRYBA1 | TSS1500 |  |
| cg15411873 | -0.08740597 | 0.017357722 | 2 | CRYGD;LOC100507443 | TSS1500;Body | S_Shore |
| cg24141198 | -0.082426094 | 2.86E-10 | 1 | CSDE1;CSDE1;NRAS;CSDE1 | 3'UTR;3'UTR;TSS1500;3'UTR |  |
| cg24655310 | -0.116657204 | 0.000120294 | 19 | CYP4F11;CYP4F11 | 1stExon;Body |  |
| cg22669123 | 0.094228563 | 3.36E-05 | 19 | CYP4F12;CYP4F12 | TSS1500;TSS1500 |  |
| cg03278573 | -0.088252237 | 3.36E-06 | 5 | DAP;DAP | Body;Body |  |
| cg07825294 | -0.08789672 | 0.001143093 | 19 | DAZAP1;DAZAP1 | Body;Body | N_Shelf |
| cg00574307 | -0.08272064 | 4.27E-06 | 13 | DCT;DCT | Body;Body |  |
| cg13795666 | 0.085482304 | 2.61E-07 | 1 | DDX59 | Body |  |
| cg24860332 | -0.091810465 | 0.005644438 | 8 | DEFA10P | TSS1500 |  |
| cg26423283 | -0.093040669 | 1.27E-07 | 8 | DEFA4 | TSS1500 |  |
| cg07341559 | -0.112743213 | 3.09E-13 | 11 | DEFB108B | Body |  |
| cg14783954 | -0.092060777 | 4.19E-05 | 8 | DENND3 | Body |  |
| cg25215021 | -0.080537206 | 9.08E-16 | 1 | DISC1-IT1;DISC1;DISC1;DISC1;DISC1;DISC1;DISC1;DISC1;DISC1;DISC1;TSNAX-DISC1 | TSS200;Body;Body;Body;Body;Body;Body;Body;Body;Body;Body |  |
| cg07068730 | -0.080760083 | 2.32E-08 | 3 | DLG1;DLG1;DLG1;DLG1 | Body;Body;Body;Body |  |
| cg00093095 | -0.085039187 | 0.021496847 | 17 | DNAH17 | Body |  |
| cg06871323 | -0.097448995 | 6.64E-12 | 7 | DPY19L1 | Body |  |
| cg00737302 | -0.080364303 | 1.94E-13 | 10 | DUSP5 | 3'UTR |  |
| cg00218840 | -0.103496422 | 2.05E-17 | 1 | DUSP5P1;RHOU | Body;Body | S_Shore |
| cg27224855 | -0.09149166 | 2.27E-13 | 6 | E2F3;E2F3 | 3'UTR;3'UTR |  |
| cg05116169 | 0.082948063 | 5.79E-05 | 6 | ECT2L;ECT2L | 5'UTR;5'UTR |  |
| cg26587553 | 0.088904555 | 0.001662153 | 12 | EEA1 | Body |  |
| cg02449698 | -0.086633067 | 1.00E-14 | 1 | EFCAB7;DLEU2L | Body;TSS200 |  |
| cg18803079 | -0.122941269 | 3.68E-17 | 1 | EFCAB7;DLEU2L | Body;TSS200 |  |
| cg18091465 | -0.084174899 | 2.18E-16 | 15 | EFTUD1;EFTUD1 | Body;Body | N_Shelf |
| cg22558269 | -0.080460479 | 4.37E-08 | 2 | EHD3 | TSS1500 | N_Shore |
| cg05332270 | -0.082003717 | 0.000941137 | 2 | EIF2AK2;EIF2AK2 | TSS1500;TSS1500 | Island |
| cg00418190 | -0.097172282 | 0.018335774 | 12 | ELK3 | Body |  |
| cg14575022 | -0.11524992 | 0.025336667 | 12 | ELK3 | Body |  |
| cg18756775 | -0.122462989 | 0.030529738 | 12 | ELK3 | Body |  |
| cg01927686 | -0.123832676 | 0.029254381 | 12 | ELK3 | Body |  |
| cg13442241 | -0.104415309 | 4.92E-14 | 8 | EMC2 | Body |  |
| cg00084591 | -0.085040399 | 1.71E-11 | 9 | ERP44 | Body |  |
| cg04217177 | -0.083484168 | 0.006973169 | 1 | ESRRG | 5'UTR | N_Shore |
| cg20689962 | -0.108486878 | 0.022421964 | 1 | EYA3;EYA3;EYA3;EYA3;EYA3 | Body;5'UTR;5'UTR;5'UTR;5'UTR |  |
| cg06051442 | -0.081981149 | 5.65E-09 | 9 | FAM120A;FAM120A;FAM120A;FAM120A | Body;Body;Body;Body | S_Shore |
| cg14115740 | -0.105248446 | 1.60E-13 | 9 | FANCC | 5'UTR | Island |
| cg25825740 | -0.087793133 | 4.92E-16 | 17 | FASN | 3'UTR | Island |
| cg18370902 | -0.087879273 | 3.03E-11 | 8 | FBXO25;FBXO25;FBXO25 | 5'UTR;Body;Body |  |
| cg13601636 | -0.095445755 | 6.47E-14 | 19 | FCAR;FCAR;FCAR;FCAR;FCAR;FCAR;FCAR;FCAR;FCAR | TSS1500;TSS1500;TSS1500;TSS1500;TSS1500;TSS1500;TSS1500;TSS1500;TSS1500 |  |
| cg14655485 | -0.08249689 | 6.91E-16 | 11 | FEN1 | Body | S_Shelf |
| cg25634666 | -0.080876164 | 0.001092173 | 11 | FOLR3;FOLR3 | 5'UTR;1stExon |  |
| cg01480180 | -0.089229502 | 0.037506098 | 7 | FZD1;FZD1 | 1stExon;3'UTR | Island |
| cg14449611 | -0.087409344 | 6.83E-07 | 2 | GALNT5 | Body |  |
| cg00872998 | -0.109066407 | 6.51E-17 | 9 | GDA;GDA;GDA;GDA | Body;Body;Body;Body |  |
| cg21086969 | 0.09022743 | 1.06E-07 | 6 | GMPR | Body |  |
| cg05531670 | -0.088248703 | 1.80E-07 | 15 | GOLGA8B | Body |  |
| cg17261508 | -0.086574525 | 0.004603501 | 5 | GPX3 | TSS1500 | N_Shore |
| cg22525688 | -0.082711679 | 0.0001444 | 7 | GTF2IRD1;GTF2IRD1 | Body;Body |  |
| cg03289961 | -0.107828407 | 7.41E-06 | 2 | HADHB;HADHB;HADHB;HADHB;HADHB | ExonBnd;ExonBnd;Body;Body;Body |  |
| cg08782481 | 0.084779634 | 7.33E-08 | 6 | HIST1H3I | 1stExon | Island |
| cg09312328 | -0.140579831 | 0.04956003 | 6 | HLA-DPA1;HLA-DPA1;HLA-DPA1 | Body;Body;Body |  |
| cg04628742 | -0.08794752 | 0.000327025 | 6 | HLA-J;NCRNA00171 | TSS1500;Body | N_Shore |
| cg14083800 | -0.10477593 | 2.88E-06 | 10 | HNRNPF;HNRNPF;HNRNPF;HNRNPF;HNRNPF | 5'UTR;1stExon;5'UTR;5'UTR;5'UTR | Island |
| cg11962649 | -0.099931915 | 4.37E-12 | 9 | HSPA5 | Body | N_Shore |
| cg00470768 | -0.098431482 | 2.35E-13 | 15 | INO80 | Body |  |
| cg15452937 | -0.113446133 | 1.61E-08 | 5 | IPO11-LRRC70;IPO11;IPO11 | Body;Body;Body |  |
| cg17983571 | -0.121489387 | 7.25E-10 | 10 | JMJD1C | Body |  |
| cg26296769 | -0.119955214 | 4.98E-14 | 6 | KLC4;KLC4;KLC4;MRPL2;MRPL2 | TSS1500;TSS1500;TSS1500;Body;Body | N_Shore |
| cg02990553 | -0.092072254 | 3.83E-10 | 17 | KRT28 | 1stExon | S_Shelf |
| cg04204557 | -0.108746215 | 1.61E-15 | 21 | KRTAP10-11;C21orf29 | TSS1500;Body |  |
| cg00451011 | -0.107437879 | 0.002898391 | 18 | L3MBTL4 | 5'UTR |  |
| cg08822578 | -0.085009508 | 7.90E-05 | 13 | LATS2 | Body | N_Shelf |
| cg02256315 | -0.110329218 | 2.36E-05 | 11 | LDHAL6A;LDHAL6A | 3'UTR;3'UTR |  |
| cg00114966 | -0.109276641 | 0.033311832 | 1 | LHX9;LHX9 | Body;Body | S_Shelf |
| cg24203469 | -0.101434549 | 2.79E-11 | 7 | LINC01162 | TSS200 |  |
| cg03017824 | -0.112934953 | 7.97E-16 | 1 | LOC100130093;LOC100130093 | TSS1500;TSS1500 | N_Shore |
| cg04676559 | -0.080945621 | 8.05E-15 | 17 | LOC100131347;PLXDC1 | Body;Body |  |
| cg02414646 | -0.084299688 | 1.22E-11 | 10 | LOC100216001 | TSS1500 |  |
| cg09141640 | -0.093810406 | 3.92E-11 | 22 | LOC100506679 | TSS1500 |  |
| cg23355197 | -0.092384626 | 2.37E-07 | 8 | LOC100506990;LOC100506990 | Body;Body |  |
| cg14008271 | -0.083647214 | 0.035522811 | 12 | LOC100507195 | TSS1500 |  |
| cg15237618 | -0.088223618 | 0.035439332 | 2 | LOC100996579;LOC100996579;LOC101929512 | TSS1500;TSS1500;Body | N_Shore |
| cg13593809 | -0.111552963 | 4.47E-09 | 17 | LOC101559451;PELP1;PELP1 | Body;TSS1500;TSS1500 | S_Shore |
| cg24674363 | -0.136128095 | 7.10E-13 | 3 | LOC101927056;LOC101927056 | Body;Body |  |
| cg22584296 | -0.084575293 | 1.35E-09 | 4 | LOC101927237 | TSS1500 |  |
| cg03472399 | -0.098729038 | 2.36E-10 | 13 | LOC101928697 | Body |  |
| cg04053260 | -0.107323158 | 0.011303898 | 14 | LOC101929718;SLC39A2;SLC39A2;SLC39A2;SLC39A2 | TSS1500;Body;Body;ExonBnd;ExonBnd |  |
| cg15207224 | 0.081180713 | 0.000416476 | 1 | LOC105748977 | Body |  |
| cg07240877 | -0.091996421 | 5.98E-12 | 1 | LOC339535 | TSS200 |  |
| cg06774087 | -0.081233502 | 2.13E-10 | 5 | LOC441089 | Body |  |
| cg12361190 | -0.082445116 | 2.77E-07 | 6 | LOC441155;EYS;EYS | TSS200;Body;Body | Island |
| cg21581312 | -0.094370051 | 1.75E-05 | 15 | LOC723972 | TSS200 |  |
| cg02356786 | -0.084037383 | 0.037862816 | 1 | LOC731275 | Body | Island |
| cg10299917 | -0.095647935 | 6.02E-16 | 22 | LRP5L | 5'UTR |  |
| cg08701935 | -0.086815572 | 1.61E-07 | 10 | LRRC20;LRRC20;LRRC20;LRRC20;LRRC20;LRRC20;LRRC20;LRRC20;LRRC20 | 5'UTR;Body;Body;Body;Body;Body;Body;Body;Body |  |
| cg03318469 | -0.081466905 | 0.004272104 | 1 | LRRC47 | Body | N_Shelf |
| cg23353000 | -0.090664021 | 4.63E-07 | 17 | LSM12 | Body |  |
| cg00586123 | -0.099174695 | 0.000247997 | 6 | LY86-AS1 | Body |  |
| cg03900366 | -0.107910912 | 1.98E-13 | 1 | MACF1 | Body |  |
| cg25960044 | -0.107654805 | 2.12E-08 | 4 | MANBA | Body |  |
| cg10006801 | -0.102957126 | 8.00E-12 | 17 | MAP2K3;MAP2K3 | Body;Body |  |
| cg12045156 | -0.10246854 | 5.59E-15 | 17 | MAP2K6 | Body |  |
| cg18806716 | -0.082004097 | 0.029305901 | 10 | MAP3K8 | TSS1500 | N_Shore |
| cg01490296 | -0.115527454 | 1.55E-13 | 10 | MCM10;MCM10 | 5'UTR;5'UTR | S_Shore |
| cg02141602 | -0.094843316 | 3.84E-12 | 19 | MEIS3;MEIS3;MEIS3 | TSS200;TSS200;TSS200 | S_Shore |
| cg07369840 | -0.087130689 | 4.97E-13 | 4 | METAP1 | Body |  |
| cg02173085 | -0.116147396 | 1.32E-10 | 2 | METTL5;METTL5;METTL5 | Body;Body;Body | N_Shelf |
| cg25552573 | -0.091054419 | 0.04360282 | 6 | MICA;MICA;MICA | TSS1500;TSS1500;TSS1500 | N_Shore |
| cg08477687 | -0.083526808 | 0.039911851 | 1 | MIR1977 | TSS1500 |  |
| cg09828968 | -0.099246981 | 8.81E-09 | 9 | MIR3134 | Body | S_Shelf |
| cg09730926 | -0.088739622 | 2.05E-05 | 1 | MIR5087;NBPF8;NBPF8;NBPF8 | TSS1500;Body;Body;Body | S_Shore |
| cg05216534 | 0.08781881 | 9.34E-09 | 8 | MIR548O2 | Body |  |
| cg02303386 | -0.097038859 | 2.10E-10 | 16 | MLKL;MLKL | TSS1500;TSS1500 | S_Shore |
| cg22431028 | -0.135127887 | 2.14E-16 | 14 | MNAT1;MNAT1 | Body;Body |  |
| cg03409548 | -0.083260859 | 1.75E-06 | 1 | MOBKL2C | TSS200 | S_Shore |
| cg05957730 | -0.092518999 | 1.18E-13 | 2 | MPHOSPH10 | Body | S_Shelf |
| cg07788360 | -0.080792384 | 8.58E-09 | 13 | MRPS31 | Body |  |
| cg19105504 | -0.081313805 | 5.14E-11 | 11 | MS4A15;MS4A15;MS4A15;MS4A15 | 5'UTR;1stExon;5'UTR;1stExon |  |
| cg06813806 | -0.096804494 | 1.80E-05 | 7 | MTURN | Body |  |
| cg16074739 | -0.099778584 | 1.31E-09 | 8 | MTUS1;MTUS1;MTUS1;MTUS1 | TSS1500;TSS1500;Body;Body |  |
| cg07074586 | -0.105513552 | 5.82E-16 | 12 | MYL6;MYL6 | 3'UTR;3'UTR |  |
| cg03889742 | -0.110322958 | 6.59E-10 | 15 | MYO5A;MYO5A | Body;Body |  |
| cg22389991 | -0.081616267 | 1.94E-13 | 18 | NAPG | Body |  |
| cg00500007 | -0.10983917 | 1.54E-08 | 13 | NBEA | Body |  |
| cg24982565 | -0.081154979 | 4.81E-07 | 1 | NBPF20;NBPF10;NBPF10 | Body;Body;Body |  |
| cg04711526 | -0.115162839 | 2.81E-12 | 14 | NDUFB1 | 3'UTR |  |
| cg15566570 | -0.084805799 | 3.81E-09 | 16 | NETO2;NETO2 | Body;Body |  |
| cg21323738 | -0.085073311 | 0.006796667 | 17 | NF1;NF1;NF1 | Body;Body;Body |  |
| cg04674762 | -0.084619004 | 0.014315518 | 14 | NID2 | Body |  |
| cg11792281 | -0.120197469 | 6.98E-08 | 17 | NLK | Body |  |
| cg27188056 | -0.080052481 | 0.000408654 | 19 | NLRP12 | TSS200 |  |
| cg19645279 | -0.089668175 | 4.54E-13 | 5 | NR3C1;NR3C1;NR3C1;NR3C1;NR3C1;NR3C1;NR3C1;NR3C1;NR3C1;NR3C1;NR3C1;NR3C1;NR3C1;NR3C1;NR3C1 | Body;Body;Body;Body;Body;Body;Body;Body;Body;Body;Body;Body;Body;Body;Body |  |
| cg12512012 | -0.080531751 | 9.01E-09 | 7 | NUDCD3 | Body |  |
| cg07986187 | -0.094909663 | 4.51E-09 | 10 | NUDT5 | Body |  |
| cg22273354 | -0.094660296 | 0.008226641 | 11 | NUP160 | Body |  |
| cg07585732 | -0.088638375 | 0.001783549 | 9 | NUTM2G;NUTM2G | Body;Body |  |
| cg14762436 | -0.092660617 | 0.019929938 | 7 | OSBPL3;OSBPL3;OSBPL3;OSBPL3 | Body;Body;Body;Body |  |
| cg23697630 | -0.102306908 | 0.003410455 | 2 | OXER1 | TSS200 |  |
| cg11652329 | -0.081247397 | 9.44E-13 | 22 | P2RX6P;LRRC74B;LRRC74B;LRRC74B | TSS1500;TSS1500;TSS1500;TSS1500 | N_Shore |
| cg13126076 | -0.088967844 | 0.008844712 | 8 | PABPC1 | Body | N_Shelf |
| cg07895657 | -0.108481024 | 0.003500338 | 22 | PANX2;PANX2;PANX2 | Body;Body;Body | Island |
| cg21015022 | 0.207453616 | 0.045215556 | 7 | PARP12 | TSS200 | S_Shore |
| cg06649000 | -0.109699561 | 3.36E-15 | 12 | PCBP2;PCBP2;PCBP2;PCBP2;PCBP2;PCBP2;PCBP2 | Body;Body;Body;Body;Body;Body;Body |  |
| cg22552448 | -0.085927243 | 1.64E-14 | 12 | PCBP2;PCBP2;PCBP2;PCBP2;PCBP2;PCBP2;PCBP2;PCBP2;PCBP2;PCBP2;PCBP2;PCBP2;PCBP2;PCBP2 | ExonBnd;ExonBnd;ExonBnd;ExonBnd;ExonBnd;ExonBnd;ExonBnd;Body;Body;Body;Body;Body;Body;Body | S_Shelf |
| cg07993731 | -0.080016477 | 1.21E-10 | 5 | PDCD6;PDCD6;PDCD6;PDCD6;PDCD6 | 5'UTR;Body;Body;Body;Body |  |
| cg18384674 | -0.082966787 | 6.30E-16 | 1 | PDE4DIP;PDE4DIP;PDE4DIP;PDE4DIP;PDE4DIP;NBPF20;NBPF9 | Body;Body;Body;Body;Body;Body;Body |  |
| cg14102005 | -0.086903455 | 5.64E-11 | 14 | PELI2 | Body |  |
| cg01625851 | -0.099691562 | 0.033278762 | 19 | PEPD;PEPD;PEPD | Body;Body;Body |  |
| cg06936779 | 0.134869591 | 0.000217224 | 1 | PIP5K1A;PIP5K1A;PIP5K1A;PIP5K1A;PIP5K1A;PIP5K1A;PIP5K1A;PIP5K1A | 5'UTR;1stExon;5'UTR;1stExon;5'UTR;1stExon;1stExon;5'UTR | Island |
| cg23117316 | -0.08165078 | 0.017798662 | 7 | PL-5283 | TSS1500 | N_Shore |
| cg24405174 | -0.086144286 | 0.043665871 | 7 | PL-5283 | TSS1500 | N_Shore |
| cg21988244 | -0.085176828 | 0.000562484 | 19 | PNMAL1;PNMAL1 | TSS1500;TSS1500 | Island |
| cg24192846 | -0.093170566 | 4.40E-14 | 6 | PPP1R11 | 3'UTR | N_Shore |
| cg07915318 | -0.087199228 | 4.37E-14 | 12 | PRH1-PRR4;PRH1;PRH1 | Body;Body;5'UTR |  |
| cg06764475 | -0.0974507 | 9.43E-15 | 6 | PRIM2 | Body |  |
| cg17896229 | -0.085734422 | 4.32E-15 | 20 | PROKR2 | 1stExon | N_Shore |
| cg25700305 | 0.081806055 | 6.79E-07 | 1 | PRPF3 | TSS200 | Island |
| cg11893547 | -0.088347814 | 3.83E-10 | 14 | PSMA3;PSMA3;PSMA3 | Body;Body;Body |  |
| cg02646643 | -0.086543887 | 6.73E-16 | 3 | PSMD2 | 3'UTR |  |
| cg03182374 | -0.082186771 | 8.41E-11 | 6 | PTP4A1 | Body |  |
| cg21705926 | -0.086436366 | 0.003087503 | 7 | PTPRN2;PTPRN2;PTPRN2;PTPRN2;PTPRN2 | Body;Body;Body;Body;Body | S_Shore |
| cg11309891 | -0.082257663 | 6.37E-09 | 11 | RAB6A;RAB6A | 3'UTR;3'UTR |  |
| cg14150708 | -0.092674057 | 1.44E-14 | 3 | RAD18 | Body |  |
| cg05465443 | -0.098922155 | 8.93E-11 | 3 | RAD54L2 | Body |  |
| cg03705048 | -0.09577088 | 0.030464323 | 18 | RALBP1 | 5'UTR |  |
| cg17988535 | 0.133931523 | 6.69E-12 | 13 | RASA3 | Body | S_Shore |
| cg27133230 | -0.112268347 | 3.85E-14 | 20 | RBPJL | Body | S_Shore |
| cg23961843 | -0.114997843 | 1.73E-11 | 1 | RGL1;APOBEC4 | 5'UTR;TSS1500 |  |
| cg17350690 | -0.083964092 | 0.000122258 | 15 | RGMA;RGMA;RGMA;RGMA;RGMA;RGMA | Body;Body;Body;Body;Body;Body | Island |
| cg27655494 | -0.098312467 | 1.03E-14 | 1 | RGS7;RGS7;RGS7;RGS7 | Body;Body;Body;Body |  |
| cg08742290 | -0.081610569 | 0.006785006 | 6 | RNASET2 | TSS1500 | S_Shore |
| cg25258033 | -0.098275042 | 0.00113035 | 6 | RNASET2 | Body | N_Shore |
| cg15812759 | -0.081298965 | 4.52E-05 | 5 | RNF14;RNF14;RNF14;RNF14;RNF14;RNF14 | 5'UTR;5'UTR;TSS1500;TSS200;TSS200;TSS200 | N_Shore |
| cg01603311 | -0.13916107 | 1.15E-13 | 17 | RNF157 | Body | N_Shore |
| cg24013599 | -0.081964536 | 4.17E-10 | 8 | RNF19A;RNF19A;RNF19A | 5'UTR;5'UTR;1stExon |  |
| cg17583390 | -0.080903844 | 2.17E-19 | 22 | RNF215 | Body | N_Shelf |
| cg12113395 | -0.115599706 | 2.10E-14 | 3 | RNF7;RNF7;RNF7;RNF7;RNF7 | 3'UTR;3'UTR;3'UTR;Body;Body |  |
| cg01973725 | -0.110928806 | 0.000303478 | 5 | RNU5E;RNU5D;ZCCHC9;ZCCHC9;ZCCHC9 | Body;Body;3'UTR;3'UTR;3'UTR |  |
| cg19963700 | -0.084537918 | 7.09E-11 | 1 | ROR1;ROR1 | Body;Body |  |
| cg18642567 | -0.086064691 | 2.15E-10 | 14 | RPGRIP1 | TSS1500 |  |
| cg04733758 | -0.097680634 | 1.58E-07 | 5 | RPL26L1 | Body |  |
| cg13922921 | -0.080568057 | 7.38E-08 | 19 | RPSAP58 | Body |  |
| cg10475689 | -0.08629878 | 1.44E-14 | 16 | RUNDC2A | Body |  |
| cg10266336 | -0.084717877 | 0.01841101 | 11 | SAA2;SAA2 | TSS200;TSS200 |  |
| ch.8.2353618R | 0.086232962 | 0.008769298 | 8 | SAMD12 | Body |  |
| cg05790002 | -0.110020736 | 2.00E-15 | 16 | SDR42E1 | Body |  |
| cg00832928 | -0.103056885 | 1.99E-15 | 3 | SELT | Body | Island |
| cg06363265 | -0.090794485 | 1.79E-14 | 7 | SEPT7;SEPT7;SEPT7 | Body;Body;Body |  |
| cg20443413 | -0.088531545 | 1.75E-11 | 6 | SGK1 | Body |  |
| cg14783439 | 0.086523971 | 9.90E-06 | 19 | SH3GL1;SH3GL1;SH3GL1 | 1stExon;1stExon;1stExon | Island |
| cg18440889 | -0.099390297 | 9.45E-16 | 11 | SHANK2 | Body |  |
| cg11320438 | -0.080708489 | 1.35E-11 | 19 | SIGLEC11;SIGLEC11 | Body;Body |  |
| cg15956162 | -0.082093227 | 1.44E-11 | 14 | SIP1;SIP1;SIP1 | Body;3'UTR;Body |  |
| cg11986223 | 0.090302034 | 9.27E-06 | 7 | SKAP2 | Body |  |
| cg12089827 | -0.080049332 | 5.61E-14 | 5 | SLC35A4;APBB3;APBB3;APBB3;APBB3 | 5'UTR;TSS1500;TSS1500;TSS1500;TSS1500 | S_Shore |
| cg10369197 | -0.106964602 | 4.26E-08 | 17 | SLC39A11;SLC39A11 | Body;Body |  |
| cg10514580 | -0.117821055 | 0.00016284 | 17 | SLFN12;SLFN12 | TSS1500;TSS200 |  |
| cg20746451 | 0.110213462 | 0.014076575 | 4 | SMARCAD1;SMARCAD1;SMARCAD1;LOC101929210 | TSS1500;TSS1500;TSS1500;Body | N_Shore |
| cg06205244 | -0.104644648 | 1.03E-12 | 6 | SNX14;SNX14;SNX14;SNX14 | 5'UTR;Body;Body;Body | N_Shelf |
| cg23509861 | -0.091036731 | 1.92E-08 | 5 | SNX2;SNX2 | TSS1500;TSS1500 | N_Shore |
| cg03456799 | -0.083018756 | 0.005343769 | 1 | SNX27 | Body |  |
| cg21610999 | -0.103374832 | 2.10E-14 | 17 | SPDYE4 | TSS200 |  |
| cg16677432 | -0.126352382 | 0.037266565 | 16 | SPG7 | Body |  |
| cg01011918 | -0.098999136 | 9.50E-15 | 1 | SPRR2D;SPRR2D | 1stExon;5'UTR |  |
| cg09309707 | -0.113708313 | 0.025226555 | 3 | SPSB4 | Body | S_Shelf |
| cg21531080 | -0.08480926 | 2.19E-10 | 1 | SPTA1 | Body |  |
| cg01993027 | -0.084764216 | 9.10E-10 | 1 | SRSF4 | Body |  |
| cg00389672 | -0.085052368 | 9.60E-14 | 17 | SSH2;SSH2;SSH2 | Body;Body;Body |  |
| cg12601237 | -0.0991237 | 5.31E-16 | 10 | ST8SIA6-AS1;ST8SIA6 | TSS1500;Body |  |
| cg00047657 | -0.081359449 | 2.51E-13 | 10 | STOX1;STOX1;STOX1;STOX1 | Body;Body;Body;Body |  |
| cg06873352 | -0.089979303 | 0.011300507 | 17 | STRADA;STRADA;STRADA;STRADA;STRADA;STRADA | TSS1500;TSS1500;TSS1500;TSS1500;TSS1500;TSS1500 | S_Shore |
| cg02718464 | -0.103217141 | 2.40E-09 | 13 | SUGT1L1;SLC25A15;MIR621 | Body;3'UTR;TSS1500 |  |
| cg15300875 | -0.08780147 | 1.20E-10 | 7 | SUMF2;SUMF2;SUMF2;SUMF2;SUMF2;CCT6A;CCT6A | TSS1500;TSS1500;TSS1500;TSS1500;TSS1500;3'UTR;3'UTR | N_Shore |
| cg15201783 | -0.087127205 | 4.12E-09 | 14 | SUSD6 | 3'UTR |  |
| cg16447252 | -0.081036282 | 8.44E-13 | 10 | SVIL-AS1;SVIL;SVIL | Body;Body;Body |  |
| cg05342835 | -0.080050918 | 0.013254402 | 1 | SYNC;SYNC | Body;Body |  |
| cg14079545 | -0.133850333 | 1.79E-14 | 6 | SYNGAP1 | Body | N_Shelf |
| cg11340603 | -0.084115861 | 0.029612542 | 12 | SYT1;SYT1;SYT1 | Body;Body;Body |  |
| cg15810754 | -0.084858931 | 1.90E-05 | 15 | TJP1;TJP1;TJP1;TJP1 | Body;Body;Body;Body |  |
| cg24299813 | -0.08425952 | 1.01E-07 | 11 | TMEM109 | 5'UTR | S_Shelf |
| cg08713543 | -0.093377251 | 2.50E-13 | 16 | TMEM231;TMEM231;TMEM231 | Body;Body;Body |  |
| cg17285822 | -0.091013661 | 3.62E-13 | 9 | TMEM245 | TSS1500 | S_Shore |
| cg23782308 | -0.086649155 | 0.020035358 | 4 | TNIP3;TNIP3;TNIP3 | TSS200;Body;Body |  |
| cg11963436 | -0.105708427 | 1.95E-05 | 10 | TNKS2 | Body |  |
| cg21210642 | -0.084465782 | 0.024833735 | 9 | TRIM14;TRIM14;TRIM14;TRIM14 | TSS1500;TSS1500;TSS1500;TSS1500 | S_Shore |
| cg23024233 | -0.127052553 | 3.21E-15 | 7 | TRIM4;TRIM4 | Body;Body |  |
| cg26118675 | -0.092097935 | 6.48E-10 | 2 | TRIM43 | TSS1500 |  |
| cg00793774 | 0.080441129 | 0.042677504 | 10 | TRIM8 | TSS1500 | Island |
| cg08399134 | 0.089584418 | 4.31E-06 | 6 | TRMT11 | Body |  |
| cg01291513 | -0.117355521 | 2.92E-10 | 3 | TSEN2;TSEN2;TSEN2;TSEN2;TSEN2;TSEN2;TSEN2;TSEN2;TSEN2 | ExonBnd;ExonBnd;ExonBnd;ExonBnd;Body;Body;Body;Body;Body |  |
| cg13405878 | -0.109366967 | 0.035985399 | 11 | TSKU;TSKU | 5'UTR;5'UTR |  |
| cg00622907 | -0.092195967 | 1.04E-16 | 8 | TSNARE1 | Body | N_Shelf |
| cg06276780 | -0.080239139 | 3.69E-14 | 8 | UBE2V2 | Body |  |
| cg08797194 | -0.096292392 | 5.39E-05 | 13 | UGGT2 | Body | N_Shore |
| cg01380308 | -0.080349562 | 0.009360483 | 10 | UNC5B;UNC5B | Body;Body |  |
| cg22995176 | 0.094219806 | 0.007650841 | 7 | UPK3B;UPK3B | TSS1500;TSS1500 |  |
| cg07046674 | 0.095877551 | 0.008420584 | 8 | VPS13B;VPS13B | Body;Body |  |
| cg04667820 | -0.082479944 | 2.34E-14 | 2 | VPS54;VPS54 | 5'UTR;5'UTR |  |
| cg12187850 | -0.12940508 | 6.30E-16 | 5 | WDR70 | Body |  |
| cg15137408 | -0.081822048 | 0.000309307 | 15 | WDR73;WDR73;WDR73;WDR73;WDR73 | Body;Body;Body;Body;Body |  |
| cg25554036 | -0.08209353 | 1.42E-07 | 4 | WFS1;WFS1 | TSS1500;TSS1500 | N_Shore |
| cg00322003 | -0.105099189 | 0.01788496 | 12 | WNK1;WNK1;WNK1;WNK1 | Body;Body;Body;Body |  |
| cg07744399 | -0.106060108 | 8.87E-09 | 7 | XRCC2 | Body |  |
| cg22726533 | -0.100100974 | 5.07E-12 | 7 | YAE1D1;YAE1D1 | Body;Body | S_Shelf |
| cg26929161 | -0.100077668 | 7.95E-17 | 20 | ZFP64 | Body | Island |
| cg19729503 | -0.097926769 | 4.54E-13 | 20 | ZHX3 | 5'UTR |  |
| cg19933320 | -0.098655619 | 9.71E-16 | 7 | ZNF107;ZNF107 | TSS1500;TSS1500 | N_Shore |
| cg00764514 | -0.08325774 | 4.07E-07 | 7 | ZNF138;ZNF138;ZNF138;ZNF138 | TSS1500;TSS1500;TSS1500;TSS1500 | N_Shore |
| cg25477163 | -0.100266726 | 3.14E-14 | 19 | ZNF211;ZNF211 | Body;Body | S_Shelf |
| cg05341610 | -0.094737805 | 4.12E-07 | 19 | ZNF264 | TSS1500 | N_Shore |
| cg07369507 | 0.088863582 | 0.018488128 | 6 | ZNF323;ZNF323;ZNF323;ZNF323;ZNF323;ZNF323 | TSS1500;TSS200;5'UTR;TSS200;5'UTR;TSS1500 |  |
| cg27224816 | -0.086898387 | 1.09E-05 | 20 | ZNF337-AS1;ZNF337-AS1;ZNF337-AS1 | Body;Body;Body |  |
| cg09314196 | -0.096869084 | 0.043979275 | 19 | ZNF492 | TSS1500 | N_Shore |
| cg17751872 | -0.084347512 | 0.034925632 | 19 | ZNF714;ZNF714 | 5'UTR;1stExon | N_Shore |
| cg25790241 | -0.080418938 | 2.16E-16 | 19 | ZNF808 | Body |  |
| cg18266588 | 0.200986239 | 0.049001861 | 2 |  |  |  |
| cg20934427 | 0.153855434 | 0.034003782 | 8 |  |  |  |
| cg04402345 | 0.122010713 | 0.049642664 | 5 |  |  |  |
| cg27245056 | 0.107502011 | 0.011459871 | 10 |  |  |  |
| cg17537073 | 0.099901764 | 0.040804039 | 13 |  |  | Island |
| cg19600494 | 0.092862056 | 5.94E-06 | 2 |  |  | Island |
| cg11254578 | 0.090816349 | 0.042541922 | 11 |  |  |  |
| cg24566341 | 0.086411974 | 1.72E-07 | 7 |  |  |  |
| cg01176670 | 0.08495968 | 0.001584668 | 6 |  |  |  |
| cg08821715 | 0.081831716 | 1.65E-05 | 10 |  |  | N_Shore |
| cg00336022 | 0.080113056 | 1.48E-05 | 16 |  |  |  |
| cg09415969 | -0.080242348 | 2.89E-10 | 19 |  |  |  |
| cg15395345 | -0.080507086 | 4.30E-09 | 8 |  |  | N_Shelf |
| cg25743234 | -0.080556303 | 1.91E-15 | 12 |  |  |  |
| cg14223654 | -0.080778082 | 0.039376405 | 2 |  |  | Island |
| cg22797297 | -0.080820281 | 0.010399796 | 13 |  |  |  |
| cg05162858 | -0.081180026 | 0.00752844 | 10 |  |  |  |
| cg11735780 | -0.08123358 | 0.01757568 | 3 |  |  |  |
| cg08077014 | -0.081256365 | 2.63E-08 | 10 |  |  |  |
| cg12719319 | -0.081463026 | 4.95E-13 | 19 |  |  | S_Shore |
| cg07283849 | -0.081499695 | 0.04655761 | 7 |  |  |  |
| cg14894801 | -0.081564514 | 5.51E-16 | 10 |  |  |  |
| cg13494229 | -0.081604391 | 0.000657473 | 13 |  |  |  |
| cg27521356 | -0.081659701 | 4.76E-11 | 22 |  |  |  |
| cg02738677 | -0.081746385 | 9.96E-06 | 3 |  |  |  |
| cg17321607 | -0.081834306 | 4.51E-15 | 17 |  |  |  |
| cg07611721 | -0.081877305 | 5.62E-10 | 5 |  |  |  |
| cg11784147 | -0.081908597 | 1.81E-11 | 20 |  |  |  |
| cg06190672 | -0.081915023 | 2.92E-08 | 20 |  |  |  |
| cg22805491 | -0.082116797 | 1.39E-13 | 14 |  |  |  |
| cg23403031 | -0.082121298 | 0.017962709 | 17 |  |  |  |
| cg03008165 | -0.082143267 | 7.39E-14 | 11 |  |  |  |
| cg26551069 | -0.08221141 | 1.01E-12 | 11 |  |  |  |
| cg17424859 | -0.08223846 | 6.48E-13 | 6 |  |  |  |
| cg08450091 | -0.082279408 | 9.48E-05 | 3 |  |  | Island |
| cg05754259 | -0.082303582 | 6.27E-12 | 6 |  |  |  |
| cg02595959 | -0.082308831 | 4.19E-05 | 6 |  |  |  |
| cg04036182 | -0.082419399 | 0.014134869 | 15 |  |  | Island |
| cg09235723 | -0.082437542 | 4.68E-13 | 11 |  |  |  |
| cg15227633 | -0.082458968 | 1.53E-07 | 6 |  |  |  |
| cg14879127 | -0.082541426 | 1.37E-14 | 15 |  |  |  |
| cg09598512 | -0.082684842 | 6.25E-13 | 3 |  |  |  |
| cg09390058 | -0.082701635 | 2.67E-07 | 17 |  |  | S_Shore |
| cg14052514 | -0.082704494 | 0.046639336 | 2 |  |  |  |
| cg00186563 | -0.082752767 | 0.033759107 | 11 |  |  |  |
| cg08213792 | -0.082849297 | 1.14E-09 | 16 |  |  |  |
| cg05521474 | -0.082953363 | 5.51E-06 | 5 |  |  |  |
| cg10087939 | -0.083118065 | 8.59E-10 | 3 |  |  |  |
| cg23047033 | -0.083185145 | 2.34E-11 | 7 |  |  |  |
| cg06936108 | -0.083264236 | 1.22E-11 | 17 |  |  | S_Shore |
| cg02663382 | -0.083298808 | 0.00038827 | 19 |  |  | Island |
| cg25949447 | -0.083416645 | 9.57E-06 | 2 |  |  | N_Shelf |
| cg22871949 | -0.083487716 | 0.000804835 | 7 |  |  | Island |
| cg25064860 | -0.083518986 | 0.02324446 | 1 |  |  |  |
| cg24609094 | -0.083610282 | 8.89E-12 | 13 |  |  | Island |
| cg17480781 | -0.08375933 | 5.93E-11 | 9 |  |  |  |
| cg05460746 | -0.084139085 | 0.004588829 | 8 |  |  |  |
| cg06028934 | -0.08422802 | 1.75E-11 | 1 |  |  |  |
| cg05742286 | -0.084337358 | 2.05E-17 | 12 |  |  | Island |
| cg16058452 | -0.084359276 | 2.43E-09 | 15 |  |  |  |
| cg21357536 | -0.084394887 | 4.47E-11 | 15 |  |  | S_Shelf |
| cg18593556 | -0.084714111 | 1.41E-10 | 20 |  |  |  |
| cg08769199 | -0.08473723 | 4.90E-05 | 11 |  |  |  |
| cg25308322 | -0.084972184 | 0.037507322 | 13 |  |  |  |
| cg06919059 | -0.084976699 | 9.78E-06 | 1 |  |  |  |
| cg14436168 | -0.085435958 | 4.73E-05 | 4 |  |  |  |
| cg00305263 | -0.085604231 | 1.20E-10 | 1 |  |  |  |
| cg13986667 | -0.085735904 | 1.03E-15 | 10 |  |  |  |
| cg09789874 | -0.085844604 | 1.94E-05 | 12 |  |  |  |
| cg00108228 | -0.085901877 | 0.007498837 | 8 |  |  |  |
| cg26174329 | -0.085974736 | 1.32E-09 | 7 |  |  | N_Shelf |
| cg21744189 | -0.086385116 | 0.04864714 | 8 |  |  |  |
| cg06153448 | -0.086534125 | 0.030900392 | 2 |  |  |  |
| cg22728268 | -0.086709144 | 3.66E-07 | 3 |  |  |  |
| cg12038352 | -0.086784763 | 0.000231647 | 19 |  |  |  |
| cg11558106 | -0.086990009 | 6.51E-17 | 7 |  |  |  |
| cg05900177 | -0.087133524 | 2.68E-08 | 4 |  |  |  |
| cg12568217 | -0.087250313 | 0.00320779 | 3 |  |  |  |
| cg19546536 | -0.087336191 | 8.10E-10 | 5 |  |  |  |
| cg06763665 | -0.087356963 | 3.59E-07 | 1 |  |  |  |
| cg02685684 | -0.08741287 | 8.96E-14 | 7 |  |  |  |
| cg21054919 | -0.087563296 | 0.002395575 | 2 |  |  | Island |
| cg17678740 | -0.088041719 | 5.66E-13 | 7 |  |  | N_Shore |
| cg00316520 | -0.08813965 | 4.80E-09 | 3 |  |  | N_Shelf |
| cg05457480 | -0.08845258 | 0.017355333 | 5 |  |  | Island |
| cg24773848 | -0.088505952 | 4.05E-09 | 2 |  |  |  |
| cg23129217 | -0.088552593 | 1.45E-09 | 12 |  |  | S_Shore |
| cg07516076 | -0.088602833 | 1.03E-13 | 7 |  |  | Island |
| cg08609238 | -0.088651281 | 1.37E-14 | 9 |  |  | Island |
| cg26488405 | -0.088971507 | 1.26E-10 | 5 |  |  | N_Shore |
| cg15310722 | -0.089185171 | 2.05E-17 | 17 |  |  |  |
| cg21399992 | -0.089686443 | 2.88E-05 | 7 |  |  |  |
| cg07303975 | -0.090041477 | 3.72E-09 | 10 |  |  |  |
| cg22221266 | -0.090405747 | 5.08E-11 | 14 |  |  |  |
| cg04791162 | -0.090496944 | 7.63E-12 | 14 |  |  | Island |
| cg14380045 | -0.090553076 | 7.97E-16 | 9 |  |  | Island |
| cg12299494 | -0.090606694 | 0.018078481 | 3 |  |  |  |
| cg12223562 | -0.090609825 | 8.73E-08 | 2 |  |  |  |
| cg09048334 | -0.090884142 | 0.003523743 | 6 |  |  | Island |
| cg15050398 | -0.090895946 | 1.36E-12 | 6 |  |  | N_Shelf |
| cg07791468 | -0.090919959 | 6.59E-14 | 2 |  |  |  |
| cg03178542 | -0.091716094 | 1.42E-14 | 10 |  |  |  |
| cg19564603 | -0.092347329 | 6.51E-17 | 7 |  |  |  |
| cg06765686 | -0.092572743 | 1.10E-12 | 16 |  |  | N_Shelf |
| cg09210005 | -0.092593103 | 2.78E-16 | 20 |  |  |  |
| cg01343093 | -0.092792971 | 5.30E-13 | 7 |  |  |  |
| cg01789846 | -0.093235637 | 1.52E-12 | 4 |  |  | Island |
| cg11002430 | -0.093496526 | 7.62E-05 | 1 |  |  | S_Shore |
| cg20653227 | -0.093706739 | 1.64E-14 | 5 |  |  |  |
| cg19183395 | -0.093728938 | 0.014645452 | 2 |  |  | Island |
| cg03543603 | -0.093763349 | 0.028213755 | 5 |  |  | S_Shelf |
| cg01741372 | -0.09397352 | 0.018836759 | 11 |  |  |  |
| cg05466248 | -0.094005797 | 1.49E-11 | 14 |  |  |  |
| cg14661028 | -0.094214646 | 1.22E-14 | 1 |  |  | S_Shore |
| cg08876558 | -0.094685956 | 1.23E-10 | 11 |  |  |  |
| cg13082173 | -0.094995828 | 0.046377559 | 10 |  |  |  |
| cg11361809 | -0.095213415 | 0.021572419 | 20 |  |  |  |
| cg06507271 | -0.095590037 | 8.01E-10 | 7 |  |  |  |
| cg04438374 | -0.095639384 | 2.63E-06 | 7 |  |  |  |
| cg19500764 | -0.096304763 | 0.012748269 | 2 |  |  |  |
| cg04299067 | -0.096353658 | 0.02395441 | 4 |  |  |  |
| cg19751754 | -0.097315497 | 8.47E-12 | 17 |  |  |  |
| cg25444615 | -0.097742801 | 3.61E-13 | 19 |  |  |  |
| cg14341057 | -0.09791827 | 1.48E-12 | 13 |  |  |  |
| cg06061636 | -0.097943164 | 1.44E-08 | 5 |  |  |  |
| cg20368377 | -0.098011529 | 4.46E-10 | 13 |  |  | N_Shore |
| cg04611493 | -0.098246175 | 1.22E-13 | 2 |  |  |  |
| cg07575344 | -0.098827149 | 2.93E-13 | 1 |  |  | N_Shelf |
| cg26873329 | -0.099262076 | 2.77E-11 | 5 |  |  |  |
| cg11753711 | -0.09937738 | 1.87E-05 | 7 |  |  |  |
| cg13356164 | -0.099434877 | 1.04E-06 | 15 |  |  |  |
| cg09427936 | -0.09952437 | 9.65E-16 | 1 |  |  |  |
| cg09651654 | -0.099579528 | 0.023260157 | 12 |  |  | S_Shore |
| cg14405103 | -0.100696257 | 0.004466768 | 5 |  |  |  |
| cg03285130 | -0.100805482 | 4.92E-14 | 10 |  |  |  |
| cg02593636 | -0.101024682 | 1.00E-09 | 12 |  |  |  |
| cg10471743 | -0.101181831 | 1.30E-06 | 5 |  |  | S_Shore |
| cg16471739 | -0.101552743 | 0.024214726 | 2 |  |  |  |
| cg04375046 | -0.101745607 | 0.014620052 | 5 |  |  |  |
| cg07538465 | -0.101867577 | 2.27E-16 | 12 |  |  |  |
| cg13881513 | -0.10187593 | 6.96E-09 | 18 |  |  | N_Shelf |
| cg06888084 | -0.102870129 | 2.91E-11 | 11 |  |  |  |
| cg17784744 | -0.103076812 | 4.94E-11 | 17 |  |  |  |
| cg15123819 | -0.103779122 | 1.62E-10 | 5 |  |  |  |
| cg25828445 | -0.10431 | 0.048708476 | 12 |  |  | S_Shore |
| cg05909891 | -0.104321075 | 7.40E-13 | 12 |  |  |  |
| cg17616250 | -0.104965301 | 0.04269962 | 6 |  |  |  |
| cg16919064 | -0.106273679 | 0.032585125 | 2 |  |  |  |
| cg24568548 | -0.106456584 | 1.02E-10 | 11 |  |  | S_Shore |
| cg21653586 | -0.107955889 | 3.60E-09 | 11 |  |  |  |
| cg12615557 | -0.10812882 | 4.68E-13 | 1 |  |  |  |
| cg02466788 | -0.108263334 | 0.012717676 | 4 |  |  |  |
| cg14298830 | -0.109986499 | 0.040718994 | 20 |  |  |  |
| cg25374269 | -0.111242678 | 0.041305243 | 13 |  |  |  |
| cg21374546 | -0.112850693 | 2.77E-12 | 10 |  |  |  |
| cg13451619 | -0.113448001 | 0.014548279 | 2 |  |  |  |
| cg09133791 | -0.113881748 | 6.58E-12 | 3 |  |  |  |
| cg23244452 | -0.113968095 | 1.64E-14 | 4 |  |  |  |
| cg04627621 | -0.115029121 | 0.011582631 | 17 |  |  |  |
| cg26769184 | -0.115766296 | 6.27E-14 | 22 |  |  |  |
| cg24713959 | -0.117690615 | 0.047941051 | 12 |  |  |  |
| cg08653580 | -0.119943565 | 4.92E-16 | 11 |  |  |  |
| cg07625092 | -0.124922737 | 1.05E-15 | 15 |  |  |  |
| cg15882280 | -0.126963253 | 6.51E-17 | 22 |  |  |  |
| cg27478848 | -0.127711875 | 3.50E-05 | 15 |  |  | S_Shore |
| cg14265604 | -0.12883906 | 2.05E-17 | 1 |  |  |  |
| cg08323960 | -0.137345821 | 1.91E-13 | 10 |  |  |  |
| cg01701555 | -0.137870658 | 8.85E-08 | 1 |  |  | Island |
| cg21922468 | -0.178598078 | 0.01040465 | 19 |  |  | Island |
| cg14502545 | -0.185172051 | 0.034983649 | 4 |  |  |  |
| cg17429539 | -0.229080953 | 0.019678728 | 6 |  |  |  |
| cg13102742 | -0.236191531 | 0.022339482 | 17 |  |  |  |
| cg12957945 | -0.254156842 | 0.015786302 | 2 |  |  |  |

### Supplementary Table S4-b.

**The differentially methylated probes between individuals with PRS and controls.**

| **Probe ID** | **Delta Beta** | **Adjust *P* value** | **chromosome** | **Gene Name** | **Gene Context** | **CpG Island** |
| --- | --- | --- | --- | --- | --- | --- |
| **cg19476594** | -0.100055457 | 1.77E-14 | 1 | ABL2;ABL2;ABL2;ABL2;ABL2;ABL2;ABL2;ABL2 | Body;Body;Body;Body;Body;Body;Body;Body |  |
| **cg11607876** | -0.132309308 | 0.039601554 | 3 | ADAMTS9-AS1;ADAMTS9-AS1;ADAMTS9 | Body;Body;Body |  |
| **cg10297223** | -0.140344732 | 1.80E-16 | 3 | AGTR1;AGTR1;AGTR1;AGTR1 | TSS1500;TSS1500;TSS1500;TSS1500 | N_Shore |
| **cg15004555** | -0.089550348 | 2.85E-13 | 1 | AIM2 | Body |  |
| **cg12316181** | -0.082559997 | 2.95E-08 | 13 | ANKRD20A9P | Body |  |
| **cg16269379** | -0.083484113 | 3.77E-09 | 12 | ANP32D | TSS200 |  |
| **cg20592391** | -0.09477514 | 0.014287396 | 15 | AP3B2 | Body |  |
| **cg10493270** | -0.080476319 | 1.21E-10 | 11 | ARHGEF12 | Body |  |
| **cg00106757** | -0.10336757 | 0.039886094 | 16 | ATF7IP2;ATF7IP2;ATF7IP2;ATF7IP2 | Body;Body;Body;Body |  |
| **cg26840922** | -0.091195405 | 1.98E-16 | 3 | ATP1B3 | Body |  |
| **cg12953621** | -0.088639899 | 1.07E-15 | 12 | ATP5G2;ATP5G2 | Body;Body |  |
| **cg14922317** | 0.09337593 | 1.12E-07 | 16 | ATXN2L;ATXN2L;ATXN2L;ATXN2L;ATXN2L;ATXN2L;ATXN2L;ATXN2L;ATXN2L;ATXN2L;ATXN2L;ATXN2L;ATXN2L;ATXN2L | 5'UTR;5'UTR;5'UTR;5'UTR;5'UTR;5'UTR;5'UTR;1stExon;1stExon;1stExon;1stExon;1stExon;1stExon;1stExon | Island |
| **cg23097266** | -0.180337047 | 0.03026641 | 9 | BICD2;BICD2 | Body;Body |  |
| **cg10795676** | -0.082870334 | 1.51E-16 | 6 | BTN3A2 | Body |  |
| **cg13961165** | -0.107331737 | 7.03E-16 | 2 | C2orf27A;C2orf27A | 1stExon;5'UTR |  |
| **cg23237765** | -0.088081327 | 0.000206717 | 7 | C7orf20 | Body | S_Shelf |
| **cg13116816** | -0.081943493 | 8.47E-09 | 7 | C7orf23 | Body |  |
| **cg04038932** | 0.092801792 | 1.54E-08 | 9 | C9orf171 | Body | S_Shore |
| **cg04158140** | -0.090415497 | 3.02E-14 | 7 | CALN1;CALN1 | Body;Body |  |
| **cg22852177** | -0.088010595 | 1.54E-06 | 15 | CASC5;CASC5 | Body;Body |  |
| **cg02941728** | -0.097724423 | 3.61E-15 | 11 | CCDC86 | 3'UTR | N_Shelf |
| **cg03960701** | -0.107890602 | 8.97E-18 | 7 | CCT6A;CCT6A;SNORA15 | Body;Body;TSS1500 |  |
| **cg05465968** | -0.123952362 | 0.020731382 | 6 | CDKAL1 | Body |  |
| **cg08152083** | -0.08449889 | 1.96E-10 | 17 | CDRT15L2 | TSS1500 |  |
| **cg03388324** | -0.100890462 | 3.56E-19 | 6 | CEP85L;CEP85L;CEP85L | Body;Body;Body |  |
| **cg02301920** | -0.112326146 | 0.017106509 | 16 | CES1;CES1;CES1 | Body;Body;Body | Island |
| **cg01166932** | -0.111562811 | 2.56E-05 | 15 | CGNL1;CGNL1 | 5'UTR;5'UTR |  |
| **cg12451061** | -0.095504319 | 0.041363878 | 12 | CISTR | Body | S_Shelf |
| **cg10690677** | -0.084203051 | 2.11E-14 | 1 | CLCA4;CLCA4 | Body;Body | Island |
| **cg10182306** | -0.086947072 | 1.13E-10 | 1 | CLCNKA;CLCNKA;CLCNKA | Body;Body;Body |  |
| **cg01304461** | 0.089644593 | 4.60E-09 | 12 | CLEC12A | TSS1500 |  |
| **cg10967735** | -0.084681844 | 2.26E-13 | 10 | CPEB3 | Body |  |
| **cg21985184** | -0.122103126 | 0.019900873 | 7 | CREB3L2 | 3'UTR |  |
| **cg22682254** | -0.082920415 | 2.04E-09 | 17 | CRYBA1 | TSS1500 |  |
| **cg04773134** | -0.088613869 | 0.037127514 | 16 | CTU2;CTU2 | Body;Body | N_Shelf |
| **cg08228804** | -0.081273812 | 1.52E-13 | 11 | CYP2R1 | TSS1500 | S_Shore |
| **cg24655310** | -0.103296524 | 0.00080019 | 19 | CYP4F11;CYP4F11 | 1stExon;Body |  |
| **cg03278573** | -0.098801829 | 5.03E-12 | 5 | DAP;DAP | Body;Body |  |
| **cg00574307** | -0.098953029 | 1.71E-14 | 13 | DCT;DCT | Body;Body |  |
| **cg07341559** | -0.112565549 | 1.80E-16 | 11 | DEFB108B | Body |  |
| **cg06012621** | -0.205062178 | 0.006756781 | 2 | DIRC3 | Body |  |
| **cg07068730** | -0.080461562 | 2.45E-08 | 3 | DLG1;DLG1;DLG1;DLG1 | Body;Body;Body;Body |  |
| **cg06871323** | -0.095674554 | 1.24E-11 | 7 | DPY19L1 | Body |  |
| **cg00218840** | -0.097389953 | 1.00E-13 | 1 | DUSP5P1;RHOU | Body;Body | S_Shore |
| **cg02449698** | -0.097466296 | 1.63E-16 | 1 | EFCAB7;DLEU2L | Body;TSS200 |  |
| **cg18803079** | -0.133651124 | 3.56E-19 | 1 | EFCAB7;DLEU2L | Body;TSS200 |  |
| **cg13648157** | -0.083033156 | 6.73E-07 | 5 | EFCAB9 | TSS1500 |  |
| **cg18815120** | -0.108419233 | 0.027849885 | 1 | EGLN1 | Body |  |
| **cg17241224** | -0.080146827 | 2.05E-10 | 2 | EHBP1;EHBP1;EHBP1;EHBP1 | Body;Body;Body;Body |  |
| **cg06362582** | 0.282037966 | 0.00092211 | 7 | ELMO1;ELMO1;ELMO1 | Body;Body;Body |  |
| **cg13442241** | -0.096080271 | 4.29E-16 | 8 | EMC2 | Body |  |
| **cg06658698** | -0.115963382 | 7.39E-08 | 12 | EPS8 | Body |  |
| **cg00084591** | -0.101034146 | 3.95E-15 | 9 | ERP44 | Body |  |
| **cg18194842** | -0.086065316 | 4.64E-15 | 1 | EYA3;EYA3;EYA3;EYA3;EYA3 | Body;5'UTR;5'UTR;5'UTR;5'UTR |  |
| **cg06051442** | -0.092518373 | 3.61E-13 | 9 | FAM120A;FAM120A;FAM120A;FAM120A | Body;Body;Body;Body | S_Shore |
| **cg14115740** | -0.115739466 | 2.25E-15 | 9 | FANCC | 5'UTR | Island |
| **cg25691044** | -0.090351659 | 1.85E-15 | 8 | FBXO25;FBXO25;FBXO25 | 5'UTR;5'UTR;5'UTR |  |
| **cg18370902** | -0.093058856 | 2.60E-15 | 8 | FBXO25;FBXO25;FBXO25 | 5'UTR;Body;Body |  |
| **cg11000420** | -0.088154761 | 9.92E-05 | 9 | GCNT1;GCNT1;GCNT1;GCNT1 | TSS1500;5'UTR;5'UTR;5'UTR |  |
| **cg00872998** | -0.097237429 | 1.21E-14 | 9 | GDA;GDA;GDA;GDA | Body;Body;Body;Body |  |
| **cg09622330** | -0.089125359 | 1.62E-16 | 16 | GRIN2A;GRIN2A;GRIN2A | Body;Body;Body |  |
| **cg00036352** | 0.129602437 | 0.014210995 | 8 | GSDMD | 5'UTR | S_Shore |
| **cg20583945** | 0.126504136 | 0.014937082 | 8 | GSDMD | 5'UTR | S_Shore |
| **cg15242686** | -0.146144447 | 0.037381569 | 22 | GSTTP1 | TSS1500 |  |
| **cg11141652** | -0.193342592 | 0.044717209 | 22 | GSTTP1 | TSS1500 |  |
| **cg01309418** | -0.09369689 | 2.61E-11 | 6 | GTF3C6 | TSS1500 | N_Shore |
| **cg10342490** | -0.090479249 | 3.35E-13 | 11 | GUCY1A2 | Body |  |
| **cg03289961** | -0.090120622 | 4.94E-06 | 2 | HADHB;HADHB;HADHB;HADHB;HADHB | ExonBnd;ExonBnd;Body;Body;Body |  |
| **cg03876548** | -0.15854755 | 0.011045741 | 6 | HCG4P6 | TSS200 | N_Shore |
| **cg02616069** | -0.081955201 | 4.39E-10 | 7 | HECW1 | Body |  |
| **cg08782481** | 0.089557052 | 4.45E-18 | 6 | HIST1H3I | 1stExon | Island |
| **cg04628742** | -0.108989623 | 1.13E-06 | 6 | HLA-J;NCRNA00171 | TSS1500;Body | N_Shore |
| **cg14750231** | -0.115564137 | 2.02E-05 | 1 | HORMAD1;HORMAD1 | Body;Body |  |
| **cg11962649** | -0.095099665 | 3.06E-11 | 9 | HSPA5 | Body | N_Shore |
| **cg02143429** | -0.086004134 | 6.04E-11 | 15 | ICE2;ICE2 | Body;Body |  |
| **cg07493237** | 0.216471371 | 0.008241829 | 12 | IL26 | TSS1500 |  |
| **cg00470768** | -0.103877017 | 4.34E-13 | 15 | INO80 | Body |  |
| **cg13707371** | -0.088903953 | 0.003553153 | 13 | ITGBL1;ITGBL1;ITGBL1;ITGBL1 | 5'UTR;Body;Body;Body |  |
| **cg17983571** | -0.129713309 | 2.41E-16 | 10 | JMJD1C | Body |  |
| **cg25748121** | -0.095297723 | 1.89E-13 | 18 | KATNAL2 | 5'UTR | N_Shore |
| **cg20762182** | -0.089313066 | 3.96E-18 | 15 | KIF23;KIF23 | Body;Body |  |
| **cg26296769** | -0.109110993 | 9.80E-16 | 6 | KLC4;KLC4;KLC4;MRPL2;MRPL2 | TSS1500;TSS1500;TSS1500;Body;Body | N_Shore |
| **cg14094333** | 0.082108539 | 7.23E-15 | 4 | KLF3;KLF3;FLJ13197 | 1stExon;5'UTR;Body | Island |
| **cg02990553** | -0.09804199 | 5.03E-12 | 17 | KRT28 | 1stExon | S_Shelf |
| **cg04204557** | -0.105946272 | 1.29E-15 | 21 | KRTAP10-11;C21orf29 | TSS1500;Body |  |
| **cg02256315** | -0.094863894 | 1.73E-05 | 11 | LDHAL6A;LDHAL6A | 3'UTR;3'UTR |  |
| **cg21400344** | 0.082480369 | 1.39E-12 | 1 | LDLRAP1;LDLRAP1 | 5'UTR;1stExon | Island |
| **cg00114966** | -0.112900114 | 0.009222021 | 1 | LHX9;LHX9 | Body;Body | S_Shelf |
| **cg07321536** | 0.080083348 | 2.24E-05 | 4 | LIAS;RPL9;LIAS;RPL9 | TSS1500;Body;TSS1500;Body | N_Shore |
| **cg24203469** | -0.08411424 | 2.84E-12 | 7 | LINC01162 | TSS200 |  |
| **cg15049005** | -0.082730717 | 0.022647102 | 15 | LINC01169 | Body | N_Shore |
| **cg14507403** | 0.103644259 | 0.04367744 | 7 | LMTK2 | Body | N_Shelf |
| **cg03264014** | -0.088526111 | 0.042825841 | 19 | LOC100128568;RFX2;RFX2 | Body;Body;Body |  |
| **cg03017824** | -0.10296244 | 1.45E-14 | 1 | LOC100130093;LOC100130093 | TSS1500;TSS1500 | N_Shore |
| **cg04676559** | -0.081954282 | 2.34E-13 | 17 | LOC100131347;PLXDC1 | Body;Body |  |
| **cg09141640** | -0.084054794 | 1.79E-10 | 22 | LOC100506679 | TSS1500 |  |
| **cg23355197** | -0.083522488 | 6.31E-07 | 8 | LOC100506990;LOC100506990 | Body;Body |  |
| **cg13593809** | -0.115922448 | 8.58E-17 | 17 | LOC101559451;PELP1;PELP1 | Body;TSS1500;TSS1500 | S_Shore |
| **cg24674363** | -0.114029617 | 2.44E-11 | 3 | LOC101927056;LOC101927056 | Body;Body |  |
| **cg03472399** | -0.096671242 | 6.04E-14 | 13 | LOC101928697 | Body |  |
| **cg03443822** | -0.120118056 | 0.024931428 | 2 | LOC101929733 | Body |  |
| **cg00137195** | -0.137536862 | 0.049228683 | 9 | LOC286297 | Body |  |
| **cg07240877** | -0.080194793 | 4.56E-10 | 1 | LOC339535 | TSS200 |  |
| **cg06774087** | -0.088777831 | 2.68E-16 | 5 | LOC441089 | Body |  |
| **cg12361190** | -0.091781808 | 5.85E-16 | 6 | LOC441155;EYS;EYS | TSS200;Body;Body | Island |
| **cg11812439** | 0.119024539 | 0.004180319 | 4 | LOC550113;SYT14P1;TMPRSS11F | Body;Body;Body |  |
| **cg24375999** | -0.083689904 | 2.54E-15 | 2 | LOC643387;LOC151174;LOC151174 | Body;TSS1500;TSS1500 | S_Shore |
| **cg21581312** | -0.121402959 | 6.34E-13 | 15 | LOC723972 | TSS200 |  |
| **cg10299917** | -0.095256466 | 1.04E-16 | 22 | LRP5L | 5'UTR |  |
| **cg00586123** | -0.080548481 | 6.02E-05 | 6 | LY86-AS1 | Body |  |
| **cg03900366** | -0.116168183 | 2.23E-16 | 1 | MACF1 | Body |  |
| **cg25960044** | -0.100519809 | 5.12E-09 | 4 | MANBA | Body |  |
| **cg10006801** | -0.102435989 | 3.24E-13 | 17 | MAP2K3;MAP2K3 | Body;Body |  |
| **cg12045156** | -0.085408503 | 7.55E-12 | 17 | MAP2K6 | Body |  |
| **cg01490296** | -0.125246726 | 9.15E-17 | 10 | MCM10;MCM10 | 5'UTR;5'UTR | S_Shore |
| **cg14353218** | -0.085928324 | 7.28E-12 | 14 | MDGA2 | Body |  |
| **cg02141602** | -0.093702052 | 2.11E-10 | 19 | MEIS3;MEIS3;MEIS3 | TSS200;TSS200;TSS200 | S_Shore |
| **cg02173085** | -0.140644631 | 6.35E-17 | 2 | METTL5;METTL5;METTL5 | Body;Body;Body | N_Shelf |
| **cg08477687** | -0.124270197 | 9.22E-05 | 1 | MIR1977 | TSS1500 |  |
| **cg12495798** | -0.094890818 | 0.013134459 | 15 | MIR4313 | TSS200 |  |
| **cg14787112** | -0.093371177 | 1.23E-09 | 19 | MIR517A | TSS1500 |  |
| **cg22431028** | -0.12549776 | 1.45E-16 | 14 | MNAT1;MNAT1 | Body;Body |  |
| **cg07788360** | -0.088373174 | 1.70E-13 | 13 | MRPS31 | Body |  |
| **cg19484299** | -0.080569193 | 1.68E-11 | 2 | MSL3L2 | Body | N_Shore |
| **cg09927022** | -0.090513732 | 1.03E-17 | 6 | MTFR2;MTFR2 | Body;Body |  |
| **cg16074739** | -0.084669196 | 8.00E-08 | 8 | MTUS1;MTUS1;MTUS1;MTUS1 | TSS1500;TSS1500;Body;Body |  |
| **cg07074586** | -0.102825101 | 5.69E-17 | 12 | MYL6;MYL6 | 3'UTR;3'UTR |  |
| **cg03889742** | -0.123602282 | 9.45E-15 | 15 | MYO5A;MYO5A | Body;Body |  |
| **cg08017634** | -0.091461461 | 0.032165711 | 8 | NAPRT1 | Body | Island |
| **cg00500007** | -0.094713817 | 2.55E-07 | 13 | NBEA | Body |  |
| **cg04711526** | -0.12043251 | 9.19E-17 | 14 | NDUFB1 | 3'UTR |  |
| **cg02636488** | 0.095028818 | 0.034444294 | 3 | NEK4 | Body | N_Shore |
| **cg25180916** | 0.080271562 | 0.003990284 | 18 | NFATC1;NFATC1;NFATC1;NFATC1;NFATC1;NFATC1;NFATC1;NFATC1;NFATC1;NFATC1 | 5'UTR;5'UTR;5'UTR;5'UTR;1stExon;1stExon;1stExon;1stExon;1stExon;5'UTR | Island |
| **cg11792281** | -0.129788841 | 8.31E-14 | 17 | NLK | Body |  |
| **cg19645279** | -0.085698023 | 7.35E-13 | 5 | NR3C1;NR3C1;NR3C1;NR3C1;NR3C1;NR3C1;NR3C1;NR3C1;NR3C1;NR3C1;NR3C1;NR3C1;NR3C1;NR3C1;NR3C1 | Body;Body;Body;Body;Body;Body;Body;Body;Body;Body;Body;Body;Body;Body;Body |  |
| **cg13423753** | -0.08083647 | 1.55E-13 | 14 | NRXN3;NRXN3 | Body;Body |  |
| **cg00404641** | -0.089864188 | 0.007889134 | 3 | NUDT16P;NUDT16P | TSS200;TSS1500 | N_Shore |
| **cg21769444** | -0.083547022 | 2.92E-06 | 6 | NUDT3 | TSS1500 | S_Shore |
| **cg22273354** | -0.091845374 | 0.002737024 | 11 | NUP160 | Body |  |
| **cg12949141** | -0.082683444 | 0.000271166 | 5 | PCBD2 | Body |  |
| **cg06649000** | -0.101677699 | 3.93E-16 | 12 | PCBP2;PCBP2;PCBP2;PCBP2;PCBP2;PCBP2;PCBP2 | Body;Body;Body;Body;Body;Body;Body |  |
| **cg22552448** | -0.093851016 | 1.31E-16 | 12 | PCBP2;PCBP2;PCBP2;PCBP2;PCBP2;PCBP2;PCBP2;PCBP2;PCBP2;PCBP2;PCBP2;PCBP2;PCBP2;PCBP2 | ExonBnd;ExonBnd;ExonBnd;ExonBnd;ExonBnd;ExonBnd;ExonBnd;Body;Body;Body;Body;Body;Body;Body | S_Shelf |
| **cg14102005** | -0.088188813 | 1.85E-12 | 14 | PELI2 | Body |  |
| **cg06936779** | 0.131878233 | 4.67E-12 | 1 | PIP5K1A;PIP5K1A;PIP5K1A;PIP5K1A;PIP5K1A;PIP5K1A;PIP5K1A;PIP5K1A | 5'UTR;1stExon;5'UTR;1stExon;5'UTR;1stExon;1stExon;5'UTR | Island |
| **cg13840476** | -0.10091391 | 0.043276243 | 7 | PKD1L1;C7orf69;C7orf69 | Body;5'UTR;Body |  |
| **cg13954435** | -0.08129967 | 2.04E-10 | 15 | PKM;PKM;PKM;PKM;PKM;PKM;PKM | Body;Body;Body;Body;Body;Body;Body |  |
| **cg15593050** | -0.090471984 | 0.046305502 | 11 | PLEKHA7 | Body |  |
| **cg24192846** | -0.087233239 | 1.56E-15 | 6 | PPP1R11 | 3'UTR | N_Shore |
| **cg14427382** | -0.090240761 | 2.55E-16 | 6 | PPP1R2P1 | Body | N_Shore |
| **cg06764475** | -0.10389269 | 1.64E-16 | 6 | PRIM2 | Body |  |
| **cg05570315** | 0.096042002 | 6.72E-08 | 9 | PRKACG | 1stExon | N_Shore |
| **cg17896229** | -0.083783512 | 2.62E-16 | 20 | PROKR2 | 1stExon | N_Shore |
| **cg25700305** | 0.089232304 | 1.16E-15 | 1 | PRPF3 | TSS200 | Island |
| **cg11893547** | -0.100040526 | 1.98E-16 | 14 | PSMA3;PSMA3;PSMA3 | Body;Body;Body |  |
| **cg03182374** | -0.089962482 | 1.45E-16 | 6 | PTP4A1 | Body |  |
| **cg07335637** | -0.08244178 | 0.002294632 | 1 | PUM1;PUM1 | Body;Body |  |
| **cg26094004** | -0.113668275 | 8.97E-18 | 17 | PYY | 5'UTR | S_Shelf |
| **cg11309891** | -0.091937625 | 7.34E-14 | 11 | RAB6A;RAB6A | 3'UTR;3'UTR |  |
| **cg14150708** | -0.091939955 | 3.62E-15 | 3 | RAD18 | Body |  |
| **cg05521364** | -0.087825374 | 1.61E-07 | 14 | RAD51B;RAD51B;RAD51B | Body;Body;Body |  |
| **cg05465443** | -0.103750409 | 8.97E-18 | 3 | RAD54L2 | Body |  |
| **cg17988535** | 0.10108455 | 3.42E-09 | 13 | RASA3 | Body | S_Shore |
| **cg06832168** | -0.090873672 | 7.43E-07 | 4 | RBM47;RBM47 | 5'UTR;5'UTR |  |
| **cg27133230** | -0.112314229 | 2.66E-15 | 20 | RBPJL | Body | S_Shore |
| **cg09797837** | -0.084416748 | 5.69E-17 | 1 | RCC2;RCC2 | 3'UTR;3'UTR | S_Shore |
| **cg23961843** | -0.085306797 | 1.44E-06 | 1 | RGL1;APOBEC4 | 5'UTR;TSS1500 |  |
| **cg01603311** | -0.150608829 | 2.01E-18 | 17 | RNF157 | Body | N_Shore |
| **cg24013599** | -0.096272778 | 2.69E-14 | 8 | RNF19A;RNF19A;RNF19A | 5'UTR;5'UTR;1stExon |  |
| **cg23031969** | 0.097113288 | 2.27E-09 | 7 | RNF216L;RNF216L;RNF216L | Body;Body;Body | Island |
| **cg16288713** | 0.113666174 | 0.033233012 | 13 | RNF219 | TSS1500 | S_Shore |
| **cg12113395** | -0.11329209 | 1.24E-16 | 3 | RNF7;RNF7;RNF7;RNF7;RNF7 | 3'UTR;3'UTR;3'UTR;Body;Body |  |
| **cg01973725** | -0.103979676 | 8.13E-05 | 5 | RNU5E;RNU5D;ZCCHC9;ZCCHC9;ZCCHC9 | Body;Body;3'UTR;3'UTR;3'UTR |  |
| **cg19963700** | -0.083281438 | 2.42E-15 | 1 | ROR1;ROR1 | Body;Body |  |
| **cg18642567** | -0.09054009 | 4.39E-16 | 14 | RPGRIP1 | TSS1500 |  |
| **cg10440639** | 0.089703938 | 0.045670887 | 17 | RPH3AL | Body | N_Shelf |
| **cg10475689** | -0.082712357 | 1.94E-12 | 16 | RUNDC2A | Body |  |
| **ch.8.2353618R** | 0.093454342 | 2.11E-06 | 8 | SAMD12 | Body |  |
| **cg05790002** | -0.100272481 | 5.65E-14 | 16 | SDR42E1 | Body |  |
| **cg00832928** | -0.109318273 | 6.35E-17 | 3 | SELT | Body | Island |
| **cg06363265** | -0.095685807 | 1.49E-16 | 7 | SEPT7;SEPT7;SEPT7 | Body;Body;Body |  |
| **cg06430632** | -0.08016915 | 1.07E-07 | 6 | SFT2D1 | Body |  |
| **cg20443413** | -0.081388445 | 2.76E-10 | 6 | SGK1 | Body |  |
| **cg14783439** | 0.089644199 | 5.14E-13 | 19 | SH3GL1;SH3GL1;SH3GL1 | 1stExon;1stExon;1stExon | Island |
| **cg18440889** | -0.100093141 | 2.01E-18 | 11 | SHANK2 | Body |  |
| **cg11320438** | -0.082387063 | 3.58E-13 | 19 | SIGLEC11;SIGLEC11 | Body;Body |  |
| **cg11581472** | 0.082905721 | 0.003092571 | 2 | SLC25A12;SLC25A12 | Body;Body |  |
| **cg10369197** | -0.104480609 | 2.29E-14 | 17 | SLC39A11;SLC39A11 | Body;Body |  |
| **cg10514580** | -0.090880774 | 0.000694569 | 17 | SLFN12;SLFN12 | TSS1500;TSS200 |  |
| **cg11855615** | -0.103021228 | 3.03E-09 | 17 | SLFN12L | Body | S_Shelf |
| **cg25953395** | -0.09347589 | 6.45E-14 | 3 | SMARCC1 | Body |  |
| **cg24152845** | -0.081368243 | 1.66E-10 | 15 | SNORD115-1 | TSS1500 |  |
| **cg19556901** | -0.088525976 | 4.59E-15 | 15 | SNORD115-1 | TSS1500 |  |
| **cg06205244** | -0.095389615 | 1.69E-13 | 6 | SNX14;SNX14;SNX14;SNX14 | 5'UTR;Body;Body;Body | N_Shelf |
| **cg21610999** | -0.108998024 | 1.26E-18 | 17 | SPDYE4 | TSS200 |  |
| **cg01011918** | -0.094503827 | 4.17E-15 | 1 | SPRR2D;SPRR2D | 1stExon;5'UTR |  |
| **cg01993027** | -0.095296676 | 1.21E-13 | 1 | SRSF4 | Body |  |
| **cg00389672** | -0.091348885 | 8.97E-18 | 17 | SSH2;SSH2;SSH2 | Body;Body;Body |  |
| **cg12601237** | -0.104305165 | 2.52E-16 | 10 | ST8SIA6-AS1;ST8SIA6 | TSS1500;Body |  |
| **cg12920781** | -0.141585182 | 0.005768449 | 13 | STK24;STK24;STK24 | Body;Body;Body |  |
| **cg00047657** | -0.097741286 | 3.16E-19 | 10 | STOX1;STOX1;STOX1;STOX1 | Body;Body;Body;Body |  |
| **cg02718464** | -0.10596985 | 4.99E-14 | 13 | SUGT1L1;SLC25A15;MIR621 | Body;3'UTR;TSS1500 |  |
| **cg16447252** | -0.084110907 | 1.59E-12 | 10 | SVIL-AS1;SVIL;SVIL | Body;Body;Body |  |
| **cg14079545** | -0.102641605 | 1.41E-10 | 6 | SYNGAP1 | Body | N_Shelf |
| **cg15810754** | -0.109736884 | 4.00E-15 | 15 | TJP1;TJP1;TJP1;TJP1 | Body;Body;Body;Body |  |
| **cg24299813** | -0.081711445 | 3.71E-09 | 11 | TMEM109 | 5'UTR | S_Shelf |
| **cg08713543** | -0.081103515 | 2.99E-10 | 16 | TMEM231;TMEM231;TMEM231 | Body;Body;Body |  |
| **cg17285822** | -0.089787214 | 3.61E-14 | 9 | TMEM245 | TSS1500 | S_Shore |
| **cg23313274** | -0.082920353 | 3.91E-11 | 17 | TMEM98;TMEM98 | 3'UTR;3'UTR |  |
| **cg09971388** | 0.09180113 | 2.62E-10 | 4 | TNIP2;TNIP2;TNIP2 | 1stExon;1stExon;TSS200 | Island |
| **cg11963436** | -0.130673697 | 7.89E-13 | 10 | TNKS2 | Body |  |
| **cg23024233** | -0.114052979 | 3.68E-12 | 7 | TRIM4;TRIM4 | Body;Body |  |
| **cg26118675** | -0.100122913 | 3.28E-13 | 2 | TRIM43 | TSS1500 |  |
| **cg21654379** | -0.081088693 | 1.32E-10 | 3 | TRIM71 | Body | S_Shore |
| **cg01291513** | -0.132823785 | 1.83E-18 | 3 | TSEN2;TSEN2;TSEN2;TSEN2;TSEN2;TSEN2;TSEN2;TSEN2;TSEN2 | ExonBnd;ExonBnd;ExonBnd;ExonBnd;Body;Body;Body;Body;Body |  |
| **cg00622907** | -0.082078812 | 1.52E-13 | 8 | TSNARE1 | Body | N_Shelf |
| **cg06276780** | -0.080154088 | 8.51E-15 | 8 | UBE2V2 | Body |  |
| **cg25971920** | -0.082065316 | 5.85E-14 | 1 | UBQLN4;UBQLN4 | Body;Body |  |
| **cg05970768** | -0.082753236 | 4.15E-16 | 5 | UBTD2 | Body | Island |
| **cg08797194** | -0.092037972 | 1.64E-05 | 13 | UGGT2 | Body | N_Shore |
| **cg04667820** | -0.083081956 | 8.98E-15 | 2 | VPS54;VPS54 | 5'UTR;5'UTR |  |
| **cg12187850** | -0.133354798 | 3.86E-17 | 5 | WDR70 | Body |  |
| **cg23252815** | -0.084754416 | 0.01842668 | 20 | WFDC3;DNTTIP1 | 5'UTR;TSS1500 | N_Shore |
| **cg07744399** | -0.08662144 | 1.03E-08 | 7 | XRCC2 | Body |  |
| **cg22726533** | -0.1058498 | 1.64E-16 | 7 | YAE1D1;YAE1D1 | Body;Body | S_Shelf |
| **cg02466933** | 0.095399907 | 4.00E-10 | 2 | YWHAQ | Body | Island |
| **cg15181003** | -0.08342619 | 3.04E-14 | 13 | ZDHHC20;ZDHHC20;MIPEPP3;ZDHHC20;ZDHHC20 | 3'UTR;3'UTR;Body;Body;Body |  |
| **cg26929161** | -0.091910698 | 7.24E-16 | 20 | ZFP64 | Body | Island |
| **cg19729503** | -0.095308975 | 1.32E-13 | 20 | ZHX3 | 5'UTR |  |
| **cg19933320** | -0.103398658 | 6.09E-16 | 7 | ZNF107;ZNF107 | TSS1500;TSS1500 | N_Shore |
| **cg25477163** | -0.102234351 | 1.12E-15 | 19 | ZNF211;ZNF211 | Body;Body | S_Shelf |
| **cg07369507** | 0.106506083 | 6.53E-08 | 6 | ZNF323;ZNF323;ZNF323;ZNF323;ZNF323;ZNF323 | TSS1500;TSS200;5'UTR;TSS200;5'UTR;TSS1500 |  |
| **cg16954629** | -0.088355984 | 3.13E-12 | 19 | ZNF888 | TSS200 |  |
| **cg06393529** | 0.210329073 | 0.023649747 | 6 |  |  |  |
| **cg10051493** | 0.166170871 | 0.029030707 | 16 |  |  |  |
| **cg23056271** | 0.161882425 | 0.02496661 | 5 |  |  |  |
| **cg16869625** | 0.139616525 | 0.034747266 | 1 |  |  |  |
| **cg10811640** | 0.139506275 | 0.04157351 | 3 |  |  |  |
| **cg04402345** | 0.131178308 | 0.008346029 | 5 |  |  |  |
| **cg01303155** | 0.096520016 | 0.021257875 | 7 |  |  |  |
| **cg26080286** | 0.090510439 | 1.03E-07 | 14 |  |  |  |
| **cg27611827** | 0.087635303 | 3.32E-10 | 17 |  |  | N_Shore |
| **cg12876682** | 0.081663336 | 0.03913196 | 1 |  |  |  |
| **cg15903595** | 0.081219551 | 0.047555178 | 20 |  |  |  |
| **cg08108965** | 0.080972296 | 7.51E-06 | 1 |  |  | S_Shore |
| **cg05764111** | 0.080705283 | 1.41E-06 | 17 |  |  |  |
| **cg11999665** | -0.080155987 | 1.64E-16 | 19 |  |  |  |
| **cg23860436** | -0.080527237 | 8.26E-11 | 12 |  |  |  |
| **cg13231117** | -0.080830089 | 2.99E-13 | 2 |  |  | S_Shore |
| **cg17424859** | -0.08090808 | 3.34E-13 | 6 |  |  |  |
| **cg02663382** | -0.080949685 | 0.00010429 | 19 |  |  | Island |
| **cg19751754** | -0.081166188 | 1.63E-08 | 17 |  |  |  |
| **cg09133791** | -0.081200351 | 3.30E-09 | 3 |  |  |  |
| **cg01249134** | -0.081206271 | 2.28E-11 | 18 |  |  | N_Shelf |
| **cg17537151** | -0.08164717 | 0.000171783 | 2 |  |  |  |
| **cg10709925** | -0.082054972 | 5.98E-13 | 3 |  |  | Island |
| **cg20767890** | -0.082353705 | 0.024617421 | 2 |  |  |  |
| **cg23347058** | -0.082473543 | 3.01E-14 | 2 |  |  |  |
| **cg26551069** | -0.082507144 | 6.35E-13 | 11 |  |  |  |
| **cg27478848** | -0.082826925 | 0.002508642 | 15 |  |  | S_Shore |
| **cg01789846** | -0.082835433 | 6.71E-14 | 4 |  |  | Island |
| **cg11805548** | -0.082884669 | 1.17E-11 | 2 |  |  |  |
| **cg11649715** | -0.08295658 | 0.001339646 | 1 |  |  | Island |
| **cg16058452** | -0.08307307 | 1.12E-10 | 15 |  |  |  |
| **cg23160376** | -0.083108526 | 5.74E-08 | 4 |  |  |  |
| **cg10087939** | -0.083221779 | 1.41E-13 | 3 |  |  |  |
| **cg11397033** | -0.083414359 | 5.65E-16 | 7 |  |  | Island |
| **cg05707985** | -0.083501627 | 0.035508754 | 7 |  |  |  |
| **cg22221266** | -0.083837674 | 4.30E-10 | 14 |  |  |  |
| **cg17678740** | -0.083861187 | 3.17E-13 | 7 |  |  | N_Shore |
| **cg09115837** | -0.084577179 | 1.64E-16 | 10 |  |  |  |
| **cg05742286** | -0.084713959 | 8.94E-18 | 12 |  |  | Island |
| **cg26537719** | -0.085188032 | 5.96E-06 | 11 |  |  |  |
| **cg06763665** | -0.085474839 | 1.42E-07 | 1 |  |  |  |
| **cg17461271** | -0.085678282 | 2.70E-13 | 1 |  |  |  |
| **cg00414890** | -0.085698692 | 0.02279283 | 19 |  |  | S_Shelf |
| **cg11753711** | -0.086008972 | 5.56E-06 | 7 |  |  |  |
| **cg04734610** | -0.086655255 | 2.00E-06 | 4 |  |  |  |
| **cg14380045** | -0.086705617 | 2.28E-12 | 9 |  |  | Island |
| **cg07594204** | -0.086952325 | 3.88E-11 | 7 |  |  |  |
| **cg16919064** | -0.087508981 | 0.042506978 | 2 |  |  |  |
| **cg15050398** | -0.087655129 | 3.95E-12 | 6 |  |  | N_Shelf |
| **cg06507271** | -0.087677391 | 6.07E-13 | 7 |  |  |  |
| **cg22615071** | -0.087679128 | 8.71E-12 | 3 |  |  |  |
| **cg16684608** | -0.087807376 | 0.000494044 | 1 |  |  | S_Shelf |
| **cg17095167** | -0.087897153 | 2.24E-13 | 13 |  |  | Island |
| **cg12050434** | -0.087931733 | 2.82E-13 | 12 |  |  |  |
| **cg15310722** | -0.088033552 | 4.31E-16 | 17 |  |  |  |
| **cg22805491** | -0.088223513 | 2.42E-14 | 14 |  |  |  |
| **cg09427936** | -0.088756169 | 9.54E-14 | 1 |  |  |  |
| **cg01686861** | -0.088758381 | 7.15E-11 | 16 |  |  |  |
| **cg12651293** | -0.088822922 | 1.18E-14 | 1 |  |  |  |
| **cg24382249** | -0.089075691 | 0.010700289 | 15 |  |  | N_Shore |
| **cg14645856** | -0.089204798 | 2.01E-16 | 6 |  |  |  |
| **cg00316520** | -0.089282864 | 4.07E-14 | 3 |  |  | N_Shelf |
| **cg26174329** | -0.089815931 | 2.27E-11 | 7 |  |  | N_Shelf |
| **cg06936108** | -0.089863892 | 4.37E-14 | 17 |  |  | S_Shore |
| **cg00305263** | -0.090099052 | 4.19E-15 | 1 |  |  |  |
| **cg19708055** | -0.090388476 | 0.006166782 | 6 |  |  |  |
| **cg08609238** | -0.090395986 | 7.29E-18 | 9 |  |  | Island |
| **cg01806526** | -0.090506045 | 1.30E-10 | 16 |  |  | N_Shore |
| **cg05526438** | -0.090557932 | 2.45E-15 | 5 |  |  |  |
| **cg23129217** | -0.090561476 | 1.17E-14 | 12 |  |  | S_Shore |
| **cg20916068** | -0.090630683 | 2.92E-15 | 15 |  |  |  |
| **cg13402292** | -0.090786018 | 1.05E-16 | 21 |  |  | Island |
| **cg21374546** | -0.091160145 | 2.70E-10 | 10 |  |  |  |
| **cg03008165** | -0.091670329 | 1.65E-18 | 11 |  |  |  |
| **cg03178542** | -0.092074276 | 2.57E-13 | 10 |  |  |  |
| **cg23877043** | -0.09214972 | 0.000575475 | 12 |  |  |  |
| **cg01593421** | -0.092359986 | 5.57E-12 | 7 |  |  |  |
| **cg20368377** | -0.092592244 | 3.35E-13 | 13 |  |  | N_Shore |
| **cg06919059** | -0.092680926 | 6.71E-10 | 1 |  |  |  |
| **cg12224834** | -0.092913944 | 2.28E-15 | 1 |  |  | N_Shelf |
| **cg07791468** | -0.092946849 | 5.42E-14 | 2 |  |  |  |
| **cg12223562** | -0.093177455 | 7.95E-08 | 2 |  |  |  |
| **cg06765686** | -0.093261331 | 9.15E-17 | 16 |  |  | N_Shelf |
| **cg25444615** | -0.093438068 | 5.23E-12 | 19 |  |  |  |
| **cg19166406** | -0.093570088 | 0.034486942 | 4 |  |  |  |
| **cg05466248** | -0.093711194 | 4.39E-16 | 14 |  |  |  |
| **cg06028934** | -0.093885603 | 3.16E-19 | 1 |  |  |  |
| **cg14661028** | -0.094874002 | 2.68E-14 | 1 |  |  | S_Shore |
| **cg14950698** | -0.095093024 | 5.19E-13 | 2 |  |  |  |
| **cg02738677** | -0.096102977 | 1.38E-15 | 3 |  |  |  |
| **cg04791162** | -0.09631518 | 8.97E-16 | 14 |  |  | Island |
| **cg19882132** | -0.096934395 | 3.63E-05 | 14 |  |  | S_Shelf |
| **cg09210005** | -0.097532735 | 2.71E-15 | 20 |  |  |  |
| **cg08365438** | -0.097550976 | 3.24E-12 | 10 |  |  |  |
| **cg07516076** | -0.097764277 | 3.36E-17 | 7 |  |  | Island |
| **cg14964336** | -0.098234828 | 0.006746326 | 4 |  |  | Island |
| **cg05521474** | -0.099262084 | 3.61E-17 | 5 |  |  |  |
| **cg09615411** | -0.100013335 | 0.04611852 | 8 |  |  |  |
| **cg21036766** | -0.100617414 | 7.48E-17 | 6 |  |  | Island |
| **cg14265604** | -0.100667129 | 3.30E-13 | 1 |  |  |  |
| **cg08876558** | -0.101771146 | 1.42E-15 | 11 |  |  |  |
| **cg04611493** | -0.101946205 | 2.64E-16 | 2 |  |  |  |
| **cg08450091** | -0.102307073 | 3.96E-14 | 3 |  |  | Island |
| **cg19064493** | -0.102594742 | 0.003245746 | 7 |  |  | Island |
| **cg03285130** | -0.103080077 | 1.43E-17 | 10 |  |  |  |
| **cg15123819** | -0.104268918 | 6.02E-09 | 5 |  |  |  |
| **cg07575344** | -0.104603652 | 2.23E-15 | 1 |  |  | N_Shelf |
| **cg06888084** | -0.104977304 | 2.02E-13 | 11 |  |  |  |
| **cg14357455** | -0.105534269 | 5.56E-13 | 6 |  |  | N_Shore |
| **cg04438374** | -0.10579378 | 1.43E-07 | 7 |  |  |  |
| **cg24568548** | -0.10706708 | 2.67E-12 | 11 |  |  | S_Shore |
| **cg15882280** | -0.107271474 | 9.91E-14 | 22 |  |  |  |
| **cg07867325** | -0.107498865 | 4.35E-16 | 3 |  |  | Island |
| **cg07538465** | -0.108203433 | 5.92E-16 | 12 |  |  |  |
| **cg14298830** | -0.108496343 | 0.035886837 | 20 |  |  |  |
| **cg23244452** | -0.108549556 | 1.71E-14 | 4 |  |  |  |
| **cg05909891** | -0.109127967 | 1.11E-17 | 12 |  |  |  |
| **cg02593636** | -0.110385675 | 2.25E-11 | 12 |  |  |  |
| **cg12615557** | -0.110658204 | 1.20E-15 | 1 |  |  |  |
| **cg07283849** | -0.115592623 | 0.00507435 | 7 |  |  |  |
| **cg16918438** | -0.115908484 | 0.012137837 | 14 |  |  |  |
| **cg14559409** | -0.116426998 | 2.65E-15 | 10 |  |  | Island |
| **cg14341057** | -0.117365889 | 1.64E-16 | 13 |  |  |  |
| **cg08653580** | -0.118218532 | 5.00E-18 | 11 |  |  |  |
| **cg16885723** | -0.121787936 | 0.036848959 | 14 |  |  |  |
| **cg07625092** | -0.12235083 | 9.95E-18 | 15 |  |  |  |
| **cg09048334** | -0.122747593 | 1.22E-10 | 6 |  |  | Island |
| **cg26769184** | -0.124716962 | 8.97E-18 | 22 |  |  |  |
| **cg13881513** | -0.126109931 | 1.52E-15 | 18 |  |  | N_Shelf |
| **cg03116837** | -0.127444669 | 0.000872384 | 8 |  |  | Island |
| **cg21653586** | -0.131045822 | 2.79E-13 | 11 |  |  |  |
| **cg11354479** | -0.134348729 | 0.000679683 | 9 |  |  | N_Shore |
| **cg06061636** | -0.139856276 | 2.60E-14 | 5 |  |  |  |
| **cg02217425** | -0.140181805 | 0.033738737 | 17 |  |  |  |
| **cg08323960** | -0.142471404 | 8.97E-18 | 10 |  |  |  |
| **cg06083932** | -0.158108232 | 0.035916109 | 19 |  |  | Island |
| **cg14317384** | -0.16063313 | 0.000582057 | 8 |  |  | Island |
| **cg23591463** | -0.175659935 | 0.024875825 | 12 |  |  | N_Shore |
| **cg03664889** | -0.1847802 | 0.03054324 | 1 |  |  |  |
| **cg17920646** | -0.230297459 | 0.000144321 | 8 |  |  | Island |

### Supplementary Table S5-a.

**The 207 differentially methylated probes shared between FES and PRS**

| Probe ID | Delta Beta | Adjust *P* value | chromosome | Gene Name | Gene Context | CpG Island |
| --- | --- | --- | --- | --- | --- | --- |
| cg00047657 | -0.097741286 | 3.16E-19 | 10 | STOX1;STOX1;STOX1;STOX1 | Body;Body;Body;Body |  |
| cg06028934 | -0.093885603 | 3.16E-19 | 1 |  |  |  |
| cg03388324 | -0.100890462 | 3.56E-19 | 6 | CEP85L;CEP85L;CEP85L | Body;Body;Body |  |
| cg18803079 | -0.133651124 | 3.56E-19 | 1 | EFCAB7;DLEU2L | Body;TSS200 |  |
| cg21610999 | -0.108998024 | 1.26E-18 | 17 | SPDYE4 | TSS200 |  |
| cg03008165 | -0.091670329 | 1.65E-18 | 11 |  |  |  |
| cg01291513 | -0.132823785 | 1.83E-18 | 3 | TSEN2;TSEN2;TSEN2;TSEN2;TSEN2;TSEN2;TSEN2;TSEN2;TSEN2 | ExonBnd;ExonBnd;ExonBnd;ExonBnd;Body;Body;Body;Body;Body |  |
| cg01603311 | -0.150608829 | 2.01E-18 | 17 | RNF157 | Body | N_Shore |
| cg18440889 | -0.100093141 | 2.01E-18 | 11 | SHANK2 | Body |  |
| cg08782481 | 0.089557052 | 4.45E-18 | 6 | HIST1H3I | 1stExon | Island |
| cg08653580 | -0.118218532 | 5.00E-18 | 11 |  |  |  |
| cg08609238 | -0.090395986 | 7.29E-18 | 9 |  |  | Island |
| cg05742286 | -0.084713959 | 8.94E-18 | 12 |  |  | Island |
| cg00389672 | -0.091348885 | 8.97E-18 | 17 | SSH2;SSH2;SSH2 | Body;Body;Body |  |
| cg05465443 | -0.103750409 | 8.97E-18 | 3 | RAD54L2 | Body |  |
| cg08323960 | -0.142471404 | 8.97E-18 | 10 |  |  |  |
| cg26769184 | -0.124716962 | 8.97E-18 | 22 |  |  |  |
| cg07625092 | -0.12235083 | 9.95E-18 | 15 |  |  |  |
| cg05909891 | -0.109127967 | 1.11E-17 | 12 |  |  |  |
| cg03285130 | -0.103080077 | 1.43E-17 | 10 |  |  |  |
| cg07516076 | -0.097764277 | 3.36E-17 | 7 |  |  | Island |
| cg05521474 | -0.099262084 | 3.61E-17 | 5 |  |  |  |
| cg12187850 | -0.133354798 | 3.86E-17 | 5 | WDR70 | Body |  |
| cg07074586 | -0.102825101 | 5.69E-17 | 12 | MYL6;MYL6 | 3'UTR;3'UTR |  |
| cg00832928 | -0.109318273 | 6.35E-17 | 3 | SELT | Body | Island |
| cg02173085 | -0.140644631 | 6.35E-17 | 2 | METTL5;METTL5;METTL5 | Body;Body;Body | N_Shelf |
| cg13593809 | -0.115922448 | 8.58E-17 | 17 | LOC101559451;PELP1;PELP1 | Body;TSS1500;TSS1500 | S_Shore |
| cg01490296 | -0.125246726 | 9.15E-17 | 10 | MCM10;MCM10 | 5'UTR;5'UTR | S_Shore |
| cg06765686 | -0.093261331 | 9.15E-17 | 16 |  |  | N_Shelf |
| cg04711526 | -0.12043251 | 9.19E-17 | 14 | NDUFB1 | 3'UTR |  |
| cg10299917 | -0.095256466 | 1.04E-16 | 22 | LRP5L | 5'UTR |  |
| cg12113395 | -0.11329209 | 1.24E-16 | 3 | RNF7;RNF7;RNF7;RNF7;RNF7 | 3'UTR;3'UTR;3'UTR;Body;Body |  |
| cg22552448 | -0.093851016 | 1.31E-16 | 12 | PCBP2;PCBP2;PCBP2;PCBP2;PCBP2;PCBP2;PCBP2;PCBP2;PCBP2;PCBP2;PCBP2;PCBP2;PCBP2;PCBP2 | ExonBnd;ExonBnd;ExonBnd;ExonBnd;ExonBnd;ExonBnd;ExonBnd;Body;Body;Body;Body;Body;Body;Body | S_Shelf |
| cg03182374 | -0.089962482 | 1.45E-16 | 6 | PTP4A1 | Body |  |
| cg22431028 | -0.12549776 | 1.45E-16 | 14 | MNAT1;MNAT1 | Body;Body |  |
| cg06363265 | -0.095685807 | 1.49E-16 | 7 | SEPT7;SEPT7;SEPT7 | Body;Body;Body |  |
| cg02449698 | -0.097466296 | 1.63E-16 | 1 | EFCAB7;DLEU2L | Body;TSS200 |  |
| cg06764475 | -0.10389269 | 1.64E-16 | 6 | PRIM2 | Body |  |
| cg14341057 | -0.117365889 | 1.64E-16 | 13 |  |  |  |
| cg22726533 | -0.1058498 | 1.64E-16 | 7 | YAE1D1;YAE1D1 | Body;Body | S_Shelf |
| cg07341559 | -0.112565549 | 1.80E-16 | 11 | DEFB108B | Body |  |
| cg10297223 | -0.140344732 | 1.80E-16 | 3 | AGTR1;AGTR1;AGTR1;AGTR1 | TSS1500;TSS1500;TSS1500;TSS1500 | N_Shore |
| cg11893547 | -0.100040526 | 1.98E-16 | 14 | PSMA3;PSMA3;PSMA3 | Body;Body;Body |  |
| cg26840922 | -0.091195405 | 1.98E-16 | 3 | ATP1B3 | Body |  |
| cg03900366 | -0.116168183 | 2.23E-16 | 1 | MACF1 | Body |  |
| cg17983571 | -0.129713309 | 2.41E-16 | 10 | JMJD1C | Body |  |
| cg12601237 | -0.104305165 | 2.52E-16 | 10 | ST8SIA6-AS1;ST8SIA6 | TSS1500;Body |  |
| cg17896229 | -0.083783512 | 2.62E-16 | 20 | PROKR2 | 1stExon | N_Shore |
| cg04611493 | -0.101946205 | 2.64E-16 | 2 |  |  |  |
| cg06774087 | -0.088777831 | 2.68E-16 | 5 | LOC441089 | Body |  |
| cg06649000 | -0.101677699 | 3.93E-16 | 12 | PCBP2;PCBP2;PCBP2;PCBP2;PCBP2;PCBP2;PCBP2 | Body;Body;Body;Body;Body;Body;Body |  |
| cg13442241 | -0.096080271 | 4.29E-16 | 8 | EMC2 | Body |  |
| cg15310722 | -0.088033552 | 4.31E-16 | 17 |  |  |  |
| cg05466248 | -0.093711194 | 4.39E-16 | 14 |  |  |  |
| cg18642567 | -0.09054009 | 4.39E-16 | 14 | RPGRIP1 | TSS1500 |  |
| cg12361190 | -0.091781808 | 5.85E-16 | 6 | LOC441155;EYS;EYS | TSS200;Body;Body | Island |
| cg07538465 | -0.108203433 | 5.92E-16 | 12 |  |  |  |
| cg19933320 | -0.103398658 | 6.09E-16 | 7 | ZNF107;ZNF107 | TSS1500;TSS1500 | N_Shore |
| cg13961165 | -0.107331737 | 7.03E-16 | 2 | C2orf27A;C2orf27A | 1stExon;5'UTR |  |
| cg26929161 | -0.091910698 | 7.24E-16 | 20 | ZFP64 | Body | Island |
| cg04791162 | -0.09631518 | 8.97E-16 | 14 |  |  | Island |
| cg26296769 | -0.109110993 | 9.80E-16 | 6 | KLC4;KLC4;KLC4;MRPL2;MRPL2 | TSS1500;TSS1500;TSS1500;Body;Body | N_Shore |
| cg12953621 | -0.088639899 | 1.07E-15 | 12 | ATP5G2;ATP5G2 | Body;Body |  |
| cg25477163 | -0.102234351 | 1.12E-15 | 19 | ZNF211;ZNF211 | Body;Body | S_Shelf |
| cg25700305 | 0.089232304 | 1.16E-15 | 1 | PRPF3 | TSS200 | Island |
| cg12615557 | -0.110658204 | 1.20E-15 | 1 |  |  |  |
| cg04204557 | -0.105946272 | 1.29E-15 | 21 | KRTAP10-11;C21orf29 | TSS1500;Body |  |
| cg02738677 | -0.096102977 | 1.38E-15 | 3 |  |  |  |
| cg08876558 | -0.101771146 | 1.42E-15 | 11 |  |  |  |
| cg13881513 | -0.126109931 | 1.52E-15 | 18 |  |  | N_Shelf |
| cg24192846 | -0.087233239 | 1.56E-15 | 6 | PPP1R11 | 3'UTR | N_Shore |
| cg07575344 | -0.104603652 | 2.23E-15 | 1 |  |  | N_Shelf |
| cg14115740 | -0.115739466 | 2.25E-15 | 9 | FANCC | 5'UTR | Island |
| cg19963700 | -0.083281438 | 2.42E-15 | 1 | ROR1;ROR1 | Body;Body |  |
| cg18370902 | -0.093058856 | 2.60E-15 | 8 | FBXO25;FBXO25;FBXO25 | 5'UTR;Body;Body |  |
| cg27133230 | -0.112314229 | 2.66E-15 | 20 | RBPJL | Body | S_Shore |
| cg09210005 | -0.097532735 | 2.71E-15 | 20 |  |  |  |
| cg02941728 | -0.097724423 | 3.61E-15 | 11 | CCDC86 | 3'UTR | N_Shelf |
| cg14150708 | -0.091939955 | 3.62E-15 | 3 | RAD18 | Body |  |
| cg00084591 | -0.101034146 | 3.95E-15 | 9 | ERP44 | Body |  |
| cg15810754 | -0.109736884 | 4.00E-15 | 15 | TJP1;TJP1;TJP1;TJP1 | Body;Body;Body;Body |  |
| cg01011918 | -0.094503827 | 4.17E-15 | 1 | SPRR2D;SPRR2D | 1stExon;5'UTR |  |
| cg00305263 | -0.090099052 | 4.19E-15 | 1 |  |  |  |
| cg06276780 | -0.080154088 | 8.51E-15 | 8 | UBE2V2 | Body |  |
| cg04667820 | -0.083081956 | 8.98E-15 | 2 | VPS54;VPS54 | 5'UTR;5'UTR |  |
| cg03889742 | -0.123602282 | 9.45E-15 | 15 | MYO5A;MYO5A | Body;Body |  |
| cg23129217 | -0.090561476 | 1.17E-14 | 12 |  |  | S_Shore |
| cg00872998 | -0.097237429 | 1.21E-14 | 9 | GDA;GDA;GDA;GDA | Body;Body;Body;Body |  |
| cg03017824 | -0.10296244 | 1.45E-14 | 1 | LOC100130093;LOC100130093 | TSS1500;TSS1500 | N_Shore |
| cg23244452 | -0.108549556 | 1.71E-14 | 4 |  |  |  |
| cg00574307 | -0.098953029 | 1.71E-14 | 13 | DCT;DCT | Body;Body |  |
| cg19476594 | -0.100055457 | 1.77E-14 | 1 | ABL2;ABL2;ABL2;ABL2;ABL2;ABL2;ABL2;ABL2 | Body;Body;Body;Body;Body;Body;Body;Body |  |
| cg10369197 | -0.104480609 | 2.29E-14 | 17 | SLC39A11;SLC39A11 | Body;Body |  |
| cg22805491 | -0.088223513 | 2.42E-14 | 14 |  |  |  |
| cg06061636 | -0.139856276 | 2.60E-14 | 5 |  |  |  |
| cg14661028 | -0.094874002 | 2.68E-14 | 1 |  |  | S_Shore |
| cg24013599 | -0.096272778 | 2.69E-14 | 8 | RNF19A;RNF19A;RNF19A | 5'UTR;5'UTR;1stExon |  |
| cg04158140 | -0.090415497 | 3.02E-14 | 7 | CALN1;CALN1 | Body;Body |  |
| cg17285822 | -0.089787214 | 3.61E-14 | 9 | TMEM245 | TSS1500 | S_Shore |
| cg08450091 | -0.102307073 | 3.96E-14 | 3 |  |  | Island |
| cg00316520 | -0.089282864 | 4.07E-14 | 3 |  |  | N_Shelf |
| cg06936108 | -0.089863892 | 4.37E-14 | 17 |  |  | S_Shore |
| cg02718464 | -0.10596985 | 4.99E-14 | 13 | SUGT1L1;SLC25A15;MIR621 | Body;3'UTR;TSS1500 |  |
| cg07791468 | -0.092946849 | 5.42E-14 | 2 |  |  |  |
| cg05790002 | -0.100272481 | 5.65E-14 | 16 | SDR42E1 | Body |  |
| cg03472399 | -0.096671242 | 6.04E-14 | 13 | LOC101928697 | Body |  |
| cg01789846 | -0.082835433 | 6.71E-14 | 4 |  |  | Island |
| cg11309891 | -0.091937625 | 7.34E-14 | 11 | RAB6A;RAB6A | 3'UTR;3'UTR |  |
| cg11792281 | -0.129788841 | 8.31E-14 | 17 | NLK | Body |  |
| cg09427936 | -0.088756169 | 9.54E-14 | 1 |  |  |  |
| cg15882280 | -0.107271474 | 9.91E-14 | 22 |  |  |  |
| cg00218840 | -0.097389953 | 1.00E-13 | 1 | DUSP5P1;RHOU | Body;Body | S_Shore |
| cg01993027 | -0.095296676 | 1.21E-13 | 1 | SRSF4 | Body |  |
| cg19729503 | -0.095308975 | 1.32E-13 | 20 | ZHX3 | 5'UTR |  |
| cg10087939 | -0.083221779 | 1.41E-13 | 3 |  |  |  |
| cg00622907 | -0.082078812 | 1.52E-13 | 8 | TSNARE1 | Body | N_Shelf |
| cg06205244 | -0.095389615 | 1.69E-13 | 6 | SNX14;SNX14;SNX14;SNX14 | 5'UTR;Body;Body;Body | N_Shelf |
| cg07788360 | -0.088373174 | 1.70E-13 | 13 | MRPS31 | Body |  |
| cg06888084 | -0.104977304 | 2.02E-13 | 11 |  |  |  |
| cg10967735 | -0.084681844 | 2.26E-13 | 10 | CPEB3 | Body |  |
| cg04676559 | -0.081954282 | 2.34E-13 | 17 | LOC100131347;PLXDC1 | Body;Body |  |
| cg03178542 | -0.092074276 | 2.57E-13 | 10 |  |  |  |
| cg21653586 | -0.131045822 | 2.79E-13 | 11 |  |  |  |
| cg15004555 | -0.089550348 | 2.85E-13 | 1 | AIM2 | Body |  |
| cg17678740 | -0.083861187 | 3.17E-13 | 7 |  |  | N_Shore |
| cg10006801 | -0.102435989 | 3.24E-13 | 17 | MAP2K3;MAP2K3 | Body;Body |  |
| cg26118675 | -0.100122913 | 3.28E-13 | 2 | TRIM43 | TSS1500 |  |
| cg14265604 | -0.100667129 | 3.30E-13 | 1 |  |  |  |
| cg17424859 | -0.08090808 | 3.34E-13 | 6 |  |  |  |
| cg20368377 | -0.092592244 | 3.35E-13 | 13 |  |  | N_Shore |
| cg11320438 | -0.082387063 | 3.58E-13 | 19 | SIGLEC11;SIGLEC11 | Body;Body |  |
| cg06051442 | -0.092518373 | 3.61E-13 | 9 | FAM120A;FAM120A;FAM120A;FAM120A | Body;Body;Body;Body | S_Shore |
| cg00470768 | -0.103877017 | 4.34E-13 | 15 | INO80 | Body |  |
| cg14783439 | 0.089644199 | 5.14E-13 | 19 | SH3GL1;SH3GL1;SH3GL1 | 1stExon;1stExon;1stExon | Island |
| cg06507271 | -0.087677391 | 6.07E-13 | 7 |  |  |  |
| cg21581312 | -0.121402959 | 6.34E-13 | 15 | LOC723972 | TSS200 |  |
| cg26551069 | -0.082507144 | 6.35E-13 | 11 |  |  |  |
| cg19645279 | -0.085698023 | 7.35E-13 | 5 | NR3C1;NR3C1;NR3C1;NR3C1;NR3C1;NR3C1;NR3C1;NR3C1;NR3C1;NR3C1;NR3C1;NR3C1;NR3C1;NR3C1;NR3C1 | Body;Body;Body;Body;Body;Body;Body;Body;Body;Body;Body;Body;Body;Body;Body |  |
| cg11963436 | -0.130673697 | 7.89E-13 | 10 | TNKS2 | Body |  |
| cg16447252 | -0.084110907 | 1.59E-12 | 10 | SVIL-AS1;SVIL;SVIL | Body;Body;Body |  |
| cg14102005 | -0.088188813 | 1.85E-12 | 14 | PELI2 | Body |  |
| cg10475689 | -0.082712357 | 1.94E-12 | 16 | RUNDC2A | Body |  |
| cg14380045 | -0.086705617 | 2.28E-12 | 9 |  |  | Island |
| cg24568548 | -0.10706708 | 2.67E-12 | 11 |  |  | S_Shore |
| cg24203469 | -0.08411424 | 2.84E-12 | 7 | LINC01162 | TSS200 |  |
| cg23024233 | -0.114052979 | 3.68E-12 | 7 | TRIM4;TRIM4 | Body;Body |  |
| cg15050398 | -0.087655129 | 3.95E-12 | 6 |  |  | N_Shelf |
| cg06936779 | 0.131878233 | 4.67E-12 | 1 | PIP5K1A;PIP5K1A;PIP5K1A;PIP5K1A;PIP5K1A;PIP5K1A;PIP5K1A;PIP5K1A | 5'UTR;1stExon;5'UTR;1stExon;5'UTR;1stExon;1stExon;5'UTR | Island |
| cg03278573 | -0.098801829 | 5.03E-12 | 5 | DAP;DAP | Body;Body |  |
| cg02990553 | -0.09804199 | 5.03E-12 | 17 | KRT28 | 1stExon | S_Shelf |
| cg25444615 | -0.093438068 | 5.23E-12 | 19 |  |  |  |
| cg12045156 | -0.085408503 | 7.55E-12 | 17 | MAP2K6 | Body |  |
| cg06871323 | -0.095674554 | 1.24E-11 | 7 | DPY19L1 | Body |  |
| cg02593636 | -0.110385675 | 2.25E-11 | 12 |  |  |  |
| cg26174329 | -0.089815931 | 2.27E-11 | 7 |  |  | N_Shelf |
| cg24674363 | -0.114029617 | 2.44E-11 | 3 | LOC101927056;LOC101927056 | Body;Body |  |
| cg11962649 | -0.095099665 | 3.06E-11 | 9 | HSPA5 | Body | N_Shore |
| cg16058452 | -0.08307307 | 1.12E-10 | 15 |  |  |  |
| cg10182306 | -0.086947072 | 1.13E-10 | 1 | CLCNKA;CLCNKA;CLCNKA | Body;Body;Body |  |
| cg09048334 | -0.122747593 | 1.22E-10 | 6 |  |  | Island |
| cg14079545 | -0.102641605 | 1.41E-10 | 6 | SYNGAP1 | Body | N_Shelf |
| cg09141640 | -0.084054794 | 1.79E-10 | 22 | LOC100506679 | TSS1500 |  |
| cg08152083 | -0.08449889 | 1.96E-10 | 17 | CDRT15L2 | TSS1500 |  |
| cg02141602 | -0.093702052 | 2.11E-10 | 19 | MEIS3;MEIS3;MEIS3 | TSS200;TSS200;TSS200 | S_Shore |
| cg21374546 | -0.091160145 | 2.70E-10 | 10 |  |  |  |
| cg20443413 | -0.081388445 | 2.76E-10 | 6 | SGK1 | Body |  |
| cg08713543 | -0.081103515 | 2.99E-10 | 16 | TMEM231;TMEM231;TMEM231 | Body;Body;Body |  |
| cg22221266 | -0.083837674 | 4.30E-10 | 14 |  |  |  |
| cg07240877 | -0.080194793 | 4.56E-10 | 1 | LOC339535 | TSS200 |  |
| cg06919059 | -0.092680926 | 6.71E-10 | 1 |  |  |  |
| cg22682254 | -0.082920415 | 2.04E-09 | 17 | CRYBA1 | TSS1500 |  |
| cg09133791 | -0.081200351 | 3.30E-09 | 3 |  |  |  |
| cg17988535 | 0.10108455 | 3.42E-09 | 13 | RASA3 | Body | S_Shore |
| cg24299813 | -0.081711445 | 3.71E-09 | 11 | TMEM109 | 5'UTR | S_Shelf |
| cg25960044 | -0.100519809 | 5.12E-09 | 4 | MANBA | Body |  |
| cg15123819 | -0.104268918 | 6.02E-09 | 5 |  |  |  |
| cg07744399 | -0.08662144 | 1.03E-08 | 7 | XRCC2 | Body |  |
| cg19751754 | -0.081166188 | 1.63E-08 | 17 |  |  |  |
| cg07068730 | -0.080461562 | 2.45E-08 | 3 | DLG1;DLG1;DLG1;DLG1 | Body;Body;Body;Body |  |
| cg07369507 | 0.106506083 | 6.53E-08 | 6 | ZNF323;ZNF323;ZNF323;ZNF323;ZNF323;ZNF323 | TSS1500;TSS200;5'UTR;TSS200;5'UTR;TSS1500 |  |
| cg12223562 | -0.093177455 | 7.95E-08 | 2 |  |  |  |
| cg16074739 | -0.084669196 | 8.00E-08 | 8 | MTUS1;MTUS1;MTUS1;MTUS1 | TSS1500;TSS1500;Body;Body |  |
| cg14922317 | 0.09337593 | 1.12E-07 | 16 | ATXN2L;ATXN2L;ATXN2L;ATXN2L;ATXN2L;ATXN2L;ATXN2L;ATXN2L;ATXN2L;ATXN2L;ATXN2L;ATXN2L;ATXN2L;ATXN2L | 5'UTR;5'UTR;5'UTR;5'UTR;5'UTR;5'UTR;5'UTR;1stExon;1stExon;1stExon;1stExon;1stExon;1stExon;1stExon | Island |
| cg06763665 | -0.085474839 | 1.42E-07 | 1 |  |  |  |
| cg04438374 | -0.10579378 | 1.43E-07 | 7 |  |  |  |
| cg00500007 | -0.094713817 | 2.55E-07 | 13 | NBEA | Body |  |
| cg23355197 | -0.083522488 | 6.31E-07 | 8 | LOC100506990;LOC100506990 | Body;Body |  |
| cg04628742 | -0.108989623 | 1.13E-06 | 6 | HLA-J;NCRNA00171 | TSS1500;Body | N_Shore |
| cg23961843 | -0.085306797 | 1.44E-06 | 1 | RGL1;APOBEC4 | 5'UTR;TSS1500 |  |
| ch.8.2353618R | 0.093454342 | 2.11E-06 | 8 | SAMD12 | Body |  |
| cg03289961 | -0.090120622 | 4.94E-06 | 2 | HADHB;HADHB;HADHB;HADHB;HADHB | ExonBnd;ExonBnd;Body;Body;Body |  |
| cg11753711 | -0.086008972 | 5.56E-06 | 7 |  |  |  |
| cg08797194 | -0.092037972 | 1.64E-05 | 13 | UGGT2 | Body | N_Shore |
| cg02256315 | -0.094863894 | 1.73E-05 | 11 | LDHAL6A;LDHAL6A | 3'UTR;3'UTR |  |
| cg00586123 | -0.080548481 | 6.02E-05 | 6 | LY86-AS1 | Body |  |
| cg01973725 | -0.103979676 | 8.13E-05 | 5 | RNU5E;RNU5D;ZCCHC9;ZCCHC9;ZCCHC9 | Body;Body;3'UTR;3'UTR;3'UTR |  |
| cg08477687 | -0.124270197 | 9.22E-05 | 1 | MIR1977 | TSS1500 |  |
| cg02663382 | -0.080949685 | 0.00010429 | 19 |  |  | Island |
| cg10514580 | -0.090880774 | 0.000694569 | 17 | SLFN12;SLFN12 | TSS1500;TSS200 |  |
| cg24655310 | -0.103296524 | 0.00080019 | 19 | CYP4F11;CYP4F11 | 1stExon;Body |  |
| cg27478848 | -0.082826925 | 0.002508642 | 15 |  |  | S_Shore |
| cg22273354 | -0.091845374 | 0.002737024 | 11 | NUP160 | Body |  |
| cg07283849 | -0.115592623 | 0.00507435 | 7 |  |  |  |
| cg04402345 | 0.131178308 | 0.008346029 | 5 |  |  |  |
| cg00114966 | -0.112900114 | 0.009222021 | 1 | LHX9;LHX9 | Body;Body | S_Shelf |
| cg14298830 | -0.108496343 | 0.035886837 | 20 |  |  |  |
| cg16919064 | -0.087508981 | 0.042506978 | 2 |  |  |  |

### Supplementary Table S5-b.

**The 148 unique genes shared between FES and PRS**

Gene_Name

**ABL2, AGTR1, AIM2, APOBEC4, ATP1B3, ATP5G2, ATXN2L, C21orf29, C2orf27A, CALN1, CCDC86, CDRT15L2, CEP85L, CLCNKA, CPEB3, CRYBA1, CYP4F11, DAP, DCT, DEFB108B, DLEU2L, DLG1, DPY19L1, DUSP5P1, EFCAB7, EMC2, ERP44, EYS, FAM120A, FANCC, FBXO25, GDA, HADHB, HIST1H3I, HLA-J, HSPA5, INO80, JMJD1C, KLC4, KRT28, KRTAP10-11, LDHAL6A, LHX9, LINC01162, LOC100130093, LOC100131347, LOC100506679, LOC100506990, LOC101559451, LOC101927056, LOC101928697, LOC339535, LOC441089, LOC441155, LOC723972, LRP5L, LY86-AS1, MACF1, MANBA, MAP2K3, MAP2K6, MCM10, MEIS3, METTL5, MIR1977, MIR621, MNAT1, MRPL2, MRPS31, MTUS1, MYL6, MYO5A, NBEA, NCRNA00171, NDUFB1, NLK, NR3C1, NUP160, PCBP2, PELI2, PELP1, PIP5K1A, PLXDC1, PPP1R11, PRIM2, PROKR2, PRPF3, PSMA3, PTP4A1, RAB6A, RAD18, RAD54L2, RASA3, RBPJL, RGL1, RHOU, RNF157, RNF19A, RNF7, RNU5D, RNU5E, ROR1, RPGRIP1, RUNDC2A, SAMD12, SDR42E1, SELT, SEPT7, SGK1, SH3GL1, SHANK2, SIGLEC11, SLC25A15, SLC39A11, SLFN12, SNX14, SPDYE4, SPRR2D, SRSF4, SSH2, ST8SIA6, ST8SIA6-AS1, STOX1, SUGT1L1, SVIL, SVIL-AS1, SYNGAP1, TJP1, TMEM109, TMEM231, TMEM245, TNKS2, TRIM4, TRIM43, TSEN2, TSNARE1, UBE2V2, UGGT2, VPS54, WDR70, XRCC2, YAE1D1, ZCCHC9, ZFP64, ZHX3, ZNF107, ZNF211, ZNF323**

### Supplementary Table S5-c.

**The 128 reported genes.**

**Gene_Name**

ABL2, AGTR1, AIM2, APOBEC4, ATP1B3, ATP5G2, ATXN2L, C21orf29, C2orf27A, CALN1, CCDC86, CEP85L, CLCNKA, CPEB3, CRYBA1, CYP4F11, DAP, DCT, DEFB108B, DLEU2L, DLG1, DPY19L1, DUSP5P1, EFCAB7, EMC2, ERP44, EYS, FAM120A, FANCC, FBXO25, GDA, HADHB, HIST1H3I, HLA-J, HSPA5, INO80, JMJD1C, KLC4, KRT28, LDHAL6A, LHX9, LOC100506990, LRP5L, LY86-AS1, MACF1, MANBA, MAP2K3, MAP2K6, MCM10, MEIS3, METTL5, MIR621, MNAT1, MRPL2, MRPS31, MTUS1, MYL6, MYO5A, NBEA, NDUFB1, NLK, NR3C1, NUP160, PCBP2, PELI2, PELP1, PIP5K1A, PLXDC1, PPP1R11, PRIM2, PROKR2, PRPF3, PSMA3, PTP4A1, RAB6A, RAD18, RAD54L2, RASA3, RBPJL, RGL1, RHOU, RNF157, RNF19A, RNF7, ROR1, RPGRIP1, RUNDC2A, SAMD12, SELT, SEPT7, SGK1, SH3GL1, SHANK2, SIGLEC11, SLC25A15, SLC39A11, SLFN12, SNX14, SPRR2D, SRSF4, SSH2, ST8SIA6, ST8SIA6-AS1, STOX1, SVIL, SVIL-AS1, SYNGAP1, TJP1, TMEM109, TMEM231, TMEM245, TNKS2, TRIM4, TRIM43, TSEN2, TSNARE1, UBE2V2, UGGT2, VPS54, WDR70, XRCC2, YAE1D1, ZCCHC9, ZFP64, ZHX3, ZNF107, ZNF211, ZNF323

### Supplementary Table S6-a.

**The top 15 significant signatures related to patients with schizophrenia in the GO category of differentially methylated probes.**

| GO ID | GO term | GO category | Symbols in list | Fold enrichment | *P* value | FDR |
| --- | --- | --- | --- | --- | --- | --- |
| GO:0005737 | cytoplasm | cellular_component | RHOU,GDA,TSNARE1,ATP5G2,MNAT1,FASN,ST8SIA6,MYL6,PDE4DIP,NBPF20,NBPF9,PSMD2,FEN1,DISC1,SHANK2,TRIM4,PCBP2,MAP2K6,PLXDC1,SPRR2D,RGS7,SYNGAP1,SEPT7,RNF7,VPS54,UBE2V2,PPP1R11,EMC2,KLC4,MRPL2,SLC35A4,APBB3,SSH2,MCM10,FANCC,NAPG,MACF1,STOX1,NR3C1,ZHX3,METAP1,CALN1,AGBL4,SVIL,NDUFB1,CEP85L,HSPA5,RASA3,MAP2K3,ERP44,RGL1,SGK1,ATP1B3,PELI2,ABL2,ROR1,PTP4A1,SUMF2,CCT6A,PDCD6,METTL5,SPTA1,CSDE1,NRAS,TSEN2,KRT28,PSMA3,RNF19A,MYO5A,ANAPC10,MTUS1,CRYBA1,SLC25A15,PELP1,FAM120A,RAB6A,MRPS31,NBEA,IPO11,SNX2,MANBA,DLG1,EHD3,AIM2,NLK,RPSAP58,TMEM109,GMPR,DEFA4,WFS1,RPL26L1,GOLGA8B,NBPF10,PRPF3,GALNT5,HNRNPF,APBB2,DCT,HADHB,SKAP2,SH3GL1,TJP1,TNKS2,NBPF8,LDHAL6A,CYP4F12,RNF14,UGGT2,ARL17A,LATS2,CYP4F11,RGMA,GTF2IRD1,PIP5K1A,NLRP12,EIF2AK2,RNASET2,DAZAP1,EEA1,PTPRN2,PANX2,AK8,SNX27,CARD11,NF1,NUP160,PABPC1,ATXN2L,STRADA,SLC39A2,ATF6,SYNC,CRYGD,WNK1,ELK3,ASAP1,OSBPL3,DNAH17,EYA3,TRIM14,SPSB4,MAP3K8,SYT1,RALBP1,SPG7,FZD1,BPTF,TRIM8,MICA,HLA-DPA1,TECR,MYBPC1,SLC27A1,BAIAP2L1,OXR1,LMTK2,AGBL1,DPYD,CYLD,OSTF1,SERPINB2,ITCH,APOL6,JPH3,CRADD,EGLN1,MRPL28,SORBS1,BRSK2,FAT1,FRMD4A,SERPINE2,MYO16,UGT2B17,DOCK2,ELMO1,ABR,MIER1,UBASH3B,ERBB4,CTSH,UFSP2,PACS1,PARVB,ADAP1,ST8SIA5,SLC8A1,DAAM1,TESC,HTRA1,SGMS1,RYR3,SLK,TBCD,LGALS8,RPTOR,SLC25A21,AKR1B1,DDX60,FNDC1,UGT2B15,ATG4D,RPS6KA2,PASK,DUSP3,USP15,IL26,SH3BP4,HOOK2,MGAT1,ALOX15B,ZCCHC7,GRB10,MYO1D,TDH,PXDNL,ZMIZ1,VWF,CNST,EEF1E1,FARP1,AHRR,HDAC4,CUL2,IL22RA2,SCD5,PPP2R1A,AK3,LHPP,PAK4,PEX14,RPS6KB1,PGM2L1,ST3GAL6,FNDC3B,TUBA3E,PRDM2,PITRM1,CUL3,PBX1,PARVA,PAWR,PODXL,WNK2,PLA2G4A,ITGA1,COL5A1,SPATS2L,P2RX6,TEK,GSTM1,HAL,OTOF,CAMK1D,NSF | 1.223796 | 5.57E-07 | 0.002097 |
| GO:0005622 | intracellular | cellular_component | RHOU,GDA,ZFP64,TSNARE1,ATP5G2,MNAT1,ZNF808,FASN,ST8SIA6,MYL6,PDE4DIP,NBPF20,NBPF9,PSMD2,FEN1,DISC1,SHANK2,ZNF107,KRTAP10-11,TRIM4,PCBP2,MAP2K6,CCDC86,PLXDC1,PRIM2,SPRR2D,BRD1,RGS7,RAD18,SYNGAP1,SEPT7,RNF7,VPS54,ZNF211,UBE2V2,RBPJL,PPP1R11,EMC2,KLC4,MRPL2,SLC35A4,APBB3,SSH2,MPHOSPH10,MCM10,FANCC,NAPG,DUSP5,MACF1,E2F3,INO80,TMEM231,STOX1,NR3C1,ZHX3,METAP1,CALN1,AGBL4,SVIL,NDUFB1,CEP85L,MEIS3,HSPA5,RASA3,MAP2K3,ERP44,RGL1,SGK1,FBXO25,ATP1B3,PELI2,ABL2,ROR1,PTP4A1,RAD54L2,SUMF2,CCT6A,PDCD6,METTL5,RPGRIP1,SPTA1,CSDE1,NRAS,TSEN2,KRT28,PSMA3,RNF19A,TRIM43,MYO5A,JMJD1C,ANAPC10,SRSF4,MTUS1,CRYBA1,SLC25A15,PELP1,NUDT5,FAM120A,RAB6A,MRPS31,XRCC2,NBEA,IPO11,SNX2,MANBA,DLG1,EHD3,AIM2,NLK,RPSAP58,TMEM109,GMPR,DEFA4,WFS1,RPL26L1,GOLGA8B,DDX59,ZNF138,ZNF264,NBPF10,PRPF3,GALNT5,HNRNPF,APBB2,DCT,HADHB,SKAP2,SH3GL1,TJP1,TNKS2,NBPF8,LDHAL6A,CYP4F12,RNF14,UGGT2,ARL17A,ECT2L,LATS2,CYP4F11,RGMA,GTF2IRD1,PIP5K1A,NLRP12,EIF2AK2,RNASET2,DAZAP1,EEA1,L3MBTL4,PTPRN2,PANX2,AK8,SNX27,CARD11,NF1,ESRRG,NUP160,PABPC1,ATXN2L,STRADA,SLC39A2,ATF6,SYNC,SMARCAD1,CRYGD,WNK1,ELK3,ASAP1,OSBPL3,DNAH17,EYA3,TRIM14,SPSB4,MAP3K8,SYT1,RALBP1,LHX9,ZNF714,SPG7,FZD1,BPTF,TRIM8,MICA,ZNF492,PARP12,HLA-DPA1,TECR,MYBPC1,SLC27A1,RAD51B,BAIAP2L1,OXR1,ZSCAN22,TBC1D1,LMTK2,TCF7L1,ISX,PKD1L1,AGBL1,DPYD,PSD3,HELZ,CYLD,OSTF1,GTF3C4,DDX31,TBC1D22A,SERPINB2,ITCH,APOL6,PLAGL1,JPH3,CRADD,EGLN1,SMC5,MYT1L,MRPL28,CDC40,SORBS1,KRTAP3-1,BRSK2,BHLHE41,FAT1,PPP2R2C,LOX,FRMD4A,SERPINE2,MYO16,UGT2B17,DOCK2,TSHZ2,ELMO1,ABR,MIER1,UBASH3B,ERBB4,CTSH,UFSP2,PACS1,PARVB,ADAP1,ST8SIA5,SLC8A1,DAAM1,TESC,HTRA1,SGMS1,RYR3,SLK,TBCD,LGALS8,RPTOR,SLC25A21,AKR1B1,DDX60,FNDC1,UGT2B15,ATG4D,GNG7,RPS6KA2,PASK,MED13L,DUSP3,USP15,IL26,SH3BP4,HOOK2,MGAT1,ALOX15B,ZCCHC7,GRB10,MYO1D,FAM120B,TDH,PXDNL,ADRA1B,ZMIZ1,VWF,CNST,EEF1E1,YPEL1,FARP1,AHRR,HDAC4,BANP,CUL2,IL22RA2,SCD5,PPP2R1A,AK3,LHPP,PAK4,PEX14,RPS6KB1,PGM2L1,ST3GAL6,GNAL,FNDC3B,TUBA3E,PRDM2,PITRM1,TSHZ1,CUL3,PBX1,PARVA,PAWR,PODXL,SLFN13,WNK2,PLA2G4A,DNTTIP2,ITGA1,COL5A1,SPATS2L,P2RX6,TEK,GSTM1,HAL,TANC1,OTOF,SOX5,CAMK1D,NSF,PLXNC1 | 1.135002 | 9.03E-07 | 0.002097 |
| GO:0044424 | intracellular part | cellular_component | RHOU,GDA,ZFP64,TSNARE1,ATP5G2,MNAT1,ZNF808,FASN,ST8SIA6,MYL6,PDE4DIP,NBPF20,NBPF9,PSMD2,FEN1,DISC1,SHANK2,ZNF107,KRTAP10-11,TRIM4,PCBP2,MAP2K6,CCDC86,PLXDC1,PRIM2,SPRR2D,BRD1,RGS7,RAD18,SYNGAP1,SEPT7,RNF7,VPS54,ZNF211,UBE2V2,RBPJL,PPP1R11,EMC2,KLC4,MRPL2,SLC35A4,APBB3,SSH2,MPHOSPH10,MCM10,FANCC,NAPG,DUSP5,MACF1,E2F3,INO80,TMEM231,STOX1,NR3C1,ZHX3,METAP1,CALN1,AGBL4,SVIL,NDUFB1,CEP85L,MEIS3,HSPA5,RASA3,MAP2K3,ERP44,RGL1,SGK1,FBXO25,ATP1B3,PELI2,ABL2,ROR1,PTP4A1,RAD54L2,SUMF2,CCT6A,PDCD6,METTL5,RPGRIP1,SPTA1,CSDE1,NRAS,TSEN2,KRT28,PSMA3,RNF19A,MYO5A,JMJD1C,ANAPC10,SRSF4,MTUS1,CRYBA1,SLC25A15,PELP1,FAM120A,RAB6A,MRPS31,XRCC2,NBEA,IPO11,SNX2,MANBA,DLG1,EHD3,AIM2,NLK,RPSAP58,TMEM109,GMPR,DEFA4,WFS1,RPL26L1,GOLGA8B,ZNF138,ZNF264,NBPF10,PRPF3,GALNT5,HNRNPF,APBB2,DCT,HADHB,SKAP2,SH3GL1,TJP1,TNKS2,NBPF8,LDHAL6A,CYP4F12,RNF14,UGGT2,ARL17A,LATS2,CYP4F11,RGMA,GTF2IRD1,PIP5K1A,NLRP12,EIF2AK2,RNASET2,DAZAP1,EEA1,L3MBTL4,PTPRN2,PANX2,AK8,SNX27,CARD11,NF1,ESRRG,NUP160,PABPC1,ATXN2L,STRADA,SLC39A2,ATF6,SYNC,SMARCAD1,CRYGD,WNK1,ELK3,ASAP1,OSBPL3,DNAH17,EYA3,TRIM14,SPSB4,MAP3K8,SYT1,RALBP1,LHX9,ZNF714,SPG7,FZD1,BPTF,TRIM8,MICA,ZNF492,PARP12,HLA-DPA1,TECR,MYBPC1,SLC27A1,RAD51B,BAIAP2L1,OXR1,ZSCAN22,TBC1D1,LMTK2,TCF7L1,ISX,PKD1L1,AGBL1,DPYD,PSD3,HELZ,CYLD,OSTF1,GTF3C4,DDX31,SERPINB2,ITCH,APOL6,PLAGL1,JPH3,CRADD,EGLN1,SMC5,MYT1L,MRPL28,CDC40,SORBS1,KRTAP3-1,BRSK2,BHLHE41,FAT1,PPP2R2C,LOX,FRMD4A,SERPINE2,MYO16,UGT2B17,DOCK2,TSHZ2,ELMO1,ABR,MIER1,UBASH3B,ERBB4,CTSH,UFSP2,PACS1,PARVB,ADAP1,ST8SIA5,SLC8A1,DAAM1,TESC,HTRA1,SGMS1,RYR3,SLK,TBCD,LGALS8,RPTOR,SLC25A21,AKR1B1,DDX60,FNDC1,UGT2B15,ATG4D,GNG7,RPS6KA2,PASK,MED13L,DUSP3,USP15,IL26,SH3BP4,HOOK2,MGAT1,ALOX15B,ZCCHC7,GRB10,MYO1D,FAM120B,TDH,PXDNL,ADRA1B,ZMIZ1,VWF,CNST,EEF1E1,YPEL1,FARP1,AHRR,HDAC4,BANP,CUL2,IL22RA2,SCD5,PPP2R1A,AK3,LHPP,PAK4,PEX14,RPS6KB1,PGM2L1,ST3GAL6,GNAL,FNDC3B,TUBA3E,PRDM2,PITRM1,TSHZ1,CUL3,PBX1,PARVA,PAWR,PODXL,WNK2,PLA2G4A,DNTTIP2,ITGA1,COL5A1,SPATS2L,P2RX6,TEK,GSTM1,HAL,TANC1,OTOF,SOX5,CAMK1D,NSF | 1.13267 | 3.74E-06 | 0.005795 |
| GO:0009898 | internal side of plasma membrane | cellular_component | RGS7,SYNGAP1,RASA3,PTP4A1,SPTA1,DLG1,NF1,CYLD,GNG7,FARP1,GNAL | 4.71533 | 2.15E-05 | 0.025002 |
| GO:0006793 | phosphorus metabolic process | biological_process | RHOU,GDA,ATP5G2,MNAT1,EFTUD1,FASN,MYL6,MAP2K6,RGS7,SYNGAP1,PPP1R11,SSH2,MPHOSPH10,DUSP5,MACF1,INO80,NDUFB1,HSPA5,RASA3,MAP2K3,RGL1,SGK1,ATP1B3,PELI2,ABL2,ROR1,PTP4A1,MLKL,NRAS,NUDT5,RAB6A,XRCC2,DLG1,EHD3,NLK,GMPR,WFS1,DDX59,HADHB,DENND3,UGGT2,ECT2L,LATS2,ATP2B4,PIP5K1A,ZCCHC9,NLRP12,EIF2AK2,PTPRN2,OXER1,AK8,CARD11,NF1,STRADA,ATF6,SMARCAD1,WNK1,ASAP1,DNAH17,EYA3,MAP3K8,RALBP1,FZD1,BPTF,TECR,SLC27A1,RAD51B,TBC1D1,LMTK2,DPYD,PSD3,DDX31,TBC1D22A,BRSK2,PPP2R2C,DOCK2,ABR,UBASH3B,ERBB4,SRMS,ADAP1,TESC,SGMS1,SLK,TBCD,RPTOR,DDX60,DHX37,FNDC1,GNG7,RPS6KA2,PASK,DUSP3,USP15,SH3BP4,MGAT1,RAMP1,GRB10,MYO1D,CNST,FARP1,IL22RA2,PPP2R1A,AK3,LHPP,PAK4,RPS6KB1,PGM2L1,GNAL,TUBA3E,WNK2,PLA2G4A,ITGA1,TEK,CAMK1D,NSF | 1.390238 | 4.81E-05 | 0.044733 |
| GO:0006796 | phosphate-containing compound metabolic process | biological_process | RHOU,GDA,ATP5G2,MNAT1,EFTUD1,MYL6,MAP2K6,RGS7,SYNGAP1,PPP1R11,SSH2,MPHOSPH10,DUSP5,MACF1,INO80,NDUFB1,HSPA5,RASA3,MAP2K3,RGL1,SGK1,ATP1B3,PELI2,ABL2,ROR1,PTP4A1,MLKL,NRAS,NUDT5,RAB6A,XRCC2,DLG1,EHD3,NLK,GMPR,WFS1,DDX59,HADHB,DENND3,UGGT2,ECT2L,LATS2,ATP2B4,PIP5K1A,ZCCHC9,NLRP12,EIF2AK2,PTPRN2,OXER1,AK8,CARD11,NF1,STRADA,ATF6,SMARCAD1,WNK1,ASAP1,DNAH17,EYA3,MAP3K8,RALBP1,FZD1,BPTF,SLC27A1,RAD51B,TBC1D1,LMTK2,DPYD,PSD3,DDX31,TBC1D22A,BRSK2,PPP2R2C,DOCK2,ABR,UBASH3B,ERBB4,SRMS,ADAP1,TESC,SGMS1,SLK,TBCD,RPTOR,DDX60,DHX37,FNDC1,GNG7,RPS6KA2,PASK,DUSP3,USP15,SH3BP4,MGAT1,RAMP1,GRB10,MYO1D,CNST,FARP1,IL22RA2,PPP2R1A,AK3,LHPP,PAK4,RPS6KB1,PGM2L1,GNAL,TUBA3E,WNK2,PLA2G4A,ITGA1,TEK,CAMK1D,NSF | 1.383981 | 7.09E-05 | 0.047264 |
| GO:0048583 | regulation of response to stimulus | biological_process | RHOU,AGTR1,DISC1,PCBP2,MAP2K6,RGS7,SYNGAP1,UBE2V2,DUSP5,MACF1,SNX14,MEIS3,HSPA5,RASA3,MAP2K3,RGL1,PELI2,NRAS,MYO5A,AIM2,NLK,SKAP2,TNKS2,CYP4F12,RNF14,ECT2L,LATS2,RGMA,PIP5K1A,NLRP12,CARD11,NF1,UNC5B,ASAP1,TNIP3,EYA3,MAP3K8,RALBP1,FZD1,MICA,HLA-DPA1,TBC1D1,TCF7L1,PSD3,CYLD,TBC1D22A,SERPINB2,ITCH,SORBS1,BRSK2,SERPINE2,DOCK2,ELMO1,ABR,MIER1,ERBB4,CTSH,PACS1,ADAP1,HTRA1,RPTOR,DDX60,GNG7,RPS6KA2,DUSP3,USP15,IL26,IL17RA,SH3BP4,RAMP1,GRB10,IL4R,ADRA1B,TMEM9B,MASP1,EEF1E1,YPEL1,FARP1,IL22RA2,PPP2R1A,RPS6KB1,CUL3,PAWR,WNK2,ITGA1,TEK,CAMK1D | 1.480637 | 7.12E-05 | 0.047264 |
| GO:0035556 | intracellular signal transduction | biological_process | RHOU,AGTR1,PSMD2,FEN1,DISC1,SHANK2,MAP2K6,RGS7,SYNGAP1,RNF7,DUSP5,MEIS3,RASA3,MAP2K3,RGL1,PELI2,NRAS,PSMA3,MYO5A,RAB6A,NLK,TMEM109,APBB2,TJP1,ARL17A,ECT2L,LATS2,PIP5K1A,NLRP12,CARD11,NF1,WNK1,ASAP1,TNIP3,SPSB4,MAP3K8,RALBP1,TBC1D1,PSD3,CYLD,TBC1D22A,ITCH,CRADD,BRSK2,SERPINE2,MYO16,DOCK2,ELMO1,ABR,MIER1,ERBB4,CTSH,ADAP1,SLC8A1,RPTOR,DDX60,RPS6KA2,DUSP3,IL26,SH3BP4,ADRA1B,TMEM9B,EEF1E1,YPEL1,FARP1,HDAC4,CUL2,IL22RA2,PPP2R1A,RPS6KB1,CUL3,WNK2,ITGA1,TEK | 1.537601 | 8.82E-05 | 0.051242 |
| GO:0005829 | cytosol | cellular_component | RHOU,GDA,FASN,MYL6,PSMD2,PCBP2,MAP2K6,RGS7,FANCC,NR3C1,AGBL4,MAP2K3,RGL1,PELI2,ABL2,CCT6A,SPTA1,PSMA3,ANAPC10,RAB6A,NBEA,DLG1,AIM2,GMPR,RPL26L1,DCT,SKAP2,TJP1,LATS2,PIP5K1A,EIF2AK2,EEA1,AK8,SNX27,CARD11,NUP160,PABPC1,STRADA,SYNC,OSBPL3,MAP3K8,RALBP1,MYBPC1,BAIAP2L1,AGBL1,DPYD,CYLD,ITCH,EGLN1,SORBS1,SERPINE2,MYO16,DOCK2,ELMO1,ABR,ERBB4,CTSH,PACS1,PARVB,HTRA1,RPTOR,AKR1B1,RPS6KA2,DUSP3,IL26,GRB10,EEF1E1,FARP1,HDAC4,CUL2,IL22RA2,PPP2R1A,LHPP,RPS6KB1,PGM2L1,PARVA,PLA2G4A,GSTM1,HAL,OTOF,NSF | 1.492016 | 1.12E-04 | 0.057621 |
| GO:0009117 | nucleotide metabolic process | biological_process | RHOU,GDA,ATP5G2,MNAT1,EFTUD1,MYL6,RGS7,SYNGAP1,MACF1,INO80,HSPA5,RASA3,RGL1,ATP1B3,NRAS,NUDT5,RAB6A,XRCC2,DLG1,EHD3,GMPR,DDX59,DENND3,UGGT2,ECT2L,ATP2B4,PIP5K1A,PTPRN2,OXER1,AK8,CARD11,NF1,SMARCAD1,ASAP1,DNAH17,RALBP1,BPTF,RAD51B,TBC1D1,DPYD,PSD3,DDX31,TBC1D22A,DOCK2,ABR,ADAP1,TBCD,DDX60,DHX37,GNG7,SH3BP4,MGAT1,RAMP1,MYO1D,FARP1,AK3,GNAL,TUBA3E,NSF | 1.602951 | 1.76E-04 | 0.059269 |
| GO:0044248 | cellular catabolic process | biological_process | RHOU,GDA,MNAT1,EFTUD1,MYL6,PSMD2,FEN1,DISC1,PCBP2,RGS7,SYNGAP1,UBE2V2,MACF1,INO80,HSPA5,RASA3,RGL1,ATP1B3,ABL2,CSDE1,NRAS,PSMA3,ANAPC10,NUDT5,RAB6A,XRCC2,MANBA,EHD3,WFS1,RPL26L1,DDX59,DAP,HADHB,CYP4F12,DENND3,ECT2L,ATP2B4,PIP5K1A,RNASET2,PTPRN2,GPX3,NF1,PABPC1,SMARCAD1,ASAP1,DNAH17,RALBP1,BPTF,RAD51B,OXR1,TBC1D1,DPYD,PSD3,CYLD,DDX31,TBC1D22A,ITCH,DOCK2,ABR,CTSH,ADAP1,TBCD,SLC25A21,DDX60,DHX37,ATG4D,USP15,SH3BP4,MGAT1,MYO1D,PXDNL,ADRA1B,FARP1,HDAC4,CUL2,PPP2R1A,GNAL,TUBA3E,CUL3,PLA2G4A,GSTM1,HAL,NSF | 1.459117 | 1.81E-04 | 0.059269 |
| GO:0007009 | plasma membrane organization | biological_process | SPTA1,MYO5A,DLG1,EHD3,PIP5K1A,SORBS1,PACS1,TESC,RAMP1,COL5A1 | 4.013199 | 1.95E-04 | 0.059269 |
| GO:0006753 | nucleoside phosphate metabolic process | biological_process | RHOU,GDA,ATP5G2,MNAT1,EFTUD1,MYL6,RGS7,SYNGAP1,MACF1,INO80,HSPA5,RASA3,RGL1,ATP1B3,NRAS,NUDT5,RAB6A,XRCC2,DLG1,EHD3,GMPR,DDX59,DENND3,UGGT2,ECT2L,ATP2B4,PIP5K1A,PTPRN2,OXER1,AK8,CARD11,NF1,SMARCAD1,ASAP1,DNAH17,RALBP1,BPTF,RAD51B,TBC1D1,DPYD,PSD3,DDX31,TBC1D22A,DOCK2,ABR,ADAP1,TBCD,DDX60,DHX37,GNG7,SH3BP4,MGAT1,RAMP1,MYO1D,FARP1,AK3,GNAL,TUBA3E,NSF | 1.596774 | 1.96E-04 | 0.059269 |
| GO:0031235 | intrinsic to internal side of plasma membrane | cellular_component | SYNGAP1,RASA3,SPTA1,NF1 | 12.61504 | 2.17E-04 | 0.059269 |

### Supplementary Table S6-b.

**The top 15 significant signatures related to psychosis risk syndrome in the GO category of differentially methylated probes.**

| GO ID | GO term | GO category | Symbols in list | Fold enrichment | *P* value | FDR |
| --- | --- | --- | --- | --- | --- | --- |
| GO:0005737 | cytoplasm | cellular_component | STOX1,CEP85L,TSEN2,SHANK2,KIF23,CCT6A,SSH2,MTFR2,ATAD3A,KRTAP6-1,RCC2,MYL6,METTL5,CUL7,MRPL2,HCFC2,PELP1,MCM10,NDUFB1,RNF7,PCBP2,MNAT1,PTP4A1,MFSD2A,SEPT7,TAF6,MYO3B,SLC33A1,PSMA3,ATP1B3,MACF1,ST8SIA6,TUBB6,UBTD2,HSPA1L,FZD5,EMC2,WEE2,GPN1,DDX60,AGPAT3,CHMP3,MCF2L,KLHL17,DTX2,KLC4,GABARAPL3,ATP5G2,PRPF3,C6orf203,ACTN1,PPP1R11,SVIL,MTCH1,FANCC,ROR1,MTERFD2,FASN,ARHGAP22,ERP44,TJP1,SPRR2D,SERAC1,EYA3,S100A7A,PPP2R5A,PPP2R2D,UBE2V2,FBXL12,VPS54,MYO5A,GDA,TRAK2,DCT,ABL2,CUL2,C1orf198,SGCZ,RNF19A,AFAP1,CALN1,ARHGAP5,EHD3,CCDC88B,SLC25A15,FAF1,UBQLN4,TXNDC12,PHACTR1,NUP210,FBXO22,RAB6A,KLHDC3,MICA,NLK,FOLH1B,RHOU,LIMK2,SNX19,HERC2,MTUS2,ZHX3,NAPG,CYP2R1,HSPB6,TSNARE1,SCYL1,MRPS31,UBB,SLC16A1,IGF2BP1,KATNAL2,NDUFB6,PLXDC1,UGT1A3,UGT1A5,UGT1A4,UGT1A8,UGT1A10,UGT1A9,UGT1A7,UGT1A6,SLC35A4,APBB3,AIM2,TEKT2,LYST,MAP2K3,TNRC6C,MAPT,MDM2,GUCY1A2,ADH1C,FAM120A,AOC2,STX8,FRMD4A,CTNNA1,CLIC6,PRKG1,SLC25A17,SH3GL1,NCAPD2,RPS15,DECR2,SP7,DISC1,GLS,FEN1,NR3C1,PEX19,RSU1,TNKS2,ARNTL2,IFI35,SYNE2,A1CF,PLG,PCDH15,MAP4,SSR1,ATP2A2,LDLRAP1,GLT1D1,PABPC1,GGNBP2,PELI2,FUT8,NGLY1,ALG14,CDC20,DST,MKL2,GDI2,PHOSPHO1,RACGAP1,PSMD2,TRIM4,HSF2BP,PLBD2,VAV2,SPIRE1,PDE4DIP,NBPF20,NBPF9,PIP5K1A,SEC16A,KRT28,ABCF2,NOSTRIN,PDCD2L,COL22A1,VPS52,FADS3,TIA1,ATCAY,MAP2K6,SULT1A2,RANBP2,UVRAG,STON2,ARL8B,SLC25A28,ULK2,CCDC40,NBPF8,ALG6,CAB39,GCAT,DBNDD1,HMBS,ADCY10,NEK7,PIGY,PDSS1,HSPA5,SNAI1,AK3,VAC14,DNAH14,TMEM98,ARL2BP,AIRE,PTPN12,RAB3GAP1,RAB43,EMILIN3,OPTN,METAP1,PARG,FLYWCH1,OAS3,TPM4,HOOK3,MYRIP,TRIM26,USP47,PDE8B,PTPN9,PABPC3,SMAD2,TNFAIP8,DLG4,MESDC2,ARHGEF12,MT1F,ANO1,TRIM71,HECW1,SMG6,SYNGAP1,ADCK5,MRPL16,RASA2,ANP32E,JMY,KARS,TERF2IP,PKM,EHBP1,UMPS,RABGAP1L,ZP3,KANK1,NEDD9,IFT80,ALKBH3,TNIP2,TMOD3,SGK1,GAS7,VHL,ARHGAP17,DYSF,MEX3A,SPTA1,TBC1D15,TMED2,MR1,YWHAQ,TUBA3C,LAP3,HNRNPD,RBFOX3,ANTXR2,ASH1L,PPP1R13L,GCH1,UBXN1,POM121,TMED5,RELL1,CDC25C,PDCD6,INVS,EML5,TMEM8B,ELF2,SHOC2,SNRPB,HTT,PRR11,POM121C,CYP3A5,HSPA13,CYP2F1,CYP27A1,PPHLN1,RAB11FIP5,DRG2,MCCC1,TAF4B,NIF3L1,WDR34,LRP5,WIPI2,RAB3A,CRYBA1,LRRC37B,MTERFD3,TRAPPC6B,TRADD,FARP1,NMNAT3,PIGK,COQ7,PRMT7,SLC7A6OS,MADD,C1QL1,RASA3,TMEM109,RGS6,CMIP,MIB1,MRPL21,ATG7,PPCDC,SNCA,ENAH,ABLIM2,MANBA,ARHGAP31,NEIL2,LTA4H,FRMD3,EXOC6,MMADHC,WDR55,FOXP1,GLUL,PRKCA,TNPO1,RPGRIP1L,RELA,CPSF3L,HEATR2,DHFR,COX6C,GCHFR,FAM129A,GNPNAT1,CAMKMT,PI4K2B,HSPA14,REEP1,FLG,CLPB,PRKD2,HK1,ASAP1,ST6GALNAC4,COL27A1,IL37,NDUFB2,SYN3,LACTB2,AGBL4,PLA2G4C,SCIN,RPL32,COBL,RGS7,GSK3B,CDC26,CSDE1,NRAS,RNASEH1,PAN3,GSTA2,USP32,SDCBP2,RTTN,CACNA1S,DLG1,SRPK2,ACOT7,GMPS,UBR5,ACN9,ARPC3,MCOLN3,NUBP1,STK40,CRYM,PATE4,PAIP2B,RBPJ,SIRT2,UBE3C,GPR161,PARN,TIMM8B,SDHD,CLIP1,SH3BGR,PACSIN2,AP3D1,MPHOSPH8,RDM1,NBPF10,TXNL4A,ZNF146,IGF2BP2,PTPRN2,KRT14,CACNA1C,TACC1,ESPN,RPS6KA2,LHPP,ELMO1,TROAP,PRKACG,SUFU,PMP2,PDE3B,FBXL14,RASGRP3,B4GALT5,EPS8,GORASP2,OSBPL5,BRPF3,NUAK1,UGT2B7,UGT2B15,MTUS1,FAHD1,HAGH,PRDX5,ACSF3,ADAM12,SLC35D1,PAPOLA,ARHGAP27,RAB7A,NCOA3,MAPK6,ATXN2L,SPTLC2,CLSTN1,MTIF2,CDC27,VCL,EIF2B5,COX6B1,BRCA1,RANBP6,ARHGAP19,RBM25,RPL3,PLEK,FNTA,SLC30A6,KIDINS220,KIF16B,HYAL1,KIAA1967,PPP3CC,NF1,KPNA1,NXN,NBEA,SCGN,NSFL1C,DHRS2,DSTN,GABRB1,TUBGCP5,API5,SCAPER,QKI,RAB3D,ACER2,TAOK3,SLC23A2,UNC93B1,N4BP2,ATP5A1,KIF26B | 1.124896 | 6.46E-11 | 5.79E-07 |
| GO:0005622 | intracellular | cellular_component | STOX1,CEP85L,TSEN2,SHANK2,ZNF324,KIF23,CCT6A,RAD54L2,SSH2,MTFR2,ATAD3A,KRTAP6-1,RCC2,MYL6,ZNF845,METTL5,CUL7,MRPL2,HCFC2,PELP1,MCM10,NDUFB1,ZBTB45,RNF7,PCBP2,MNAT1,PTP4A1,MFSD2A,SEPT7,TAF6,GRIN2A,PRIM2,MYO3B,SLC33A1,PSMA3,ATP1B3,ORC4,MACF1,JMJD1C,ST8SIA6,PURG,WRN,TUBB6,UBTD2,HSPA1L,FZD5,EMC2,RPGRIP1,WEE2,GPN1,DDX60,AGPAT3,CHMP3,ZNF107,ZFP64,MCF2L,KLHL17,DTX2,KLC4,GABARAPL3,ATP5G2,ZNF211,PRPF3,C6orf203,KRTAP10-11,ACTN1,ZNF605,PPP1R11,SVIL,FBXO25,MTCH1,FANCC,ROR1,MTERFD2,RBPJL,FASN,RBM39,BANP,GNL2,CCDC86,RAD18,ARHGAP22,ERP44,TJP1,SPRR2D,SERAC1,GTF2H5,EYA3,S100A7A,TET1,PPP2R5A,PPP2R2D,KLF3,ZNF628,STAG1,UBE2V2,FBXL12,VPS54,MYO5A,SBF1,GDA,TRAK2,EZH2,RBMS1,DCT,ABL2,CUL2,DDX11,C1orf198,SGCZ,RNF19A,AFAP1,CALN1,ARHGAP5,HIST3H3,EHD3,CCDC88B,SLC25A15,HIST3H2BB,HIST3H2A,CLUAP1,FAF1,UBQLN4,TXNDC12,PHACTR1,NUP210,FBXO22,SMARCC1,BRD1,RAB6A,KLHDC3,MICA,NLK,FOLH1B,RHOU,LIMK2,SNX19,PPFIBP2,SRSF4,HERC2,MTUS2,CARD16,ZHX3,NAPG,CYP2R1,HSPB6,TSNARE1,SCYL1,MRPS31,UBB,GTF2A2,SLC16A1,ZNF431,IGF2BP1,KATNAL2,NDUFB6,PLXDC1,UGT1A3,UGT1A5,UGT1A4,UGT1A8,UGT1A10,UGT1A9,UGT1A7,UGT1A6,SLC35A4,APBB3,AIM2,TEKT2,LYST,MAP2K3,TRIM43,TNRC6C,MAPT,MDM2,GUCY1A2,ADH1C,RPF1,FAM120A,AOC2,STX8,FRMD4A,INO80,CTNNA1,CLIC6,PRKG1,SLC25A17,SH3GL1,RB1,NCAPD2,RPS15,DECR2,SP7,DISC1,GLS,FEN1,NR3C1,PEX19,RSU1,RRN3,TNKS2,ARNTL2,IFI35,ZNF610,SYNE2,A1CF,PLG,PCDH15,MAP4,ZNF607,SSR1,ATP2A2,LDLRAP1,GLT1D1,PABPC1,GGNBP2,TRPS1,PELI2,PPIE,FUT8,NGLY1,ALG14,CDC20,DST,MKL2,GDI2,ZNF888,NMNAT1,PHOSPHO1,RACGAP1,PSMD2,TRIM4,HSF2BP,PLBD2,VAV2,SPIRE1,PDE4DIP,NBPF20,NBPF9,PIP5K1A,SEC16A,KRT28,TBC1D26,ABCF2,NOSTRIN,PDCD2L,COL22A1,VPS52,FADS3,DIP2A,TIA1,ATCAY,ZNF808,MAP2K6,SULT1A2,TFAP2E,RANBP2,MYT1L,UVRAG,STON2,ARL8B,SLC25A28,DUSP5,ULK2,ZNF701,CCDC40,HSPA6,NBPF8,ALG6,CAB39,GCAT,DBNDD1,MBD5,HMBS,ADCY10,GTF3C6,E2F3,NEK7,SUMO4,PIGY,PDSS1,HSPA5,CHFR,ZNF195,SNAI1,AK3,VAC14,DNAH14,FCF1,TMEM98,ARL2BP,AIRE,PTPN12,KLF2,RAB3GAP1,RAB43,HIST1H2BA,HIST1H2AA,EMILIN3,OPTN,ZNF521,METAP1,PARG,PRKDC,FLYWCH1,OAS3,TPM4,HOOK3,ZNF433,ZNF485,SMUG1,MYRIP,TRIM26,USP47,ZNF28,PDE8B,PTPN9,ZNF354B,PABPC3,SMAD2,TNFAIP8,DLG4,MESDC2,FAM188A,ARHGEF12,MT1F,ANO1,TRIM71,HECW1,SMG6,SYNGAP1,KRTAP1-4,ADCK5,MRPL16,FANCD2,RASA2,ANP32E,JMY,C11orf49,ZNF200,KARS,TERF2IP,SCMH1,PKM,EHBP1,UMPS,MEIS3,CTTNBP2NL,RABGAP1L,ZP3,KANK1,NEDD9,IFT80,ALKBH3,ZNF486,TNIP2,TMOD3,SGK1,GAS7,TMEM231,VHL,ARHGAP17,DYSF,MEX3A,SPTA1,TBC1D15,TMED2,MR1,ZNF177,YWHAQ,TUBA3C,ZNF429,LAP3,HNRNPD,RBFOX3,IGF1R,ANTXR2,ASH1L,PPP1R13L,GCH1,UBXN1,POM121,TMED5,RELL1,CDC25C,PDCD6,INVS,EML5,TMEM8B,ELF2,SHOC2,SNRPB,HTT,ZNF611,PRR11,POM121C,FSD1L,FNBP4,CYP3A5,HSPA13,NCOA6,ZNF716,CYP2F1,CYP27A1,PPHLN1,RAB11FIP5,DRG2,MCCC1,UBA2,TAF4B,NIF3L1,WDR34,LRP5,WIPI2,RAB3A,CRYBA1,TFDP2,EP400,LRRC37B,MTERFD3,ZNF876P,TRAPPC6B,TRADD,FARP1,NMNAT3,CCDC59,PIGK,COQ7,PRMT7,SLC7A6OS,MADD,C1QL1,RASA3,TMEM109,RGS6,CMIP,MIB1,MRPL21,ATG7,FAM32A,CARD6,PPCDC,DUXA,SNCA,ZNF257,KDM2A,ENAH,WDR36,ABLIM2,MANBA,ARHGAP31,ZNF91,NEIL2,LTA4H,DCAF4,FRMD3,EXOC6,ZNF747,CHD7,MMADHC,WDR55,FOXP1,EXD3,GLUL,PRKCA,TNPO1,RPGRIP1L,RELA,CPSF3L,HEATR2,DHFR,COX6C,GCHFR,GTF3A,ANKRD42,ZNF587B,FAM129A,GNPNAT1,CAMKMT,PI4K2B,MPHOSPH10,HSPA14,REEP1,HN1,FERMT3,XRCC2,FLG,RSBN1L,CLPB,DIP2C,VPS72,HP1BP3,PRKD2,HK1,ASAP1,ST6GALNAC4,COL27A1,IL37,NDUFB2,SYN3,CMAS | 1.076695 | 1.48E-10 | 6.20E-07 |
| GO:0044424 | intracellular part | cellular_component | STOX1,CEP85L,TSEN2,SHANK2,ZNF324,KIF23,CCT6A,RAD54L2,SSH2,MTFR2,ATAD3A,KRTAP6-1,RCC2,MYL6,ZNF845,METTL5,CUL7,MRPL2,HCFC2,PELP1,MCM10,NDUFB1,ZBTB45,RNF7,PCBP2,MNAT1,PTP4A1,MFSD2A,SEPT7,TAF6,GRIN2A,PRIM2,MYO3B,SLC33A1,PSMA3,ATP1B3,ORC4,MACF1,JMJD1C,ST8SIA6,PURG,WRN,TUBB6,UBTD2,HSPA1L,FZD5,EMC2,RPGRIP1,WEE2,GPN1,DDX60,AGPAT3,CHMP3,ZNF107,ZFP64,MCF2L,KLHL17,DTX2,KLC4,GABARAPL3,ATP5G2,ZNF211,PRPF3,C6orf203,KRTAP10-11,ACTN1,ZNF605,PPP1R11,SVIL,FBXO25,MTCH1,FANCC,ROR1,MTERFD2,RBPJL,FASN,RBM39,BANP,GNL2,CCDC86,RAD18,ARHGAP22,ERP44,TJP1,SPRR2D,SERAC1,GTF2H5,EYA3,S100A7A,TET1,PPP2R5A,PPP2R2D,KLF3,ZNF628,STAG1,UBE2V2,FBXL12,VPS54,MYO5A,SBF1,GDA,TRAK2,EZH2,RBMS1,DCT,ABL2,CUL2,DDX11,C1orf198,SGCZ,RNF19A,AFAP1,CALN1,ARHGAP5,HIST3H3,EHD3,CCDC88B,SLC25A15,HIST3H2BB,HIST3H2A,CLUAP1,FAF1,UBQLN4,TXNDC12,PHACTR1,NUP210,FBXO22,SMARCC1,BRD1,RAB6A,KLHDC3,MICA,NLK,FOLH1B,RHOU,LIMK2,SNX19,SRSF4,HERC2,MTUS2,ZHX3,NAPG,CYP2R1,HSPB6,TSNARE1,SCYL1,MRPS31,UBB,GTF2A2,SLC16A1,ZNF431,IGF2BP1,KATNAL2,NDUFB6,PLXDC1,UGT1A3,UGT1A5,UGT1A4,UGT1A8,UGT1A10,UGT1A9,UGT1A7,UGT1A6,SLC35A4,APBB3,AIM2,TEKT2,LYST,MAP2K3,TNRC6C,MAPT,MDM2,GUCY1A2,ADH1C,RPF1,FAM120A,AOC2,STX8,FRMD4A,INO80,CTNNA1,CLIC6,PRKG1,SLC25A17,SH3GL1,RB1,NCAPD2,RPS15,DECR2,SP7,DISC1,GLS,FEN1,NR3C1,PEX19,RSU1,RRN3,TNKS2,ARNTL2,IFI35,ZNF610,SYNE2,A1CF,PLG,PCDH15,MAP4,ZNF607,SSR1,ATP2A2,LDLRAP1,GLT1D1,PABPC1,GGNBP2,TRPS1,PELI2,PPIE,FUT8,NGLY1,ALG14,CDC20,DST,MKL2,GDI2,ZNF888,NMNAT1,PHOSPHO1,RACGAP1,PSMD2,TRIM4,HSF2BP,PLBD2,VAV2,SPIRE1,PDE4DIP,NBPF20,NBPF9,PIP5K1A,SEC16A,KRT28,ABCF2,NOSTRIN,PDCD2L,COL22A1,VPS52,FADS3,DIP2A,TIA1,ATCAY,ZNF808,MAP2K6,SULT1A2,TFAP2E,RANBP2,MYT1L,UVRAG,STON2,ARL8B,SLC25A28,DUSP5,ULK2,ZNF701,CCDC40,HSPA6,NBPF8,ALG6,CAB39,GCAT,DBNDD1,MBD5,HMBS,ADCY10,GTF3C6,E2F3,NEK7,SUMO4,PIGY,PDSS1,HSPA5,CHFR,ZNF195,SNAI1,AK3,VAC14,DNAH14,FCF1,TMEM98,ARL2BP,AIRE,PTPN12,KLF2,RAB3GAP1,RAB43,HIST1H2BA,HIST1H2AA,EMILIN3,OPTN,ZNF521,METAP1,PARG,PRKDC,FLYWCH1,OAS3,TPM4,HOOK3,ZNF433,ZNF485,SMUG1,MYRIP,TRIM26,USP47,ZNF28,PDE8B,PTPN9,ZNF354B,PABPC3,SMAD2,TNFAIP8,DLG4,MESDC2,FAM188A,ARHGEF12,MT1F,ANO1,TRIM71,HECW1,SMG6,SYNGAP1,KRTAP1-4,ADCK5,MRPL16,FANCD2,RASA2,ANP32E,JMY,C11orf49,ZNF200,KARS,TERF2IP,SCMH1,PKM,EHBP1,UMPS,MEIS3,CTTNBP2NL,RABGAP1L,ZP3,KANK1,NEDD9,IFT80,ALKBH3,ZNF486,TNIP2,TMOD3,SGK1,GAS7,TMEM231,VHL,ARHGAP17,DYSF,MEX3A,SPTA1,TBC1D15,TMED2,MR1,ZNF177,YWHAQ,TUBA3C,ZNF429,LAP3,HNRNPD,RBFOX3,IGF1R,ANTXR2,ASH1L,PPP1R13L,GCH1,UBXN1,POM121,TMED5,RELL1,CDC25C,PDCD6,INVS,EML5,TMEM8B,ELF2,SHOC2,SNRPB,HTT,ZNF611,PRR11,POM121C,FNBP4,CYP3A5,HSPA13,NCOA6,ZNF716,CYP2F1,CYP27A1,PPHLN1,RAB11FIP5,DRG2,MCCC1,UBA2,TAF4B,NIF3L1,WDR34,LRP5,WIPI2,RAB3A,CRYBA1,TFDP2,EP400,LRRC37B,MTERFD3,ZNF876P,TRAPPC6B,TRADD,FARP1,NMNAT3,CCDC59,PIGK,COQ7,PRMT7,SLC7A6OS,MADD,C1QL1,RASA3,TMEM109,RGS6,CMIP,MIB1,MRPL21,ATG7,FAM32A,PPCDC,DUXA,SNCA,ZNF257,KDM2A,ENAH,WDR36,ABLIM2,MANBA,ARHGAP31,ZNF91,NEIL2,LTA4H,DCAF4,FRMD3,EXOC6,CHD7,MMADHC,WDR55,FOXP1,GLUL,PRKCA,TNPO1,RPGRIP1L,RELA,CPSF3L,HEATR2,DHFR,COX6C,GCHFR,GTF3A,ANKRD42,ZNF587B,FAM129A,GNPNAT1,CAMKMT,PI4K2B,MPHOSPH10,HSPA14,REEP1,HN1,FERMT3,XRCC2,FLG,RSBN1L,CLPB,DIP2C,VPS72,HP1BP3,PRKD2,HK1,ASAP1,ST6GALNAC4,COL27A1,IL37,NDUFB2,SYN3,CMAS,LACTB2,AGBL4,ZNF33A,EHMT2,PLA2G4C,SCIN,RPL32,FAM101B | 1.079191 | 2.08E-10 | 6.20E-07 |
| GO:0006796 | phosphate-containing compound metabolic process | biological_process | SSH2,MYL6,CUL7,NDUFB1,MNAT1,PTP4A1,CDK19,MYO3B,ATP1B3,MACF1,PPP1R2P1,WRN,TUBB6,FZD5,WEE2,DDX60,AGPAT3,ERBB3,MCF2L,ATP5G2,PPP1R11,ROR1,GNL2,ARHGAP22,SERAC1,EYA3,PPP2R5A,PPP2R2D,EFTUD1,SBF1,GDA,EZH2,ABL2,DDX11,ARHGAP5,EHD3,FAF1,PHACTR1,RAB6A,KLHDC3,NLK,RHOU,LIMK2,HERC2,SCYL1,UBB,KATNAL2,NDUFB6,LYST,MAP2K3,GUCY1A2,INO80,PRKG1,RB1,EPHA1,SSR1,ATP2A2,PELI2,DST,GDI2,NMNAT1,PHOSPHO1,RACGAP1,VAV2,SBK2,PIP5K1A,TBC1D26,ABCF2,IL34,MAP2K6,SULT1A2,ARL8B,DUSP5,ULK2,CAB39,ADCY10,NEK7,PIGY,HSPA5,AK3,VAC14,ARL2BP,PTPN12,RAB3GAP1,RAB43,PRKDC,SMUG1,PDE8B,PTPN9,C3,SMAD2,ARHGEF12,SMG6,SYNGAP1,ADCK5,ULK4,RASA2,ANP32E,KARS,EPHB4,PKM,UMPS,RABGAP1L,ZP3,SGK1,ARHGAP17,TBC1D15,TMED2,TUBA3C,IGF1R,PPP1R2,GCH1,BMPR2,PTPRU,CDC25C,SHOC2,ADCY9,HTT,TPTE,LRP5,RAB3A,ZCCHC9,HBS1L,FARP1,NMNAT3,MLKL,PIGK,COQ7,MADD,RASA3,RGS6,PPCDC,SNCA,ARHGAP31,ABCC3,PRKCA,CD3E,DHFR,FAM129A,GNPNAT1,PI4K2B,MPHOSPH10,PTPRJ,XRCC2,PRKD2,HK1,ASAP1,NDUFB2,PLA2G4C,RGS7,GSK3B,NRAS,PAN3,ADCY6,DLG1,SRPK2,GMPS,STK40,SIRT2,GPR161,ARHGAP23,PTPRN2,PTPRG,RPS6KA2,LHPP,PRKACG,PDE3B,RASGRP3,EPS8,NUAK1,PRDX5,SLC35D1,ARHGAP27,RAB7A,GMFB,MAPK6,SPTLC2,MTIF2,EIF2B5,SEPHS1,ARHGAP19,RAD51B,GNB3,PLEK,KIDINS220,PPP3CC,NF1,ADARB1,PTPRK,RAB3D,ACER2,TAOK3,N4BP2,ATP5A1,RIN3,ACER3,NUDT5,PLCG2,GRK5,MTMR12,CCNY,TOP2B,MAPKAP1,ARHGEF4,RFC1,SLC44A2,TPX2,IQSEC1,STK3,GRK6,ILKAP,DNMBP,RRAGC,ARHGAP12,RGL1,MINK1,CASC5,GTF2H4,PILRB,SMAD1,LYN,DDX43,DDX59,AGPAT5,AGAP11,LATS2,TTK,S1PR2,NUDT3,PDE6A,SPAG9,CLN8,RORA,ABI2,PAK2,TBC1D16,IRAK4,BPNT1,MCEE,ARHGAP32,HADHB,TBC1D1,KRIT1,IMPA1,RGS9,PDE7A,PTPN11,PPM1D,PRKRA,TTBK2,DYNC1H1,RPTOR,MTMR3,RGS8,SULT1A1,IKBKB,LIMD1,OPA1,KIF1B,SCARB1,IMPACT,RYR2,CALCR,PPP1R14D,PPP6R3,YES1,FASTKD2,CCL21,CAPRIN2,ATP6V0C,RSF1,MUC20,MAP3K2,TEC,HSPA2,UGGT2,GK5,SERTAD1,BUB1,ECT2L,FLT1,NT5C2,AXL,DNAH3,ATP2B4,RAB3GAP2,CHUK,ADCK2,SIK3,MUL1,INPP5K,PPP5C,CMSS1,FRS2,MSH4,RAPGEF6,MYH10,PTEN,RNASEL,TTBK1,WFS1,MAML1,SRGAP1,PPP2R1A,NUDT15,CDK12,NLRP12,EPHA10,MRAP,OBSCN,GTF2H1,DOCK3,DOCK6,TATDN2,PFAS,CERS4,MARK2,CSNK2A1,KALRN,ST5,EGFR,HMGXB3,CTGF,DDX24,RIMBP2,IRAK3,IL12B,RANBP10,ADCK4,ITPKC,IQGAP1,CHRM5,PIK3C2G,NEK10,TBC1D8,ERC1,CERS6,LPIN1,ATP11B,AGPAT4,RASL10B,DENND3,UHMK1,VAV1,PTPRO,SMAP1,TNXB,STK31,RALGDS,TPM1,CCDC88A,GBA2,FDPS,GNA12,ELANE,AK2,C5,ARHGAP42,EPHA5,STK24,GBP1,YWHAG,CCL3,CTBP1,SSH1,TGFA,MGLL,DNAJC6,LONRF2,PFKFB3,MEN1,F2R,PGAM2,TBC1D2B,PTPN14,ATP8A2,AK8,ARSB,BPTF,SET,CEP192,SPTLC1,EPHB1,RBM26,WNK4,ASNS,NAPRT1,DLC1,NEK4,SLK,OXER1,MKNK1,LMTK2,FGFR2,BRSK2,CSGALNACT1,FNDC1,CCDC155,CHKB-CPT1B,DNAH17,GRB10,NEK11,RTEL1-TNFRSF6B,DDX31,ARHGEF37,DCLK1,GNAT2,NGEF,ATP10A,SERINC5,SMARCAD1,FZD1,ARHGAP10,ABCA13,SGSM3,INPP5B,SH3BP4,MAPK1,RPS6KB1,DAB1,ABR,PRTFDC1,LCLAT1,PPP4R2,CDK11B,EYA4,DUSP22,ATP5S,PFKP,ARHGEF10,SLC27A1,MEF2A,SH2D4A,PTPRM,HIBADH,PRKCH,PFKL,PTPN1,DUSP3,ATP11A,EIF2AK4,DBNL,EPHB2,GPHN,PDIA6,CHP2,DOCK1,DPYD,CAT,DAPK2,PGS1,CARKD,NSF,GAB1,FBP1,PRKAG2,PSD4,DAPK1,ATXN1,CNST,PHACTR3,PASK,TEK,GNG7,GNAO1,FBXO8,ITGA1,ARHGAP18,PTPRE,UPP2,PLA2G4A,ASCC3,PRKCE,SFI1,ATL1,P2RX7,IL22RA2,ATP10D,ABCC4,PPP2R2C,ADAP1,NOS1,NRG1,UBASH3B,SGMS1,PGM2L1,ARHGAP30,ATP13A5,STK32C,ATP6V1D | 1.216177 | 1.62E-07 | 3.63E-04 |
| GO:0006793 | phosphorus metabolic process | biological_process | SSH2,MYL6,CUL7,NDUFB1,MNAT1,PTP4A1,CDK19,MYO3B,ATP1B3,MACF1,PPP1R2P1,WRN,TUBB6,FZD5,WEE2,DDX60,AGPAT3,ERBB3,MCF2L,ATP5G2,PPP1R11,ROR1,FASN,GNL2,ARHGAP22,SERAC1,EYA3,PPP2R5A,PPP2R2D,EFTUD1,SBF1,GDA,EZH2,ABL2,DDX11,ARHGAP5,EHD3,FAF1,PHACTR1,RAB6A,KLHDC3,NLK,RHOU,LIMK2,HERC2,SCYL1,UBB,KATNAL2,NDUFB6,LYST,MAP2K3,GUCY1A2,INO80,PRKG1,RB1,EPHA1,SSR1,ATP2A2,PELI2,DST,GDI2,NMNAT1,PHOSPHO1,RACGAP1,VAV2,SBK2,PIP5K1A,TBC1D26,ABCF2,IL34,MAP2K6,SULT1A2,ARL8B,DUSP5,ULK2,CAB39,ADCY10,NEK7,PIGY,HSPA5,AK3,VAC14,ARL2BP,PTPN12,RAB3GAP1,RAB43,PRKDC,SMUG1,PDE8B,PTPN9,C3,SMAD2,ARHGEF12,SMG6,SYNGAP1,ADCK5,ULK4,RASA2,ANP32E,KARS,EPHB4,PKM,UMPS,RABGAP1L,ZP3,SGK1,ARHGAP17,TBC1D15,TMED2,TUBA3C,IGF1R,PPP1R2,GCH1,BMPR2,PTPRU,CDC25C,SHOC2,ADCY9,HTT,TPTE,LRP5,RAB3A,ZCCHC9,HBS1L,FARP1,NMNAT3,MLKL,PIGK,COQ7,MADD,RASA3,RGS6,PPCDC,SNCA,ARHGAP31,ABCC3,PRKCA,CD3E,DHFR,FAM129A,GNPNAT1,PI4K2B,MPHOSPH10,PTPRJ,XRCC2,PRKD2,HK1,ASAP1,NDUFB2,PLA2G4C,RGS7,GSK3B,NRAS,PAN3,ADCY6,DLG1,SRPK2,GMPS,STK40,SIRT2,GPR161,ARHGAP23,PTPRN2,PTPRG,RPS6KA2,LHPP,PRKACG,PDE3B,RASGRP3,EPS8,NUAK1,PRDX5,SLC35D1,ARHGAP27,RAB7A,GMFB,MAPK6,SPTLC2,MTIF2,EIF2B5,SEPHS1,ARHGAP19,RAD51B,GNB3,PLEK,KIDINS220,PPP3CC,NF1,ADARB1,PTPRK,RAB3D,ACER2,TAOK3,N4BP2,ATP5A1,RIN3,ACER3,NUDT5,PLCG2,GRK5,MTMR12,CCNY,TOP2B,MAPKAP1,ARHGEF4,RFC1,SLC44A2,TPX2,IQSEC1,STK3,GRK6,ILKAP,DNMBP,RRAGC,ARHGAP12,RGL1,MINK1,CASC5,GTF2H4,PILRB,SMAD1,LYN,DDX43,DDX59,AGPAT5,AGAP11,LATS2,TTK,S1PR2,NUDT3,PDE6A,SPAG9,CLN8,RORA,ABI2,PAK2,TBC1D16,IRAK4,BPNT1,MCEE,ARHGAP32,HADHB,TBC1D1,KRIT1,IMPA1,RGS9,PDE7A,PTPN11,PPM1D,PRKRA,TTBK2,DYNC1H1,RPTOR,MTMR3,RGS8,SULT1A1,IKBKB,LIMD1,OPA1,KIF1B,SCARB1,IMPACT,RYR2,CALCR,PPP1R14D,PPP6R3,YES1,FASTKD2,CCL21,CAPRIN2,ATP6V0C,RSF1,MUC20,MAP3K2,TEC,HSPA2,UGGT2,GK5,SERTAD1,BUB1,ECT2L,FLT1,NT5C2,AXL,DNAH3,ATP2B4,RAB3GAP2,CHUK,ADCK2,SIK3,MUL1,INPP5K,PPP5C,CMSS1,FRS2,MSH4,RAPGEF6,MYH10,PTEN,RNASEL,TTBK1,WFS1,MAML1,SRGAP1,PPP2R1A,NUDT15,CDK12,NLRP12,EPHA10,MRAP,OBSCN,GTF2H1,DOCK3,DOCK6,TATDN2,PFAS,CERS4,MARK2,CSNK2A1,KALRN,ST5,EGFR,HMGXB3,CTGF,DDX24,RIMBP2,IRAK3,IL12B,RANBP10,ADCK4,ITPKC,IQGAP1,CHRM5,PIK3C2G,NEK10,TBC1D8,ERC1,CERS6,LPIN1,ATP11B,AGPAT4,RASL10B,DENND3,UHMK1,VAV1,PTPRO,SMAP1,TNXB,STK31,RALGDS,TPM1,CCDC88A,GBA2,FDPS,GNA12,ELANE,AK2,C5,ARHGAP42,EPHA5,STK24,GBP1,YWHAG,CCL3,CTBP1,SSH1,TGFA,MGLL,DNAJC6,LONRF2,ACOT1,PFKFB3,MEN1,F2R,PGAM2,TBC1D2B,PTPN14,ATP8A2,AK8,ARSB,BPTF,SET,CEP192,SPTLC1,EPHB1,RBM26,WNK4,ASNS,NAPRT1,DLC1,NEK4,SLK,OXER1,MKNK1,LMTK2,FGFR2,BRSK2,CSGALNACT1,FNDC1,TECR,CCDC155,CHKB-CPT1B,DNAH17,GRB10,NEK11,RTEL1-TNFRSF6B,DDX31,ARHGEF37,DCLK1,GNAT2,NGEF,ATP10A,SERINC5,SMARCAD1,FZD1,ARHGAP10,ABCA13,SGSM3,INPP5B,SH3BP4,MAPK1,RPS6KB1,DAB1,ABR,PRTFDC1,LCLAT1,PPP4R2,CDK11B,EYA4,DUSP22,ATP5S,PFKP,ARHGEF10,SLC27A1,MEF2A,SH2D4A,PTPRM,HIBADH,PRKCH,PFKL,PTPN1,DUSP3,ATP11A,EIF2AK4,DBNL,EPHB2,GPHN,PDIA6,CHP2,DOCK1,DPYD,CAT,DAPK2,PGS1,CARKD,NSF,GAB1,FBP1,PRKAG2,PSD4,DAPK1,ATXN1,CNST,PHACTR3,PASK,TEK,GNG7,GNAO1,FBXO8,ITGA1,ARHGAP18,PTPRE,UPP2,PLA2G4A,ASCC3,PRKCE,SFI1,ATL1,P2RX7,IL22RA2,ATP10D,ABCC4,PPP2R2C,ADAP1,NOS1,NRG1,UBASH3B,SGMS1,PGM2L1,ARHGAP30,ATP13A5,STK32C,ATP6V1D | 1.207962 | 3.54E-07 | 5.02E-04 |
| GO:0008092 | cytoskeletal protein binding | molecular_function | KIF23,SSH2,MYO3B,MACF1,KLHL17,ACTN1,SVIL,FBXO25,MYO5A,ABL2,AFAP1,CCDC88B,PHACTR1,MTUS2,MAPT,INO80,CTNNA1,SYNE2,MAP4,DST,MKL2,RACGAP1,SPIRE1,ATCAY,ARL8B,TPM4,HOOK3,MYRIP,JMY,TMOD3,GAS7,SPTA1,HTT,RAB11FIP5,FARP1,SNCA,ENAH,ABLIM2,NEIL2,FRMD3,REEP1,AGBL4,SCIN,FAM101B,COBL,GSK3B,DLG1,ARPC3,CLIP1,PACSIN2,CACNA1C,ESPN,EPS8,GMFB,VCL,BRCA1,FNTA,DSTN,TUBGCP5,PLEC,MKL1,SPAG9,ABI2,EPB41L5,WASF2,PLS1,KIF1B,FHOD1,MICAL3,CGN,STARD9,MYH10,FLNB,SBDS,OBSCN,WIPF3,EGFR,RANBP10,CROCC,NRCAM,KLHL5,EPB41L4A,DLG5,KCNQ2,TPM1,CCDC88A,SSH1,MYH3,FMN2,TAGLN,RPH3AL,LMTK2,UTRN,NCALD,MYO5B,TLN2,MYO1B,MYH11,ARHGEF10,ACE,DBNL,MYBPC1,XIRP2,MYH15,HOOK2,PALLD,PHACTR3,DAAM1,FHL3,CAPZB,PKNOX2,PRKCE,MICAL2,PAWR,ANK1,SLC8A1 | 1.568697 | 4.03E-07 | 5.02E-04 |
| GO:0043167 | ion binding | molecular_function | EFCAB7,RNF157,ZNF324,KIF23,CCT6A,RAD54L2,ATAD3A,MYL6,ZNF845,MCM10,ZBTB45,RNF7,MNAT1,CDK19,RNF215,SEPT7,GRIN2A,PRIM2,MYO3B,ORC4,MACF1,JMJD1C,ANKMY1,WRN,TUBB6,HSPA1L,WEE2,GPN1,DDX60,EYS,ZNF107,ERBB3,ZFP64,MCF2L,DTX2,ZNF211,ACTN1,ZNF605,ROR1,SNED1,FASN,GNL2,RAD18,EYA3,S100A7A,TET1,ADAMTS2,KLF3,ZNF628,MYO5A,SCUBE1,EFTUD1,SBF1,GDA,DCT,ABL2,DDX11,RNF19A,CALN1,ZDHHC20,ARHGAP5,EHD3,BRD1,RAB6A,NLK,FOLH1B,RHOU,LIMK2,HERC2,ZHX3,CYP2R1,NRXN3,SCYL1,ZNF431,KATNAL2,UGT1A3,UGT1A4,UGT1A8,UGT1A10,UGT1A9,UGT1A7,UGT1A6,MAP2K3,TRIM43,CAPS2,MDM2,GUCY1A2,ADH1C,AOC2,INO80,PRKG1,C1S,SP7,FEN1,NR3C1,TNKS2,ZNF610,EPHA1,PCDH15,ZNF607,ATP2A2,LDLRAP1,TRPS1,PROCA1,ITIH1,NGLY1,DST,GDI2,ZNF888,NMNAT1,PHOSPHO1,RACGAP1,TRIM4,VAV2,ECEL1,SBK2,PIP5K1A,ABCF2,FADS3,ZNF808,EFCAB5,MAP2K6,RANBP2,MYT1L,ARL8B,ULK2,ZNF701,HSPA6,GIMAP7,GCAT,ADCY10,NEK7,PDSS1,HSPA5,CHFR,ZNF195,SNAI1,AK3,DNAH14,ANXA13,AIRE,KLF2,ADAM32,RAB43,ZNF521,METAP1,PRKDC,FLYWCH1,OAS3,TPM4,ZNF433,ZNF485,MYRIP,TRIM26,ZNF28,PDE8B,ZNF354B,SMAD2,FAM188A,ARHGEF12,MT1F,TRIM71,SMG6,ULK4,RASA2,ZNF200,KARS,EPHB4,PKM,ALKBH3,ZNF486,SGK1,MEX3A,SPTA1,ZNF177,TUBA3C,ZNF429,LAP3,IGF1R,ANTXR2,ASH1L,GCH1,BMPR2,PDCD6,KCNT2,CDH5,ADCY9,ZNF611,CYP3A5,HSPA13,ZNF716,CYP2F1,CYP27A1,DRG2,MCCC1,CDH18,UBA2,ADPRHL1,WIPI2,RAB3A,EP400,ZCCHC9,ZNF876P,HBS1L,FARP1,NMNAT3,SLFN12L,MLKL,COQ7,MADD,RASA3,MIB1,SNCA,SH3RF3,ZNF257,KDM2A,ABLIM2,ZNF91,NEIL2,LTA4H,ZNF747,ABCC3,CHD7,FOXP1,GLUL,PRKCA,RELA,CSAD,GCHFR,GTF3A,ZNF587B,RNF145,PI4K2B,HSPA14,XRCC2,FLG,CLPB,PRKD2,HK1,ASAP1,COL27A1,SYN3,LACTB2,AGBL4,ZNF33A,EHMT2,SCIN,GSK3B,NRAS,RNASEH1,PAN3,ADCY6,ITGAM,USP32,CACNA1S,SRPK2,ACOT7,ZNF83,GMPS,MARCH7,UBR5,NUBP1,STK40,KDM4C,ASXL2,SIRT2,ZNF709,PARN,TIMM8B,SDHD,CLIP1,ZNF146,CACNA1C,ACTL6A,RPS6KA2,LHPP,PRKACG,RLF,PMP2,PDE3B,RASGRP3,B4GALT5,BRPF3,NUAK1,UGT2B7,UGT2B15,ZNF138,FAHD1,HAGH,PDZD8,ACSF3,ADAM12,PAPOLA,RAB7A,PRDM16,KIAA2018,MAPK6,ZNF311,SPTLC2,CLSTN1,MTIF2,EIF2B5,SEPHS1,RNF38,BRCA1,ZNF3,ZNF420,RAD51B,PLEK,KIF16B,ZNF43,PPP3CC,NWD1,ZC3H3,ADARB1,FGF12,SCGN,SCAPER,RAB3D,TAOK3,MMP28,N4BP2,ATP5A1,TNS3,KIF26B,PAPPA2,RIN3,NUDT5,LOXL2,TRIM39,ZNF717,PTGES,TRIM36,FRMPD2,EGLN3,EFCAB9,ZNF813,ZFYVE9,AMDHD2,GRK5,KDM4A,CYP19A1,HS3ST5,TOP2B,MAPKAP1,STAT5B,ARHGEF4,RFC1,ZNF662,TPX2,IQSEC1,STK3,GRK6,FOXK1,ILKAP,DNMBP,RRAGC,TPD52,RGL1,APOBEC4,ZNF578,MINK1,SMAD1,ZSCAN5B,LYN,DDX43,ZNF268,DDX59,AGAP11,ZNF606,LATS2,TTK,ARL15,GSTM4,NFX1,FAT2,CPA5,ZNF148,NUDT3,PDE6A,CBLB,ZNF438,RORA,PAK2,GALNT3,EFCAB14,ENTPD1,ZNF532,TMEM173,IRAK4,BPNT1,ENPEP,HADHB,ACTG1,PRDM10,IMPA1,PDE7A,PPM1D,TTBK2,HDAC5,DYNC1H1,ZNF638,CDH26,ZNF264,MTMR3,SLC25A24,INTS12,PLS1,ZC3H6,IKBKB,LIMD1,CELA1,ZBTB46,SMYD3,KLF17,BAZ2B,OPA1,KIF1B,SCARB1,RYR2,PLEKHM3,YES1,CSHL1,ZNF254,RSF1,MAP3K2,PHF17,TEC,CA5A,ZNF766,LIMS1,MICAL3,SUMF2,HSPA2,CCBE1,ZNF726,GK5,AGBL2,ZNF608,GATA4,RNF14,PARD3,GLRX5,LIAS,PEX10,MSS51,BUB1,RNF135,ECT2L,ZIC2,FLT1,NT5C2,AXL,CLVS1,EFHB,DNAH3,NUP153,ATP2B4,STARD9,TAF15,UBE2I,GPR12,CHUK,ZNF860,ADCK2,SIK3,REV3L,ACAD10,MYL10,MCTP1,PLCL2,MUL1,PPP5C,NSD1,CMSS1,MSH4,RAPGEF6,MYH10,PTEN,KAT7,RNASEL,ZNF574,ACSS3,TTBK1,THAP4,ZNF541,ANXA3,POTEF,NUDT15,EFHD1,CDH12,CDK12,OTOF,NLRP12,EPHA10,MAN2A2,OBSCN,HECTD1,DOCK3,RERE,DOCK6 | 1.138172 | 4.36E-07 | 5.02E-04 |
| GO:0019899 | enzyme binding | molecular_function | RAD54L2,CUL7,PCBP2,FZD5,RAD18,PPP2R5A,MYO5A,TRAK2,CUL2,FAF1,NLK,HERC2,UGT1A3,UGT1A4,UGT1A8,UGT1A10,UGT1A9,UGT1A7,UGT1A6,MAP2K3,MAPT,MDM2,STX8,SH3GL1,RB1,SP7,PEX19,RRN3,TNKS2,EPHA1,CDC20,RACGAP1,PDE4DIP,PIP5K1A,MAP2K6,RANBP2,CAB39,ADCY10,HSPA5,SNAI1,RAB3GAP1,PRKDC,MYRIP,SMAD2,DLG4,RABGAP1L,ZP3,TNIP2,VHL,UBXN1,CDC25C,SHOC2,CDH5,NCOA6,RAB3A,TRADD,SNCA,PRKCA,CD3E,RELA,GCHFR,PTPRJ,GSK3B,ADCY6,DLG1,KDM4C,SIRT2,PARN,SUFU,PDE3B,RASGRP3,EPS8,RAB7A,MAPK6,CDC27,VCL,BRCA1,GNB3,PLEK,KIF16B,KIAA1967,NBEA,PTPRK,RAB3D,GTF2I,RIN3,EXOC2,MEF2D,GRK5,KDM4A,CASC3,GGA2,CCNY,TOP2B,MAPKAP1,TPX2,SMAD1,LYN,GSTM4,SERPINB6,SPAG9,ABI2,PAK2,TMEM173,PTPN11,SNX3,HDAC5,RPTOR,IKBKB,RYR2,PPP6R3,YES1,ATP6V0C,MAP3K2,PARD3,RAB11FIP2,UBE2I,MUL1,RAPGEF6,PTEN,WFS1,MAML1,SRGAP1,PPP1R16B,SKAP1,DFFB,EGFR,FAF2,NCF2,RANBP9,RANBP10,RAB11FIP3,IQGAP1,ERC1,LPIN1,USP37,UHMK1,WWOX,CCDC88A,NFATC1,ELANE,HLCS,YWHAG,RNF8,FOXO1,NBR1,CEP192,THRB,BICD2,RPH3AL,BRSK2,UTRN,CNTNAP2,S100A1,MYO5B,SGSM3,SH3BP4,MAPK1,MBP,IFNAR2,MEF2A,SH2D4A,PRKCH,PFKL,PTPN1,CAT,CUL3,CYB5A,HDAC4,PRKAG2,CNST,DAAM1,ATF2,PRKCE,SFI1,VWF,SLN,PAWR,ANK1,CRADD | 1.417178 | 4.48E-07 | 5.02E-04 |
| GO:0016791 | phosphatase activity | molecular_function | SSH2,PTP4A1,PPP1R2P1,PPP1R11,EYA3,PPP2R5A,PPP2R2D,SBF1,PHACTR1,PHOSPHO1,DUSP5,PTPN12,PTPN9,ANP32E,PPP1R2,PTPRU,CDC25C,SHOC2,HTT,TPTE,ZCCHC9,FARP1,MPHOSPH10,PTPRJ,DLG1,PTPRN2,PTPRG,LHPP,NUAK1,PLEK,PPP3CC,PTPRK,MTMR12,ILKAP,CASC5,BPNT1,IMPA1,PTPN11,PPM1D,MTMR3,PPP1R14D,PPP6R3,NT5C2,INPP5K,PPP5C,FRS2,PTEN,PPP2R1A,CSNK2A1,RIMBP2,LPIN1,PTPRO,SSH1,DNAJC6,PFKFB3,PGAM2,PTPN14,SET,CEP192,RBM26,LMTK2,CCDC155,INPP5B,PPP4R2,EYA4,DUSP22,MEF2A,SH2D4A,PTPRM,PTPN1,DUSP3,CHP2,FBP1,CNST,PHACTR3,PTPRE,SFI1,PPP2R2C,UBASH3B | 1.720613 | 8.00E-07 | 7.96E-04 |
| GO:0016311 | dephosphorylation | biological_process | SSH2,PTP4A1,PPP1R2P1,PPP1R11,EYA3,PPP2R5A,PPP2R2D,SBF1,PHACTR1,PHOSPHO1,DUSP5,PTPN12,PTPN9,SMG6,ANP32E,PPP1R2,PTPRU,CDC25C,SHOC2,HTT,TPTE,ZCCHC9,FARP1,MPHOSPH10,PTPRJ,DLG1,PTPRN2,PTPRG,LHPP,NUAK1,PLEK,PPP3CC,PTPRK,MTMR12,ILKAP,CASC5,BPNT1,IMPA1,PTPN11,PPM1D,MTMR3,PPP1R14D,PPP6R3,NT5C2,INPP5K,PPP5C,FRS2,PTEN,PPP2R1A,CSNK2A1,RIMBP2,IQGAP1,CHRM5,LPIN1,PTPRO,SSH1,DNAJC6,PFKFB3,PGAM2,PTPN14,SET,CEP192,RBM26,DLC1,LMTK2,CCDC155,INPP5B,PPP4R2,EYA4,DUSP22,MEF2A,SH2D4A,PTPRM,PTPN1,DUSP3,CHP2,FBP1,CNST,PHACTR3,PTPRE,SFI1,PPP2R2C,UBASH3B | 1.686427 | 1.00E-06 | 8.35E-04 |
| GO:0003824 | catalytic activity | molecular_function | TSEN2,KIF23,RAD54L2,SSH2,ATAD3A,MYL6,METTL5,CUL7,NDUFB1,RNF7,MNAT1,PTP4A1,CDK19,PRIM2,MYO3B,AGTR1,PSMA3,ATP1B3,ORC4,MACF1,JMJD1C,ST8SIA6,PPP1R2P1,WRN,TUBB6,FZD5,WEE2,GPN1,DDX60,AGPAT3,ERBB3,MCF2L,DTX2,KLC4,PPP1R11,FBXO25,MTCH1,ROR1,FASN,GNL2,RAD18,ARHGAP22,ERP44,SERAC1,EYA3,TET1,PPP2R5A,ADAMTS2,PPP2R2D,B3GNTL1,UBE2V2,MYO5A,EFTUD1,SBF1,GDA,EZH2,DCT,ABL2,DDX11,RNF19A,ZDHHC20,ARHGAP5,EHD3,FAF1,SDR42E1,TXNDC12,PHACTR1,FBXO22,RAB6A,KLHDC3,NLK,FOLH1B,RHOU,LIMK2,PPFIBP2,HERC2,CARD16,KLK2,CYP2R1,SCYL1,UBB,KATNAL2,NDUFB6,SERHL,UGT1A3,UGT1A5,UGT1A4,UGT1A8,UGT1A10,UGT1A9,UGT1A7,UGT1A6,EPHX4,AIM2,MAP2K3,MDM2,GUCY1A2,ADH1C,AOC2,INO80,PRKG1,RB1,C1S,DECR2,GLS,FEN1,TNKS2,EPHA1,PLG,SSR1,ATP2A2,GLT1D1,PELI2,PPIE,PROCA1,FUT8,ITIH1,NGLY1,CDC20,DST,GDI2,USP24,NMNAT1,PHOSPHO1,RACGAP1,PSMD2,PLBD2,VAV2,ECEL1,SBK2,PIP5K1A,DAP,EDN2,TBC1D26,ABCF2,NOSTRIN,FADS3,DIP2A,MAP2K6,SULT1A2,RANBP2,ARL8B,DUSP5,ULK2,ART3,ALG6,CAB39,GCAT,HMBS,ADCY10,NEK7,SUMO4,PDSS1,HSPA5,CHFR,AK3,VAC14,DNAH14,ARL2BP,PTPN12,RAB3GAP1,ADAM32,RAB43,METAP1,PARG,PRKDC,OAS3,DHRS7,SMUG1,TRMT12,USP47,PDE8B,PTPN9,C3,TNFAIP8,ARHGEF12,TRIM71,HECW1,SMG6,SYNGAP1,HABP2,ADCK5,ULK4,RASA2,ANP32E,PYROXD1,KARS,EPHB4,PKM,UMPS,RABGAP1L,ZP3,ALKBH3,CCDC111,SGK1,VHL,ARHGAP17,TBC1D15,TMED2,TUBA3C,GXYLT2,LAP3,IGF1R,ASH1L,PPP1R2,GCH1,BMPR2,PTPRU,CDC25C,PDCD6,MACROD2,SHOC2,ADCY9,HTT,TPTE,CYP3A5,HPR,CYP2F1,CYP27A1,MCCC1,UBA2,LRP5,ADPRHL1,TRPM2,RAB3A,EP400,ZCCHC9,HBS1L,TRADD,FARP1,NMNAT3,MLKL,PIGK,COQ7,PRMT7,MADD,RASA3,RGS6,MIB1,ATG7,PPCDC,SNCA,KDM2A,MANBA,ARHGAP31,NEIL2,LTA4H,ABCC3,CHD7,EXD3,GLUL,PRKCA,CPSF3L,DHFR,CSAD,COX6C,GCHFR,GNPNAT1,CAMKMT,PI4K2B,MPHOSPH10,PTPRJ,XRCC2,CLPB,DIP2C,PRKD2,HK1,ASAP1,ST6GALNAC4,NDUFB2,SYN3,CMAS,LACTB2,AGBL4,EHMT2,PLA2G4C,RGS7,GSK3B,CDC26,NRAS,RNASEH1,PAN3,GSTA2,ADCY6,USP32,DLG1,SRPK2,ABHD3,ACOT7,GMPS,MARCH7,UBR5,NUBP1,STK40,CRYM,KDM4C,RBPJ,SIRT2,UBE3C,ARHGAP23,PARN,SDHD,POLR3C,ITIH5,PTPRN2,PTPRG,RPS6KA2,LHPP,PRKACG,PDE3B,FBXL14,RASGRP3,B4GALT5,EPS8,NUAK1,UGT2B7,UGT2B15,FAHD1,HAGH,PRDX5,TRMT112,ACSF3,ADAM12,PAPOLA,ARHGAP27,RAB7A,GMFB,NCOA3,KIAA2018,MAPK6,SPTLC2,MTIF2,CDC27,EIF2B5,SEPHS1,COX6B1,RAD1,BRCA1,ARHGAP19,RAD51B,GNB3,PLEK,FNTA,KIDINS220,KIF16B,HYAL1,KIAA1967,PPP3CC,NF1,N6AMT2,AKR1B15,ADARB1,NXN,DHRS2,PTPRK,RAB3D,ACER2,TAOK3,MMP28,N4BP2,ATP5A1,KIF26B,PAPPA2,RIN3,CD101,ACER3,NUDT5,MSRA,LOXL2,TRIM39,PLCG2,USP28,PTGES,TRIM36,ACAA2,EGLN3,TRMT11,HIBCH,ZFYVE9,AMDHD2,GRK5,KDM4A,HERC4,COX6A2,CYP19A1,MTMR12,ALDH3B1,HS3ST5,CCNY,TOP2B,MAPKAP1,ARHGEF4,RFC1,THSD4,TPX2,DCLRE1C,ANAPC10,IQSEC1,STK3,GRK6,USP43,ILKAP,ITPR2,DNMBP,RRAGC,ARHGAP12,RGL1,APOBEC4,MINK1,CASC5,GTF2H4,PILRB,LYN,DDX43,DDX59,ATG10,AGPAT5,CHST6,AGAP11,BDH2,LATS2,TTK,CHST10,GSTM4,SERPINB6,NFX1,S1PR2,CPA5,NUDT3,PDE6A,SPAG9,CLN8,CBLB,PAK2,GALNT3,HSD3B2,ENTPD1,TBC1D16,IRAK4,BPNT1,MCEE,ARHGAP32,ENPEP,HADHB,DIS3L2,TBC1D1,KRIT1,PPT2,IMPA1,RGS9,PDE7A,KLK3,PTPN11,PSMD1,PPM1D,PRKRA,SLC27A4,TPP2,TTBK2,HDAC5,DYNC1H1,RPTOR,MTMR3,DIABLO,RGS8,TMPRSS11F,SULT1A1,IKBKB,CELA1,SMYD3,OPA1,KIF1B,SCARB1,RYR2,TTLL5,CALCR,OAZ3,PPP1R14D,PPIL3,PPP6R3,YES1,FASTKD2,SPACA3,CCL21,ATP6V0C,RSF1,MUC20,CTSS,MAP3K2,TEC,CA5A,MICAL3,HSPA2,UGGT2,LDHAL6A,GK5,AGBL2,RNF14,GLRX5,LIAS,SERTAD1 | 1.120045 | 1.05E-06 | 8.35E-04 |
| GO:0005488 | binding | molecular_function | STOX1,EFCAB7,TSEN2,RNF157,SHANK2,ZNF324,KIF23,CCT6A,RAD54L2,PYY,SSH2,ATAD3A,MYL6,ZNF845,METTL5,CUL7,PELP1,MCM10,LRP5L,ZBTB45,RNF7,PCBP2,MNAT1,CDK19,RNF215,SEPT7,TAF6,GRIN2A,PRIM2,MYO3B,AGTR1,PSMA3,ORC4,MACF1,JMJD1C,ANKMY1,PURG,WRN,TUBB6,HSPA1L,FZD5,PRH1,RPGRIP1,WEE2,GPN1,DDX60,EYS,CHMP3,ZNF107,ERBB3,ZFP64,MCF2L,KLHL17,DTX2,ATP5G2,ZNF211,PRPF3,ACTN1,ZNF605,WNT10A,SVIL,FBXO25,MTCH1,FANCC,ROR1,MTERFD2,SNED1,RBPJL,FASN,RBM39,BANP,GNL2,RAD18,ARHGAP22,TJP1,EYA3,S100A7A,TET1,PPP2R5A,ADAMTS2,KLF3,ZNF628,STAG1,UBE2V2,FBXL12,VPS54,MYO5A,SCUBE1,EFTUD1,SBF1,GDA,TRAK2,EZH2,RBMS1,DCT,ABL2,CUL2,DDX11,RNF19A,AFAP1,CALN1,ZDHHC20,ARHGAP5,HIST3H3,EHD3,CCDC88B,HIST3H2BB,HIST3H2A,CLUAP1,FAF1,SDR42E1,UBQLN4,PHACTR1,NUP210,SMARCC1,BRD1,RAB6A,KLHDC3,MICA,NLK,FOLH1B,RHOU,LIMK2,SNX19,PPFIBP2,FCAR,SRSF4,HERC2,MTUS2,ZHX3,NAPG,CYP2R1,HSPB6,TSNARE1,NRXN3,SCYL1,SNX14,MRPS31,UBB,GTF2A2,ZNF431,IGF2BP1,KATNAL2,RRP7A,CPEB3,PLXDC1,UGT1A3,UGT1A4,UGT1A8,UGT1A10,UGT1A9,UGT1A7,UGT1A6,AIM2,LYST,MAP2K3,TRIM43,CAPS2,TNRC6C,MAPT,MDM2,GUCY1A2,ADH1C,RPF1,FAM120A,SLC9A3,AOC2,STX8,FRMD4A,INO80,CTNNA1,CLIC6,PRKG1,SLC25A17,SH3GL1,RB1,TMEM181,NCAPD2,C1S,RPS15,DECR2,SP7,DISC1,GLS,FEN1,NR3C1,PEX19,RRN3,TNKS2,ARNTL2,IFI35,ZNF610,SYNE2,OSBPL9,A1CF,EPHA1,PLG,PCDH15,MAP4,ZNF607,ATP2A2,WNT5B,LDLRAP1,PABPC1,TRPS1,PELI2,PPIE,PROCA1,FUT8,ITIH1,NGLY1,CDC20,DST,MKL2,GDI2,EVA1C,ZNF888,NMNAT1,PHOSPHO1,RACGAP1,PSMD2,TRIM4,VAV2,ECEL1,SPIRE1,PDE4DIP,SBK2,PIP5K1A,DAP,EDN2,ABCF2,IL34,NOSTRIN,VPS52,FADS3,DIP2A,TIA1,ATCAY,ZNF808,EFCAB5,MAP2K6,TFAP2E,RANBP2,MYT1L,UVRAG,STON2,ARL8B,ULK2,ZNF701,HSPA6,GIMAP7,CXCL11,CAB39,GCAT,MBD5,ADCY10,GTF3C6,DLGAP4,E2F3,NEK7,PDSS1,HSPA5,CHFR,ZNF195,SNAI1,AK3,DNAH14,ANXA13,IFNA17,ARL2BP,AIRE,PTPN12,KLF2,RAB3GAP1,ADAM32,RAB43,HIST1H2BA,HIST1H2AA,OPTN,ZNF521,METAP1,PRKDC,FLYWCH1,OAS3,TPM4,HOOK3,DHRS7,ZNF433,ZNF485,SMUG1,MYRIP,TRIM26,USP47,ZNF28,PDE8B,PTPN9,ZNF354B,PABPC3,C3,SMAD2,DLG4,MESDC2,FAM188A,ARHGEF12,MT1F,ANO1,TRIM71,SMG6,SYNGAP1,HABP2,MRPL16,FANCD2,ULK4,RASA2,JMY,ZNF200,KARS,TERF2IP,SCMH1,EPHB4,PKM,MEIS3,CTTNBP2NL,RABGAP1L,ZP3,KANK1,NEDD9,ALKBH3,ZNF486,TNIP2,TMOD3,SGK1,GAS7,VHL,ARHGAP17,DYSF,MEX3A,SPTA1,TNFRSF10D,TBC1D15,TMED2,ZNF177,YWHAQ,TUBA3C,ZNF429,LAP3,HNRNPD,RBFOX3,IGF1R,ANTXR2,ASH1L,PPP1R13L,PPP1R2,GCH1,UBXN1,BMPR2,PTPRU,CDC25C,PDCD6,INVS,TMEM8B,ELF2,SHOC2,SNRPB,KCNT2,CDH5,ADCY9,HTT,ZNF611,POM121C,CYP3A5,HSPA13,NCOA6,HPR,ZNF716,CYP2F1,CYP27A1,RAB11FIP5,HTN1,DRG2,MCCC1,SUSD2,CDH18,UBA2,TAF4B,NIF3L1,LRP5,ADPRHL1,WIPI2,RAB3A,CRYBA1,TFDP2,CSF2RB,EP400,ZCCHC9,MTERFD3,ZNF876P,HBS1L,TRADD,FARP1,NMNAT3,SLFN12L,CCDC59,MLKL,PIGK,EPCAM,COQ7,PRMT7,MADD,RASA3,RGS6,CMIP,MIB1,MRPL21,ATG7,DUXA,SNCA,SH3RF3,ZNF257,KDM2A,CLEC12A,ENAH,ABLIM2,MANBA,ARHGAP31,ZNF91,NEIL2,LTA4H,FRMD3,ZNF747,ABCC3,CHD7,FOXP1,EXD3,GLUL,PRKCA,TNPO1,RPGRIP1L,CD3E,RELA,DHFR,CSAD,GCHFR,GTF3A,ANKRD42,ZNF587B,RNF145,GNPNAT1,PI4K2B,MPHOSPH10,HSPA14,CDNF,REEP1,PTPRJ,FERMT3,XRCC2,FLG,CLPB,DIP2C,VPS72,HP1BP3,PRKD2,HK1,ASAP1,COL27A1,IL37,SYN3,LACTB2,AGBL4,ZNF33A,EHMT2,PLA2G4C,SCIN,FAM101B,COBL,RGS7,GSK3B,CDC26,CSDE1,NRAS,RNASEH1,PAN3,ADCY6,ITGAM,USP32,SDCBP2,CACNA1S,DLG1,SRPK2,RMI1,ACOT7,ZNF83,GMPS,MARCH7,UBR5,ARPC3,NUBP1,STK40,CRYM,KDM4C,PAIP2B | 1.052845 | 1.12E-06 | 8.35E-04 |
| GO:0016787 | hydrolase activity | molecular_function | TSEN2,KIF23,RAD54L2,SSH2,ATAD3A,MYL6,CUL7,MNAT1,PTP4A1,MYO3B,AGTR1,PSMA3,ATP1B3,ORC4,MACF1,PPP1R2P1,WRN,TUBB6,GPN1,DDX60,MCF2L,KLC4,PPP1R11,MTCH1,FASN,GNL2,ARHGAP22,SERAC1,EYA3,PPP2R5A,ADAMTS2,PPP2R2D,MYO5A,EFTUD1,SBF1,GDA,EZH2,DDX11,ARHGAP5,EHD3,PHACTR1,RAB6A,KLHDC3,FOLH1B,RHOU,HERC2,CARD16,KLK2,KATNAL2,SERHL,EPHX4,AIM2,MDM2,INO80,PRKG1,C1S,GLS,FEN1,EPHA1,PLG,SSR1,ATP2A2,PROCA1,ITIH1,NGLY1,DST,GDI2,USP24,PHOSPHO1,RACGAP1,PLBD2,VAV2,ECEL1,PIP5K1A,DAP,TBC1D26,ABCF2,ARL8B,DUSP5,HSPA5,DNAH14,ARL2BP,PTPN12,RAB3GAP1,ADAM32,RAB43,METAP1,PARG,OAS3,SMUG1,USP47,PDE8B,PTPN9,C3,TNFAIP8,ARHGEF12,SMG6,SYNGAP1,HABP2,RASA2,ANP32E,RABGAP1L,ALKBH3,ARHGAP17,TBC1D15,TMED2,TUBA3C,LAP3,PPP1R2,GCH1,PTPRU,CDC25C,PDCD6,MACROD2,SHOC2,ADCY9,HTT,TPTE,ADPRHL1,TRPM2,RAB3A,EP400,ZCCHC9,HBS1L,TRADD,FARP1,PIGK,MADD,RASA3,RGS6,SNCA,MANBA,ARHGAP31,NEIL2,LTA4H,ABCC3,CHD7,EXD3,PRKCA,CPSF3L,GCHFR,MPHOSPH10,PTPRJ,XRCC2,CLPB,PRKD2,ASAP1,LACTB2,AGBL4,PLA2G4C,RGS7,GSK3B,NRAS,RNASEH1,ADCY6,USP32,DLG1,ABHD3,ACOT7,GMPS,NUBP1,SIRT2,ARHGAP23,PARN,ITIH5,PTPRN2,PTPRG,LHPP,PRKACG,PDE3B,RASGRP3,NUAK1,FAHD1,HAGH,PRDX5,ADAM12,ARHGAP27,RAB7A,KIAA2018,MTIF2,EIF2B5,RAD1,ARHGAP19,RAD51B,GNB3,PLEK,FNTA,KIF16B,HYAL1,PPP3CC,NF1,ADARB1,PTPRK,RAB3D,ACER2,MMP28,N4BP2,ATP5A1,KIF26B,PAPPA2,RIN3,CD101,ACER3,NUDT5,PLCG2,USP28,EGLN3,HIBCH,ZFYVE9,AMDHD2,MTMR12,TOP2B,ARHGEF4,RFC1,THSD4,DCLRE1C,IQSEC1,USP43,ILKAP,ITPR2,DNMBP,RRAGC,ARHGAP12,RGL1,APOBEC4,MINK1,CASC5,GTF2H4,DDX43,DDX59,AGAP11,SERPINB6,CPA5,NUDT3,PDE6A,PAK2,ENTPD1,TBC1D16,BPNT1,ARHGAP32,ENPEP,DIS3L2,TBC1D1,KRIT1,PPT2,IMPA1,RGS9,PDE7A,KLK3,PTPN11,PPM1D,TPP2,HDAC5,DYNC1H1,MTMR3,DIABLO,RGS8,TMPRSS11F,CELA1,OPA1,KIF1B,RYR2,CALCR,PPP1R14D,PPP6R3,SPACA3,CCL21,ATP6V0C,RSF1,CTSS,AGBL2,CGNL1,USP34,ECT2L,FLT1,CGN,NT5C2,DNAH3,ATP2B4,STARD9,RAB3GAP2,REV3L,ACAD10,PLCL2,MUL1,INPP5K,PPP5C,CMSS1,FRS2,MSH4,RAPGEF6,MYH10,PTEN,RNASEL,RPP21,WFS1,ANXA3,SRGAP1,PPP2R1A,NUDT15,NLRP12,MAN2A2,OBSCN,GTF2H1,DOCK3,DOCK6,TATDN2,CNOT1,PRSS48,BTD,CSNK2A1,KALRN,DFFB,ST5,EGFR,USP8,CTGF,DDX24,RIMBP2,NUDT6,DNASE2,RANBP10,KLC1,MASP2,TRHDE,ACOT11,LMLN,MTHFD1L,IQGAP1,CHRM5,TBC1D8,LPIN1,USP37,USP14,ATP11B,RASL10B,DENND3,VAV1,SERPINB4,PTPRO,SMAP1,RALGDS,TPM1,GBA2,PTTG1,GLS2,PCSK6,GNA12,ELANE,C5,ARHGAP42,DPP9,EPHA5,GBP1,IMMP1L,RNASET2,CCL3,SSH1,MYH3,MGLL,DNAJC6,LONRF2,ACOT1,PFKFB3,ADAM33,MEN1,F2R,PGAM2,TBC1D2B,PTPN14,ATP8A2,CES1,ARSB,WFDC3,BPTF,SET,CEP192,RHBDL2,RBM26,ASNS,ERAP2,DLC1,ADAMTS9,ACSBG2,LMTK2,CELA3A,FGFR2,LGMN,CCDC155,DNAH17,TPSD1,RTEL1-TNFRSF6B,DDX31,ARHGEF37,GNAT2,NGEF,PM20D1,ATP10A,AICDA,SMARCAD1,CD27,MYO5B,THOP1,EXOSC9,ARHGAP10,ABCA13,SGSM3,AARS,INPP5B,SH3BP4,ABR,MYO1B,PPP4R2,MYH11,KIF25,SPG7,ADAM23,EYA4,DUSP22,ARHGEF10,ACE,MEF2A,SH2D4A,KIF19,PTPRM,SETX,PTPN1,DUSP3,ATP11A,MMP21,ADAMTS12,APLF,TG,PDIA6,CHP2,DOCK1,CAT,PSMB7,MYH15,COL28A1,PXDNL,CPVL,CNOT2,NSF,PLCXD2,RFC2,HDAC4,FBP1,ADAM21,PSD4,PEPD,CNST,PHACTR3,CYFIP2,PITRM1,SCPEP1,GNAO1,FBXO8,ARHGAP18,METAP1D,PTPRE,ROBO1,TRABD2B,AEBP1,PLA2G4A,ASCC3,PRKCE,PDE9A,SFI1,ITPR1,ATL1,ATP10D,ABCC4,PPP2R2C,ADAP1,UBASH3B,ARHGAP30,ATP13A5,COL6A3,ATP6V1D | 1.197018 | 1.47E-06 | 9.46E-04 |
| GO:0005829 | cytosol | cellular_component | KIF23,CCT6A,KRTAP6-1,RCC2,MYL6,CUL7,PCBP2,PSMA3,CHMP3,MCF2L,ACTN1,FANCC,FASN,ARHGAP22,TJP1,GDA,DCT,ABL2,CUL2,ARHGAP5,FAF1,UBQLN4,PHACTR1,RAB6A,KLHDC3,RHOU,UBB,IGF2BP1,AIM2,LYST,MAP2K3,TNRC6C,MAPT,MDM2,ADH1C,CTNNA1,PRKG1,RPS15,GLS,NR3C1,PEX19,RSU1,IFI35,LDLRAP1,PABPC1,PELI2,CDC20,GDI2,PHOSPHO1,RACGAP1,PSMD2,HSF2BP,VAV2,PIP5K1A,SEC16A,MAP2K6,SULT1A2,RANBP2,CAB39,HMBS,ADCY10,PTPN12,PARG,OAS3,TPM4,PDE8B,SMAD2,ARHGEF12,SMG6,KARS,PKM,UMPS,TNIP2,VHL,ARHGAP17,SPTA1,HNRNPD,GCH1,CDC25C,SNRPB,HTT,WIPI2,RAB3A,TRADD,FARP1,NMNAT3,PRMT7,PPCDC,SNCA,ENAH,ARHGAP31,LTA4H,GLUL,PRKCA,TNPO1,RELA,DHFR,GCHFR,GNPNAT1,PI4K2B,HSPA14,HK1,IL37,AGBL4,PLA2G4C,RPL32,RGS7,GSK3B,CDC26,PAN3,GSTA2,DLG1,ACOT7,GMPS,NUBP1,PARN,CLIP1,SH3BGR,PACSIN2,IGF2BP2,KRT14,RPS6KA2,LHPP,ELMO1,PRKACG,PDE3B,OSBPL5,BRPF3,FAHD1,PRDX5,CDC27,VCL,EIF2B5,ARHGAP19,RPL3,PLEK,FNTA,KIDINS220,PPP3CC,KPNA1,NXN,NBEA,TUBGCP5,N4BP2,TRIM39,PLCG2,EGLN3,PDCD6IP,CASC3,HERC4,TOP2B,MAPKAP1,STAT5B,ARHGEF4,ANAPC10,STK3,ARHGAP12,RGL1,CASC5,SMAD1,NPHP4,LYN,OSBPL3,LATS2,PLEC,SERPINB6,PDE6A,SPAG9,CBLB,ABI2,PAK2,IRAK4,BPNT1,ARHGAP32,ACTG1,PDE7A,C9orf89,PTPN11,PSMD1,TPP2,TTBK2,HDAC5,DYNC1H1,RPTOR,MTMR3,DIABLO,BNIPL,SULT1A1,IKBKB,OAZ3,YES1,TEC,LIMS1,SH3D19,PUM1,AGBL2,PARD3,RPL9,BUB1,RNF135,NT5C2,CHUK,INPP5K,PPP5C,PTEN,RNASEL,FLNB,SRGAP1,PPP2R1A,AHSP,OTOF,OBSCN,DOCK6,WWP1,PFAS,CNOT1,AKR7A3,SNX27,CSNK2A1,KALRN,NUP107,DFFB,USP8,CTGF,MAD1L1,NCF2,RANBP9,RANBP10,KLC1,RPL26L1,TNRC6A,MTHFD1L,PIK3C2G,STAM,BCAT1,ERC1,ITCH,LPIN1,APBB1IP,WWOX,VAV1,CRYL1,KNTC1,RALGDS,NUP160,TPM1,CCDC88A,PTTG1,FDPS,NFATC1,AK2,AAGAB,SLC7A5,DPP9,STK24,GBP1,HLCS,YWHAG,CCL3,CTBP1,MYH3,IL26,DNAJC6,ACOT1,PFKFB3,MEN1,FMN2,HIF3A,PGAM2,FOXO1,NBR1,AK8,SET,CEP192,ASNS,EGLN1,NAPRT1,DLC1,IRF1,MKNK1,NCALD,GRB10,NGEF,PACS1,S100A13,NDRG4,EXOSC9,ARHGAP10,AARS,INPP5B,MAPK1,RPS6KB1,DAB1,ABR,PRTFDC1,PSMD5,MYH11,AP2A2,VPS28,PFKP,PARD6G,ARHGEF10,EEF1E1,RBP2,PRKCH,PFKL,PTPN1,DUSP3,EIF2AK4,EDC3,DBNL,APLF,DOCK1,DPYD,CAT,MYBPC1,ANO7,PSMB7,GSTT1,CNOT2,NSF,GAB1,HDAC4,FBP1,URM1,PRKAG2,S100A16,SCPEP1,CNOT3,ARHGAP18,CAPZB,UPP2,AMD1,PLA2G4A,COG5,PRKCE,PDE9A,SFI1,IL22RA2,AKR1B1,ANK1,NOS1,PGM2L1,ARHGAP30,ATP6V1D | 1.253819 | 1.48E-06 | 9.46E-04 |
| GO:0030054 | cell junction | cellular_component | SHANK2,GRIN2A,KLHL17,ACTN1,SVIL,TJP1,STAG1,AFAP1,PHACTR1,RHOU,PLXDC1,FRMD4A,CTNNA1,SH3GL1,DISC1,SYNE2,DST,ATCAY,PTPN12,DLG4,NEDD9,ARHGAP17,ASH1L,PTPRU,CDH5,EPCAM,SNCA,ENAH,ARHGAP31,RPGRIP1L,PTPRJ,FERMT3,SYN3,DLG1,CHRNG,GJB4,CLSTN1,VCL,BRCA1,GABRB1,PTPRK,TNS3,GJC1,ITGB5,FRMPD2,IQSEC1,DNMBP,MINK1,NPHP4,PLEC,FAT2,ABI2,GABRG3,ARHGAP32,CRIPT,KRIT1,EPB41L5,LIMD1,CACNG8,LIMS1,PARD3,CGNL1,GRIA1,CGN,OLFM2,TTBK1,FLNB,AJAP1,OTOF,CTBP2,RIMBP2,LMLN,IQGAP1,CHRM5,SEPT11,SCN1A,APBB1IP,VAMP3,FAT1,SLC17A7,DLG5,TMEM163,LRFN1,ZNRF1,PANX2,JAM2,GABRB3,WNK4,DLC1,NFIA,PLEKHA7,FNDC1,UTRN,SPECC1L,PKP1,GRIN3B,PODXL,S100A14,CLDN6,SH3PXD2B,AMPH,CCDC85C,CADM1,SGSM3,LIG4,TLN2,MAPK1,RPS6KB1,PARD6G,COL13A1,PTPRM,GPHN,HMCN1,CLCA2,SHISA9,ANO7,XIRP2,SMAGP,PSD4,PALLD,TANC1,CYFIP2,TEK,FBXO8,GRID1,P2RX6,SH3PXD2A,P2RX7,GSG1L,SLC8A1 | 1.486183 | 1.82E-06 | 1.09E-03 |

### Supplementary Table S7-a.

**The top 15 significant signatures related to patients with schizophrenia in the KEGG pathway enrichment analysis of differentially methylated probes.**

| Pathway ID | Pathway name | Pathway class | Symbols in list | Fold enrichment | *P* value | FDR |
| --- | --- | --- | --- | --- | --- | --- |
| hsa00040 | Pentose and glucuronate interconversions | Carbohydrate metabolism | UGT2B17,AKR1B1,UGT2B15 | 3.64910249 | 0.04741887 | 0.9153466 |
| hsa04144 | Endocytosis | Transport and catabolism | EHD3,SH3GL1,PIP5K1A,FOLR3,EEA1,ASAP1,PSD3,ITCH,ERBB4 | 1.797319137 | 0.05411106 | 0.9153466 |
| hsa04150 | mTOR signaling pathway | Signal transduction | STRADA,RPTOR,RPS6KA2,RPS6KB1 | 2.676008493 | 0.06012207 | 0.9153466 |
| hsa04012 | ErbB signaling pathway | Signal transduction | ABL2,NRAS,ERBB4,PAK4,RPS6KB1 | 2.306903873 | 0.06243264 | 0.9153466 |
| hsa04020 | Calcium signaling pathway | Signal transduction | AGTR1,ATP2B4,ERBB4,SLC8A1,RYR3,ADRA1B,GNAL,P2RX6 | 1.774149277 | 0.07238588 | 0.9153466 |
| hsa04666 | Fc gamma R-mediated phagocytosis | Immune system | PIP5K1A,ASAP1,DOCK2,RPS6KB1,PLA2G4A | 2.205501505 | 0.07264604 | 0.9153466 |
| hsa00590 | Arachidonic acid metabolism | Lipid metabolism | CYP4F11,GPX3,ALOX15B,PLA2G4A | 2.432734993 | 0.0792996 | 0.9153466 |
| hsa05211 | Renal cell carcinoma | Cancers: Specific types | NRAS,EGLN1,CUL2,PAK4 | 2.432734993 | 0.0792996 | 0.9153466 |
| hsa04010 | MAPK signaling pathway | Signal transduction | MAP2K6,DUSP5,MAP2K3,NRAS,NLK,NF1,MAP3K8,RPS6KA2,DUSP3,PLA2G4A | 1.574122643 | 0.08535486 | 0.9153466 |
| hsa04664 | Fc epsilon RI signaling pathway | Immune system | MAP2K6,MAP2K3,NRAS,PLA2G4A | 2.293721565 | 0.09355703 | 0.9153466 |
| hsa01040 | Biosynthesis of unsaturated fatty acids | Lipid metabolism | TECR,SCD5 | 3.822869275 | 0.09465914 | 0.9153466 |
| hsa04520 | Adherens junction | Cell communication | NLK,TJP1,TCF7L1,SORBS1 | 2.199459035 | 0.10498024 | 0.9153466 |
| hsa00062 | Fatty acid elongation | Lipid metabolism | HADHB,TECR | 3.49044586 | 0.1104296 | 0.9153466 |
| hsa04151 | PI3K-Akt signaling pathway | Signal transduction | SGK1,NRAS,PPP2R2C,RPTOR,GNG7,IL4R,VWF,PPP2R1A,RPS6KB1,ITGA1,COL5A1,TEK | 1.404319326 | 0.11226083 | 0.9153466 |

### Supplementary Table S7-b.

**The top 15 significant signatures related to psychosis risk syndrome in the KEGG pathway enrichment analysis of differentially methylated probes.**

| Pathway ID | Pathway name | Pathway class | Symbols in list | Fold enrichment | *P* value | FDR |
| --- | --- | --- | --- | --- | --- | --- |
| hsa00983 | Drug metabolism - other enzymes | Xenobiotics biodegradation and metabolism | UGT1A3,UGT1A5,UGT1A4,UGT1A8,UGT1A10,UGT1A9,UGT1A7,UGT1A6,UMPS,CYP3A5,GMPS,UGT2B7,UGT2B15,CES1,DPYD,UPP2,UGT2B17 | 2.885531136 | 2.96E-05 | 0.0037392 |
| hsa00040 | Pentose and glucuronate interconversions | Carbohydrate metabolism | UGT1A3,UGT1A5,UGT1A4,UGT1A8,UGT1A10,UGT1A9,UGT1A7,UGT1A6,UGT2B7,UGT2B15,CRYL1,UGT2B17,AKR1B1 | 3.41017316 | 3.57E-05 | 0.0037392 |
| hsa04144 | Endocytosis | Transport and catabolism | HSPA1L,RNF103-CHMP3,ERBB3,EHD3,MDM2,SH3GL1,LDLRAP1,PIP5K1A,HSPA6,SMAD2,IGF1R,RAB11FIP5,ASAP1,RAB7A,PDCD6IP,ZFYVE9,GRK5,IQSEC1,GRK6,CBLB,HSPA2,PARD3,RAB11FIP2,FLT1,WWP1,EGFR,USP8,RAB11FIP3,STAM,ITCH,SMAP1,IL2RA,DNAJC6,F2R,FGFR2,HLA-C,RUFY1,AP2A2,VPS28,PARD6G,MVB12B,PSD4 | 1.808840413 | 4.38E-05 | 0.0037392 |
| hsa00053 | Ascorbate and aldarate metabolism | Carbohydrate metabolism | UGT1A3,UGT1A5,UGT1A4,UGT1A8,UGT1A10,UGT1A9,UGT1A7,UGT1A6,UGT2B7,UGT2B15,UGT2B17 | 3.526760277 | 1.01E-04 | 0.0064418 |
| hsa04520 | Adherens junction | Cell communication | ACTN1,TJP1,NLK,CTNNA1,SNAI1,SMAD2,IGF1R,PTPRJ,VCL,ACTG1,WASF2,YES1,PARD3,CSNK2A1,EGFR,IQGAP1,MAPK1,PTPRM,PTPN1 | 2.253085955 | 4.09E-04 | 0.0201276 |
| hsa04724 | Glutamatergic synapse | Nervous system | SHANK2,GRIN2A,GLS,DLG4,SLC38A1,ADCY9,GLUL,PRKCA,PLA2G4C,ADCY6,CACNA1C,PRKACG,GNB4,GNB3,PPP3CC,ITPR2,GRIA1,SLC17A7,GLS2,GRIN3B,MAPK1,CHP2,GNG7,GNAO1,PLA2G4A,ITPR1 | 1.940270936 | 4.72E-04 | 0.0201276 |
| hsa04730 | Long-term depression | Nervous system | GUCY1A2,PRKG1,IGF1R,PRKCA,PLA2G4C,NRAS,ITPR2,LYN,GRIA1,PPP2R1A,GNA12,MAPK1,GNAO1,PLA2G4A,ITPR1,NOS1 | 2.308424908 | 8.79E-04 | 0.0321351 |
| hsa00980 | Metabolism of xenobiotics by cytochrome P450 | Xenobiotics biodegradation and metabolism | UGT1A3,UGT1A5,UGT1A4,UGT1A8,UGT1A10,UGT1A9,UGT1A7,UGT1A6,ADH1C,CYP3A5,CYP2F1,GSTA2,UGT2B7,UGT2B15,ALDH3B1,GSTM4,AKR7A3,GSTT1,UGT2B17 | 2.081965503 | 1.16E-03 | 0.0369856 |
| hsa05204 | Chemical carcinogenesis | Cancers: Overview | UGT1A3,UGT1A5,UGT1A4,UGT1A8,UGT1A10,UGT1A9,UGT1A7,UGT1A6,ADH1C,SULT1A2,CYP3A5,GSTA2,UGT2B7,UGT2B15,ALDH3B1,GSTM4,SULT1A1,GSTT1,UGT2B17 | 2.055940934 | 1.35E-03 | 0.0385422 |
| hsa04727 | GABAergic synapse | Nervous system | TRAK2,GLS,SLC38A1,ADCY9,GLUL,PRKCA,ADCY6,CACNA1S,CACNA1C,PRKACG,GNB4,GNB3,GABRB1,GABRG3,GLS2,GABRB3,GPHN,NSF,GNG7,GNAO1 | 1.967407592 | 1.80E-03 | 0.0459974 |
| hsa00860 | Porphyrin and chlorophyll metabolism | Metabolism of cofactors and vitamins | UGT1A3,UGT1A5,UGT1A4,UGT1A8,UGT1A10,UGT1A9,UGT1A7,UGT1A6,HMBS,UGT2B7,UGT2B15,UGT2B17 | 2.415793509 | 2.55E-03 | 0.0548998 |
| hsa00982 | Drug metabolism - cytochrome P450 | Xenobiotics biodegradation and metabolism | UGT1A3,UGT1A5,UGT1A4,UGT1A8,UGT1A10,UGT1A9,UGT1A7,UGT1A6,ADH1C,CYP3A5,GSTA2,UGT2B7,UGT2B15,ALDH3B1,GSTM4,GSTT1,UGT2B17 | 2.043917888 | 2.57E-03 | 0.0548998 |
| hsa00140 | Steroid hormone biosynthesis | Lipid metabolism | UGT1A3,UGT1A5,UGT1A4,UGT1A8,UGT1A10,UGT1A9,UGT1A7,UGT1A6,CYP3A5,UGT2B7,UGT2B15,CYP19A1,HSD3B2,UGT2B17 | 2.203496503 | 2.93E-03 | 0.0577845 |
| hsa04010 | MAPK signaling pathway | Signal transduction | HSPA1L,NLK,MAP2K3,MAPT,MAP2K6,DUSP5,HSPA6,RASA2,PRKCA,RELA,PLA2G4C,NRAS,CACNA1S,CACNA1C,RPS6KA2,PRKACG,RASGRP3,PPP3CC,NF1,FGF12,TAOK3,STK3,CD14,PAK2,IKBKB,CACNG8,MAP3K2,HSPA2,CHUK,PPP5C,FLNB,EGFR,GNA12,MECOM,MKNK1,FGFR2,MAPK1,DUSP22,DUSP3,CHP2,ATF2,PLA2G4A,CACNG3 | 1.45973928 | 3.47E-03 | 0.0635142 |
| hsa04114 | Oocyte meiosis | Cell growth and death | PPP2R5A,CDC20,YWHAQ,IGF1R,CDC25C,ADCY9,CDC26,ADCY6,RPS6KA2,PRKACG,CDC27,PPP3CC,ANAPC10,ITPR2,BUB1,PPP2R1A,PTTG1,YWHAG,SLK,MAPK1,CHP2,ITPR1 | 1.731318681 | 5.64E-03 | 0.0921213 |

### Supplementary Table S8.

**Methylation specific Haplotype analysis of *SYNGAP1* and *L3MBTL4* genes.**

| Target Genes | Haplotype | Healthy control  Mean(S.D.)  (n=10) | Psychosis risk syndrome  Mean(S.D.)  (N’s=41) | *p*-value | First-episode schizophrenia  Mean(S.D.)  (N’s=39) | *p*-value |
| --- | --- | --- | --- | --- | --- | --- |
| *L3MBTL4* | ttt | 0.24(0.06) | 0.29(0.13) | >0.05 | 0.32(0.13)** | 0.006 |
|  | ttc | 0.13(0.03) | 0.11(0.04) | >0.05 | 0.11(0.04)* | 0.04 |
|  | ctt | 0.08(0.01) | 0.08(0.06) | >0.05 | 0.10(0.05)* | 0.03 |
|  | ctc | 0.08(0.01) | 0.07(0.03)* | 0.04 | 0.08(0.04) | >0.05 |
|  | cct | 0.07(0.01) | 0.07(0.04) | >0.05 | 0.07(0.03) | >0.05 |
| *SYNGAP1* | c | 0.85(0.03) | 0.80(0.06)* | 0.01 | 0.79(0.06)*** | 0.0009 |
|  | t | 0.15(0.03) | 0.20(0.06)* | 0.01 | 0.21(0.06)*** | 0.0009 |

Statistical significance of haplotypes of *SYNGAP1* and *L3MBTL4* was defined from t-test between the two groups. S.D.=Standard Deviation. *** *p*<.001, ***p*<0.01, **p*<0.05.

### Supplementary Table S9-a.

**Effects Of Being On Antipsychotic Medication The Methylation Values In The 18 Selected Genes In Subjects With First-Episode Schizophrenia (Discovery Cohort)**

| Gene | N | No Antipsychotics  (Mean± S.D.) | N | On Antipsychotics  (Mean ± S.D.) | Test (T, DF, *p*) |
| --- | --- | --- | --- | --- | --- |
| *AGTR1*-cg10297223 | 11 | 0.5401908(0.0524148) | 29 | 0.5075359(0.0541755) | 1.717, 38, 0.094 |
| *DISC1*-cg25215021 | 11 | 0.6945067(0.0228670) | 29 | 0.7038906(0.0320903) | -0.885, 38, 0.382 |
| *SHANK2*-cg18440889 | 11 | 0.7280985(0.0491298) | 29 | 0.7098626(0.0451256) | 1.114, 38, 0.272 |
| *SYNGAP1*-cg14079545 | 11 | 0.4374861(0.0718348) | 29 | 0.4560020(0.0505144) | -0.919, 38, 0.364 |
| *L3MBTL4*-cg00451011 | 11 | 0.4529138(0.1051795) | 29 | 0.4423988(0.1408960) | 0.224, 38, 0.824 |
| *BRD1*-cg19984503 | 11 | 0.7570759(0.0376902) | 29 | 0.7528558(0.0375817) | 0.317, 38, 0.753 |
| *CPEB3*-cg10967735 | 11 | 0.5845885(0.0093450) | 29 | 0.6010418(0.0289923)* | -2.708, 37.551, 0.010 |
| *EMC2*-cg13442241 | 11 | 0.7392528(0.0577374) | 29 | 0.6982498(0.0530825)* | 2.131, 38, 0.040 |
| *NAPG*-cg22389991 | 11 | 0.6640542(0.0338285) | 29 | 0.6710013(0.0342029) | -0.575, 38, 0.569 |
| *NR3C1*-cg19645279 | 11 | 0.5827970(0.0325560) | 29 | 0.5887860(0.0368159) | -0.473, 38, 0.639 |
| *CALN1*-cg04158140 | 11 | 0.5865799(0.0232832) | 29 | 0.6109054(0.0424207) | -1.793, 38, 0.081 |
| *ROR1*-cg19963700 | 11 | 0.8051075(0.0623849) | 29 | 0.7812180(0.0527840) | 1.216, 38, 0.231 |
| *JMJD1C*-cg17983571 | 11 | 0.7814971(0.0976078) | 29 | 0.7296032(0.0802369) | 1.721, 38, 0.093 |
| *NBEA*-cg00500007 | 11 | 0.53591846(0.0755927) | 29 | 0.4907832(0.0634260) | 1.907, 38, 0.064 |
| *DLG1*-cg07068730 | 11 | 0.6554056(0.0590820) | 29 | 0.6598571(0.0434715) | -0.261, 38, 0.795 |
| *DAP*-cg03278573 | 11 | 0.8033684(0.0817259) | 29 | 0.7646056(0.0843495) | 1.308, 38, 0.199 |
| *RGMA*-cg17350690 | 11 | 0.6990340(0.0547490) | 29 | 0.7218138(0.0845300) | -0.827, 38, 0.414 |
| *PTPRN2*-cg21705926 | 11 | 0.5621491(0.0931914) | 29 | 0.6476326(0.0982104)* | -2.491, 38, 0.017 |

Statistical significance of individual component of each group was defined upon t-test between the two groups. S.D.=Standard Deviation. *p<0.05.

### Supplementary Table S9-b.

**Effects Of Being On Antipsychotic Medication The Methylation Values In The 18 Selected Genes In Subjects With First-Episode Schizophrenia (Validation Cohort)**

| Gene | N | No Antipsychotics  (Mean± S.D.) | N | On Antipsychotics  (Mean ± S.D.) | Test (T, DF, *p*) |
| --- | --- | --- | --- | --- | --- |
| *AGTR1*-cg10297223 | 21 | 0.9356153(0.0199868) | 18 | 0.9195643(0.0278379)* | 2.089, 37, 0.044 |
| *DISC1*-cg25215021 | 21 | 0.9515966(0.0117766) | 18 | 0.9530880(0.0105109) | -0.414, 37, 0.681 |
| *SHANK2*-cg18440889 | 21 | 0.8584819(0.0173874) | 18 | 0.8557992(0.0170726) | 0.484, 37, 0.631 |
| *SYNGAP1*-cg14079545 | 21 | 0.7829755(0.0704661) | 18 | 0.7888651(0.0489944) | -0.298, 37, 0.767 |
| *L3MBTL4*-cg00451011 | 21 | 0.4078158(0.1701942) | 18 | 0.3190115(0.1490913) | 1.719, 37, 0.094 |
| *BRD1*-cg19984503 | 21 | 0.9619640(0.0036785) | 18 | 0.9617239(0.0039505) | 0.196, 37, 0.845 |
| *CPEB3*-cg10967735 | 21 | 0.6412960(0.0215659) | 18 | 0.6482647(0.0187063) | -1.069, 37, 0.292 |
| *EMC2*-cg13442241 | 21 | 0.9721471(0.0037875) | 18 | 0.9718600(0.0022595) | 0.281, 37, 0.780 |
| *NAPG*-cg22389991 | 21 | 0.9777588(0.0029966) | 18 | 0.9764257(0.0034483) | 1.292, 37, 0.204 |
| *NR3C1*-cg19645279 | 21 | 0.9666773(0.0075158) | 18 | 0.9680004(0.0064966) | -0.583, 37, 0.563 |
| *CALN1*-cg04158140 | 21 | 0.9600280(0.0047964) | 18 | 0.9573774(0.0071712) | 1.374, 37, 0.178 |
| *ROR1*-cg19963700 | 21 | 0.9017400(0.0157417) | 18 | 0.8981340(0.0193376) | 0.642, 37, 0.525 |
| *JMJD1C*-cg17983571 | 21 | 0.9740779(0.0127229) | 18 | 0.9779701(0.0016798) | -1.286, 37, 0.206 |
| *NBEA*-cg00500007 | 21 | 0.7090063(0.0428052) | 18 | 0.6832885(0.0524394) | 1.686, 37, 0.100 |
| *DLG1*-cg07068730 | 21 | 0.9831544(0.0025768) | 18 | 0.9829421(0.0034697) | 0.219, 37, 0.828 |
| *DAP*-cg03278573 | 21 | 0.9302413(0.0115777) | 18 | 0.9321618(0.0104163) | -0.541, 37, 0.592 |
| *RGMA*-cg17350690 | 21 | 0.6430159(0.0376896) | 18 | 0.6368226(0.0243855) | 0.598, 37, 0.554 |
| *PTPRN2*-cg21705926 | 21 | 0.6266264(0.0857246) | 18 | 0.5918817(0.0622245) | 1.426, 37, 0.162 |

Statistical significance of individual component of each group was defined upon t-test between the two groups. S.D.=Standard Deviation. *p<0.05.
